# Genome-wide association analysis and replication in 810,625 individuals identifies novel therapeutic targets for varicose veins

**DOI:** 10.1101/2020.05.14.095653

**Authors:** Waheed-Ul-Rahman Ahmed, Akira Wiberg, Michael Ng, Wei Wang, Adam Auton, 23andMe Research Team, Regent Lee, Ashok Handa, Krina T Zondervan, Dominic Furniss

**Author notes:** **Correspondence:** Professor Dominic Furniss, Nuffield Department of Orthopaedics, Rheumatology and Musculoskeletal Science, University of Oxford, Botnar Research Centre, Windmill Road, Oxford, OX3 7LD, United Kingdom. **Subject Terms:** Genetic, Association Studies; Genetics; Vascular disease.

## Abstract

**Background:** Varicose veins (VVs) affect one-third of Western society, with a significant subset of patients developing venous ulceration, and ongoing management of venous leg ulcers costing around $14.9 billion annually in the USA. There is no current medical management for VVs, with approaches limited to compression stockings, ablation techniques, or open surgery for more advanced disease. A significant proportion of patients report a positive family history, and heritability is ~17%, suggesting a strong genetic component. We aimed to identify novel therapeutic targets by improving our understanding of the aetiopathology and genetic architecture of VVs.

**Methods:** We performed the largest two-stage genome-wide association study of VVs in 401,656 subjects from UK Biobank, and replication in 408,969 subjects from 23andMe (total 135,514 varicose veins cases and 675,111 controls). We constructed a genetic risk score for VVs to investigate its use as a prognostic tool. Genes and pathways were prioritised using a suite of bioinformatic tools, and therapeutic targets identified using the Open Targets Platform.

**Results:** We discovered 49 signals at 46 susceptibility loci associated with VVs, including 29 previously unreported genetic associations (28 susceptibility loci). We demonstrated that patients with VVs requiring surgery have a higher genetic risk score than those managed non-surgically. We map 237 genes to these loci, many of which are biologically relevant and tractable to therapeutic targeting or repurposing (notably *VEGFA*, *COL27A1*, *EFEMP1*, *PPP3R1* and *NFATC2*). Tissue enrichment analyses implicated vascular tissue, and several genes were enriched in biological pathways relating to extracellular matrix biology, inflammation, angiogenesis, lymphangiogenesis, vascular smooth muscle cell migration, and apoptosis.

**Conclusions:** Genes and pathways identified represent biologically plausible contributors to the pathobiology of VVs, identifying promising candidates for further investigation of venous biology and potential therapeutic targets. We have provided the proof-of-principle that genetic risk score correlates with disease severity, which represents a first step in personalised medicine approaches to varicose veins.

## Introduction

Varicose veins (VVs) are a very common manifestation of chronic venous disease, affecting over 30% of the population in Western countries.^1^ In the USA, chronic venous disease affects over 11 million males and 22 million females aged 40-80 years^2^, meaning it is twice as prevalent as coronary heart disease.^3^ Chronic venous insufficiency leads to serious complications in 10% of cases, including lipodermatosclerosis, venous ulceration, and rarely amputation. Despite best care, 25-50% of venous leg ulcers remain unhealed after 6 months of treatment.^4^ Ongoing management of venous leg ulcers costs around $14.9 billion annually and 4.5 million work days per year are lost to venous-related illness in the USA.^5,6^ Despite this, at present there are no medical treatments for VVs. For symptomatic patients, endovenous ablation is the first-line treatment approach.^7^ However, recurrence following surgery is 20%, with no difference in rate compared to conventional open surgery.^8^

VVs are thought to develop from a combination of valvular insufficiency, venous wall alterations, and haemodynamic changes that precipitate venous reflux, stasis, and hypertension of the venous network, causing varicosities.^9,10^ Risk factors for VVs include older age, female sex, pregnancy, a positive family history, obesity, tall height, and previous DVT.^3,11,12^ Many patients with VVs report a positive family history^13^, and amongst offspring with one affected parent the familial standardised incidence ratio is 2.39^14^, with a heritability of 17%^15^, suggesting a genetic component to aetiology. Two recent genome-wide association studies (GWAS) of VVs have been described. Ellinghaus *et al*.^16^ tested for associations in 323 cases and 4,619 controls, with suggestive associations examined in an independent cohort totalling 1,946 cases and 3,146 controls. They reported two associations, mapped to *EFEMP1* and *KCNH8*. Fukaya *et al*.^17^ used UK Biobank to identify a further 30 putative associations (27 loci) associated with VVs. However, their cases were defined only by the International Classification of Diseases (ICD) diagnostic codes, meaning that thousands of cases defined by operative intervention codes were misclassified as controls. Moreover, the genetic associations discovered were not replicated in an independent cohort.

By undertaking the largest two-stage GWAS of VVs to date, we aimed to advance substantially our understanding of the aetiopathology and genetic architecture of VVs; discover clinically-relevant biologic pathways and prioritise targets for therapeutic development; and through genetic risk scoring pave the way for personalised medicine approaches to VV management.

## Methods

### Ethical approval

UK Biobank obtained ethical approval from the North West Multi-Centre Research Ethics Committee (MREC) (11/NW/0382). This study was conducted under UK Biobank study ID 10948. All 23andMe research participants included in this study provided informed consent for their genotype data to be utilised for research purposes under a protocol approved by the external AAHRPP-accredited IRB, Ethical & Independent Review Services.

### Population and phenotype definition

The characteristics of the full UK Biobank cohort are described in detail elsewhere.^18^ In the discovery analysis, VV cases were identified from the UK Biobank data showcase using diagnostic, operative and self-report codes (Supplementary Table 1). Following quality control, we identified 22,473 VV cases and the remaining 379,183 subjects were defined as controls.

In the 23andMe replication cohort, participants answered the question *‘Do you have varicose veins on your legs? (Yes/Not Sure/No)*. VV cases were identified if they answered *‘Yes’*, whilst controls were identified by answering *‘No’.* 113,041 VV cases and 295,928 controls were included in the final replication analysis.

### Genotyping

UK Biobank subjects were genotyped on UK BiLeve Axiom and UK Biobank Axiom arrays, with ~95% shared content. 805,426 directly genotyped variants were available prior to quality control. The 23andMe replication cohort was genotyped on one of four custom arrays (v1/v2, v3, v4, v5). Illumina HumanHap550+ BeadChip was used for v1/v2 (1,680 cases, 4,882 controls) and the Illumina OmniExpress+ BeadChip was used for v3 (21,342 cases, 56,448 controls). For v4 a fully customized array (58,883 cases, 148,637 controls) was used, and for v5, the Illumina Infinium Global Screening Array was used (31,136 cases, 85,961 controls). Successive arrays contained significant overlap with all previous arrays.

### Quality control

Quality control (QC) for the UK Biobank discovery cohort was conducted using PLINK v1.9^19^ and R v3.3.1, as previously described (Supplementary Figure 1).^20^ Following the QC protocol, our final discovery GWAS consisted of 401,667 subjects and 547,011 genotyped single-nucleotide polymorphisms (SNPs).

For the 23andMe replication analysis, samples were restricted to individuals from European ancestry determined through an analysis of local ancestry.^21^ A maximal set of unrelated individuals was chosen for each analysis using a segmental identity-by-descent (IBD) estimation algorithm, defining related individuals if they shared more than 700 cM IBD, including regions where the two individuals share either one or both genomic segments IBD. Cases were preferentially retained in the analysis. Variant QC was applied independently to genotyped and imputed GWAS results. The SNPs failing QC were flagged based on multiple criteria, such as HWE P-value, call rate, imputation R-square and test statistics of batch effects.

### Imputation

The phasing and imputation of UK Biobank has been previously described.^22^ In 23andMe, out-of-sample modified versions of the Beagle graph-based haplotype phasing algorithm^23^ and Eagle2^24^ algorithm were used to phase samples. We imputed samples against a single unified imputation reference panel combining the 1000 Genomes Phase 3 haplotypes^25^ with the UK10K imputation reference panel^26^ using Minimac3.^27^

### Association analysis

In the UK Biobank, we performed GWAS using a linear mixed non-infinitesimal model implemented in BOLT-LMM v2.323^28^ across 547,011 genotyped SNPs (minor allele frequency (MAF) ≥ 0.01) and 8,397,536 imputed SNPs (MAF ≥ 0.01 and INFO score ≥ 0.90), adjusting for genotyping platform and genetic sex. Conditional regression analysis was performed at the top signal at each of 109 associated loci in BOLT-LMM^28^, excluding the MHC region.

In 23andMe, summary statistics were generated via logistic regression assuming an additive model for allelic effects. Association analysis was performed adjusting for age, sex, the first five principal components, and genotyping platform. The top 118 independent variants from the discovery GWAS were tested for their association with VVs in the 23andMe cohort. The Bonferroni-corrected significance threshold for replication was set at P < 4.24×10^−4^ (0.05/118). Data for 108 of 118 variants was available; nine variants failed to meet the SNP QC within 23andMe, and one (10:79677281_CA_C) was not identified in 23andMe. A fixed-effects meta-analysis was performed in GWAMA.^29^

### Genetic risk score

Using the 49 replicated signals, we calculated a weighted genetic risk score (wGRS) for individuals in the UK Biobank cohort as previously described.^30^ The effect alleles for each variant were used to compute SNP dosage, using QCTOOL v2. wGRS calculations and unpaired t-testing between the different subgroups was performed in R v3.3.1.

### Functional annotation of SNPs

To annotate SNPs at our susceptibility loci, SNP2GENE was performed in Functional Mapping and Annotation of GWAS (FUMA) v1.3.3^31^ using summary statistics from the UK Biobank discovery cohort and default settings. FUMA defined risk loci borders by using all independent genome-wide significant SNPs (r^2^ < 0.6), and identified all SNPs that were in linkage disequilibrium with one of these candidate SNPs. Using ANNOVAR, candidate SNPs within the replicated loci were annotated on the basis of genomic location. Exonic SNPs were investigated further using Gnomad and Ensembl genome browsers to uncover non-synonymous variants. All candidate SNPs were annotated with Combined Annotation Dependent Depletion (CADD)^32^, RegulomeDB^33^, and 15-core chromatin states (predicted by the Roadmap ChromHMM model^34^) to predict any regulatory or transcription effects from chromatin states at each SNP.

### Candidate gene mapping

Four gene mapping approaches – positional mapping, eQTL mapping, MAGMA gene mapping, and summary-based mendelian randomisation (SMR) – were used to map putative genes at the replicated loci. For FUMA positional mapping, all genome-wide significant SNPs at each locus were mapped to genes within a positional window of 10Kb.^31^ eQTL gene mapping was used to map genes based on having at least one genome-wide significant cis-eQTL in GTEx v8 tibial artery tissue.^31^ Using MAGMA v.1.07^35^, we conducted a genome-wide, gene-based association study, testing 17,966 protein-coding genes. The significance threshold was set at P < 2.78×10^−6^ (0.05/17966).

To identify gene expression levels associated with VVs due to pleiotropy, SMR and HEIDI analyses were performed.^36^ Summary statistics from the UK Biobank discovery GWAS were used, alongside eQTL data for GTEx V7 tibial artery. The top associated eQTL for each gene was used as an instrumental variable to examine association with VVs. A Bonferroni-corrected significance for all SMR probes was set at P_SMR_ < 1.01×10^−5^ (0.05/4946 probes). Subsequently, a HEIDI test was conducted across 44 SMR-significant probes to examine for heterogeneity in SMR estimates (significance set at P < 1.12×10^−3^ (0.05/44)).

### Gene set, tissue-specific, and pathway enrichment analysis

Gene set analysis was implemented in MAGMA v1.07^35^ across 15,496 gene sets derived from MSigDB v8.0^37^ with a significance threshold of P < 3.23×10^−6^ (0.05/15496). Tissue-specific gene property analysis was also performed in MAGMA v1.07 to determine the expression of the protein-coding genes in VV-related tissue types in GTEx v8.0.^38^

Pathway analysis was performed in eXploring Genomic Relations (XGR).^39^ XGR ontology enrichment analysis was performed across all mapped genes, in ‘canonical pathways’ with the following settings: hypergeometric distribution testing, any number of genes annotated, any overlap with input genes, and an adjusted FDR < 0.05.

### SNP-based heritability and genetic correlation

Linkage Disequilibrium Score (LDSC) regression^40^ was used to calculate the LDSC intercept, mean chi-squared test and attenuation score, and SNP-based heritability (h^2^_g_).^41^ LDSC regression was also used to calculate the genetic correlation between VVs and 51 preselected traits, across nine trait categories: metabolites, glycaemic traits, autoimmune diseases, anthropometric traits, smoking behaviour, lipids, cardiometabolic traits, reproductive traits and haematological traits, from publicly-available GWAS data within LDHub. LDHub is a centralised database which contains summary-level GWAS data from 855 GWAS studies across 41 GWAS consortia.^41^ Traits were selected based on putative epidemiological associations with VVs from the literature.

### Drug target enrichment analysis

The prioritised genes at our replicated loci were queried on the Open Targets Platform,^42^ assessing whether encoded proteins were tractable to small molecule or antibody targeting, or drug targets in any phase of clinical trial. Genes were also analysed for enriched drug pathways, with a nominal P < 0.05 threshold of significance.

## Results

The overall analytic workflow is shown in Figure 1.

**Figure 1.**
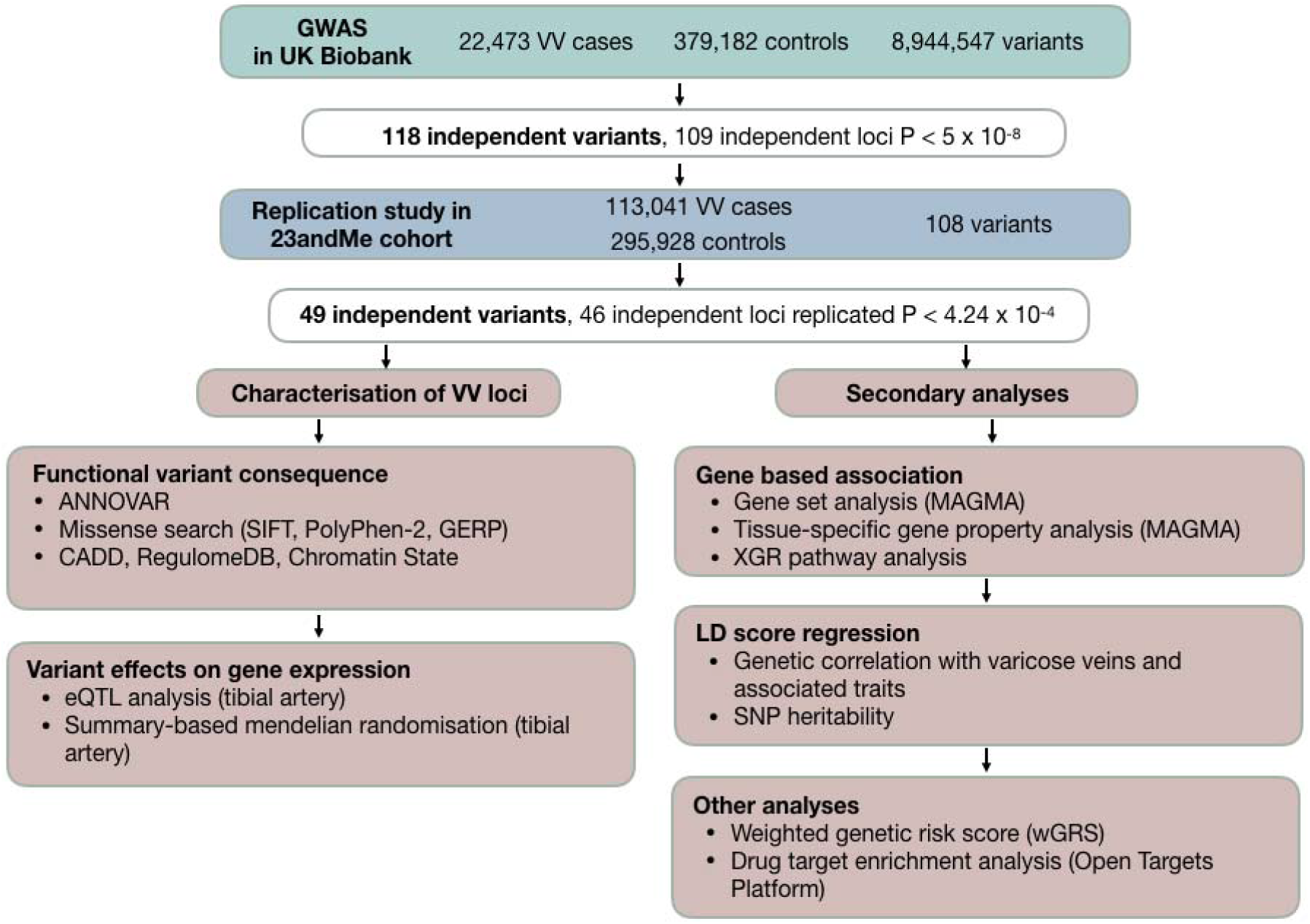
Study design and analysis workflow. A discovery GWAS was performed in the UK Biobank cohort, following which, the top independent lead variants were tested within the 23andMe replication cohort. Of the 118 tested variants, data on 117 variants was available for replication in the 23andMe Cohort, of which 108 passed quality control within the replication cohort (see Methods). In total, 49 independent variants at 46 loci met the Bonferroni-corrected threshold in the replication cohort (presented in Table 1), and were interrogated further in subsequent analyses.

**Table 1.**
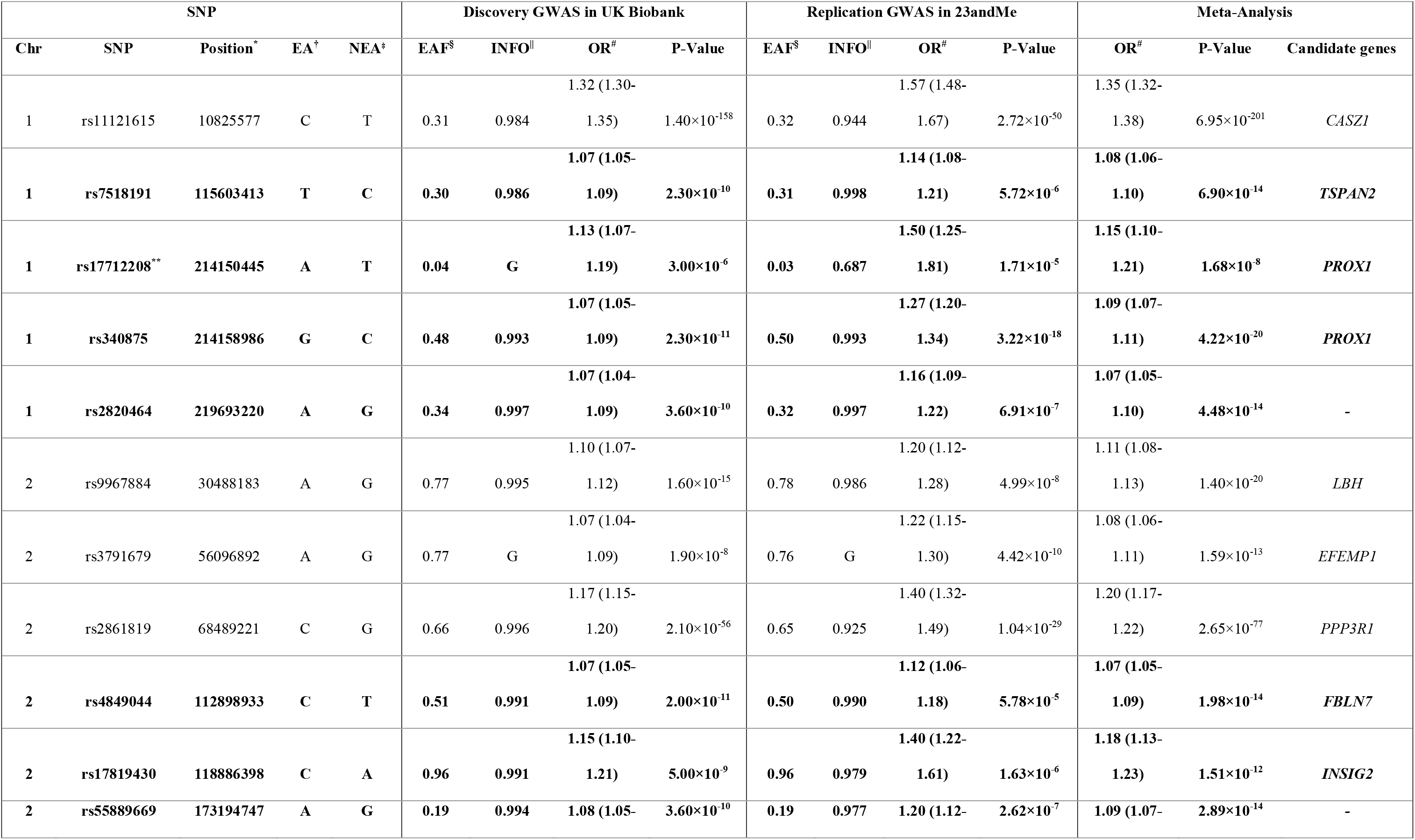
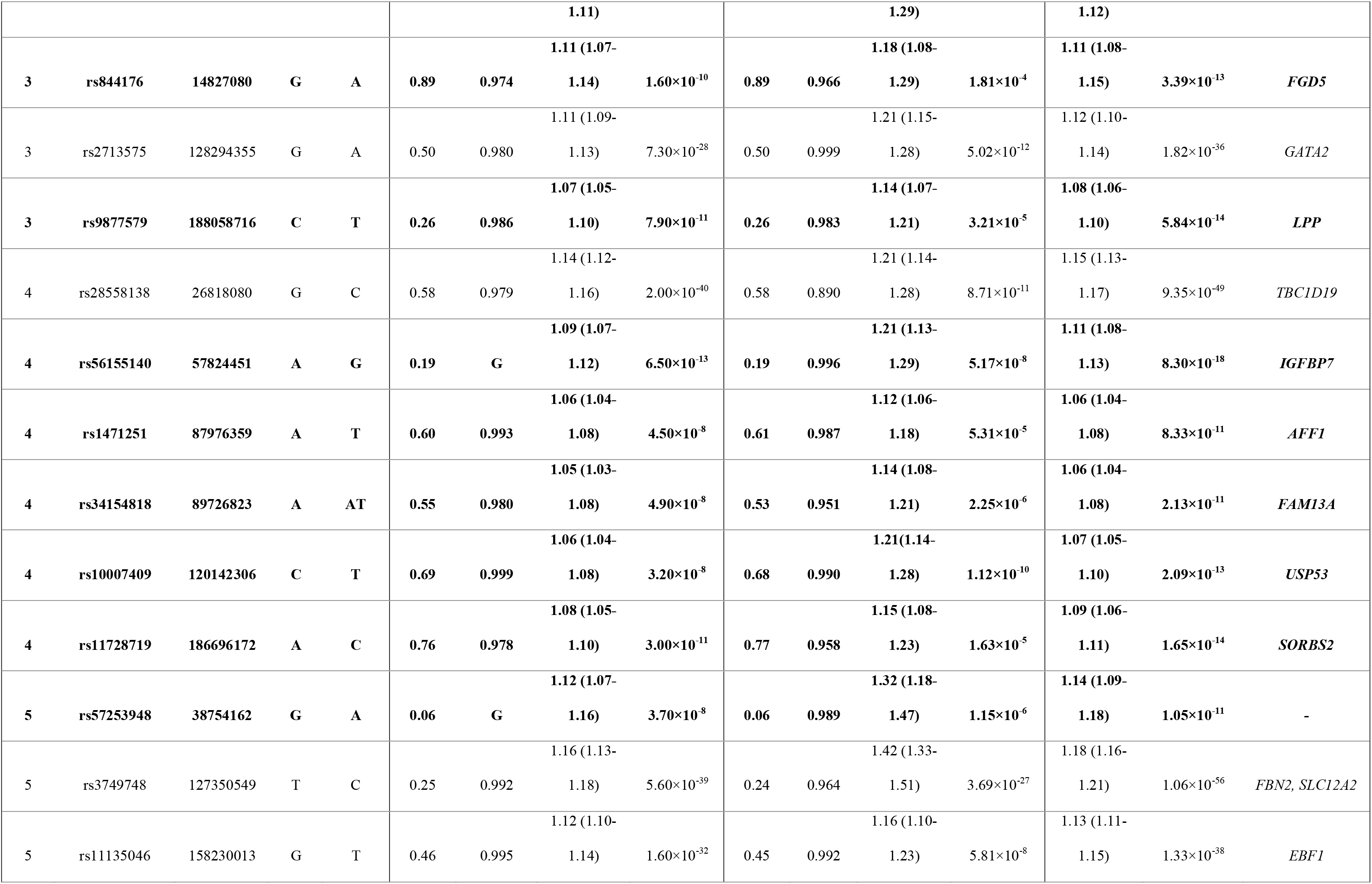
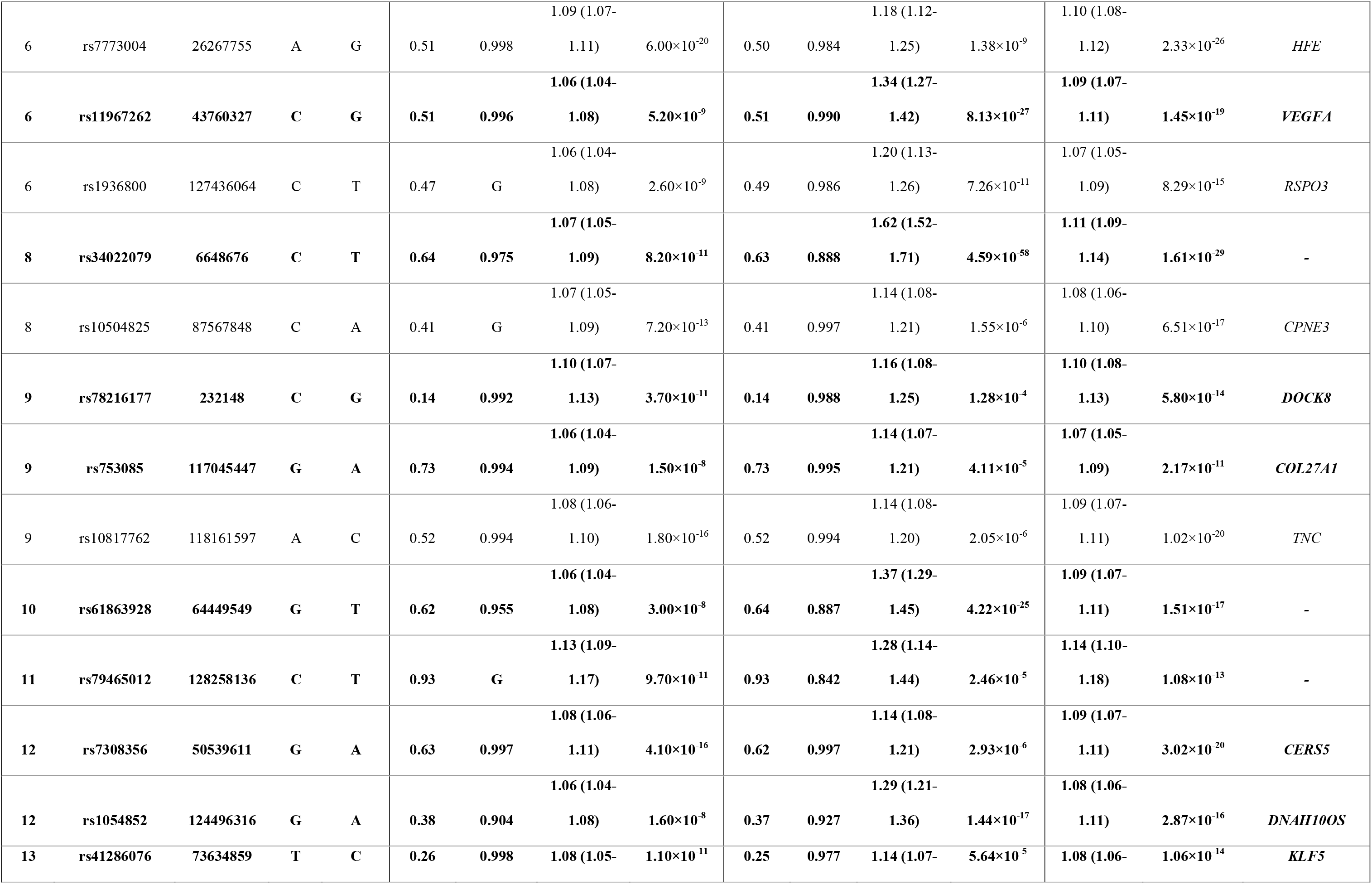
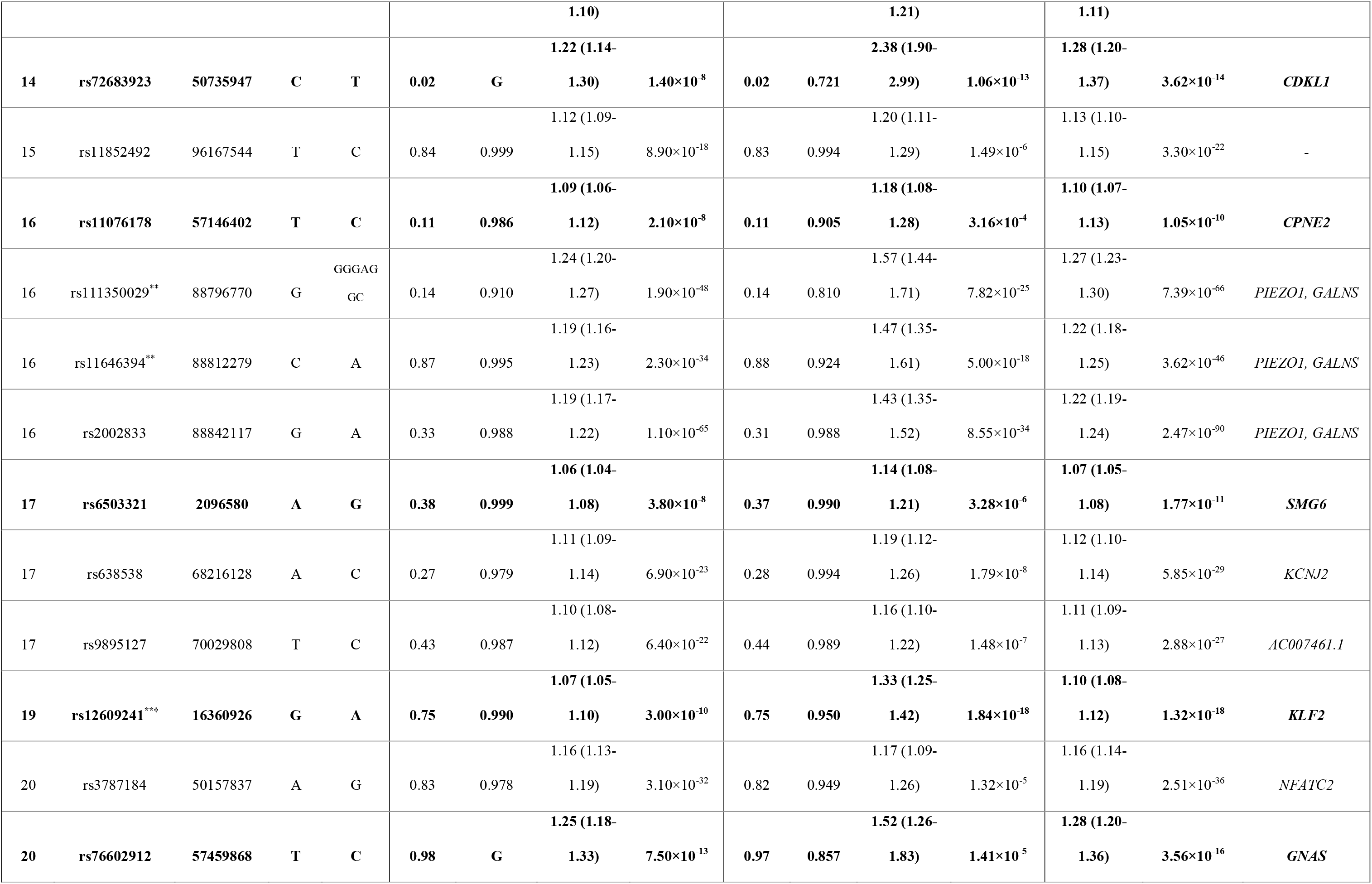
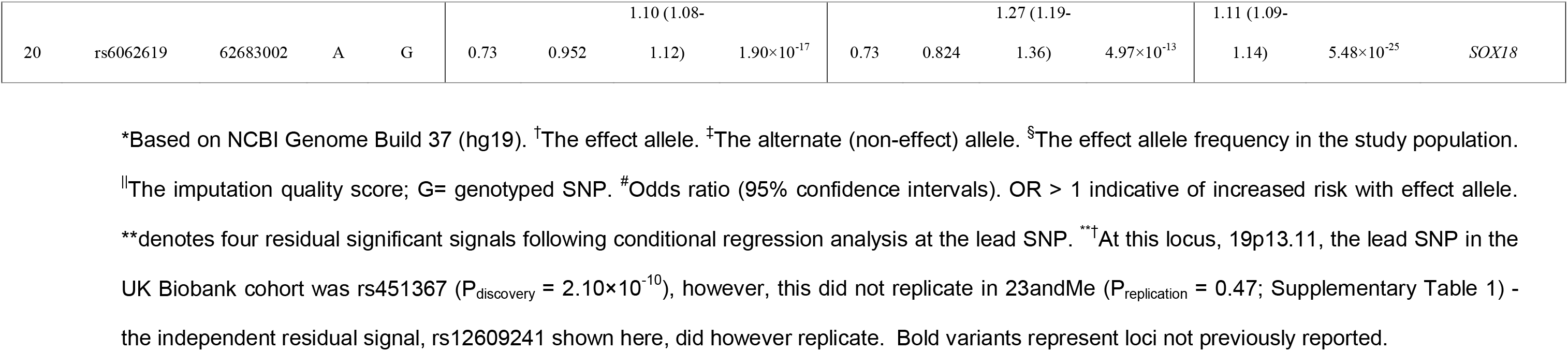
Forty-nine variants at 46 susceptibility loci significantly associated with varicose veins.

### Association analysis

GWAS of the UK Biobank discovery cohort (22,473 cases and 379,183 controls) yielded genome-wide significant associations (P < 5×10^−8^) at 109 risk loci (12,391 variants; Supplementary Data 1). Conditional regression yielded a further nine independent signals at eight of the 109 loci. As expected, the large discovery sample led to the λ_GC_ showing nominal inflation (1.25). The linkage disequilibrium score (LDSC) regression intercept (1.06), and attenuation ratio of 0.13 is fully in keeping with expectations of polygenicity (Figure 2).^40^

**Figure 2.**
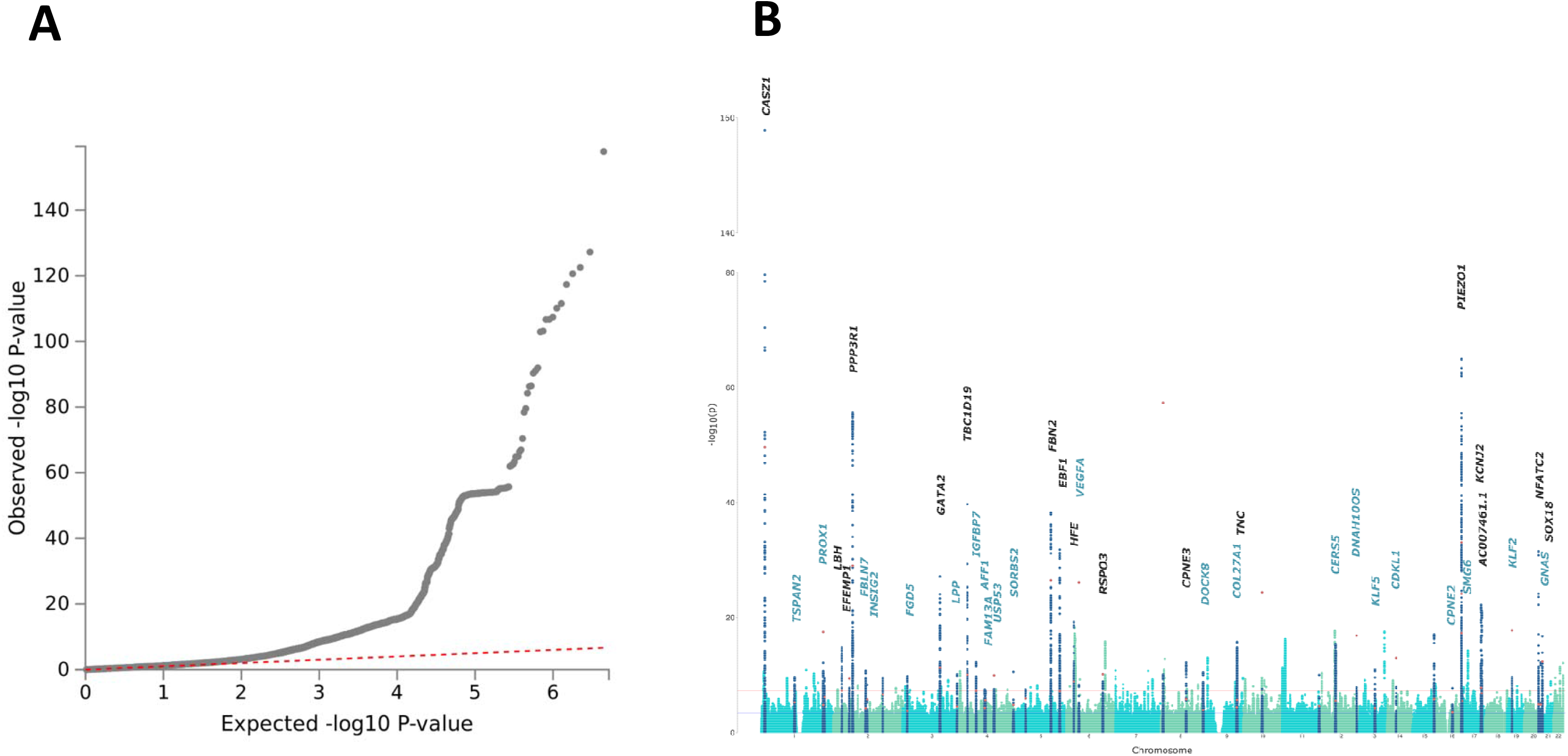
Results of genome-wide association study in varicose veins. A) QQ plot of observed vs. expected P-values. B) Manhattan plot showing genome-wide P-values plotted against position on each of the autosomes. The dark blue, light blue, and green dots refer to the discovery UK Biobank Cohort, with the red dots corresponding to the 49 variants from the 23andMe cohort at each replicated locus (shown in Table 1). The dark blue peaks correspond to the 46 loci that replicated in the 23andMe cohort at a Bonferroni-corrected threshold of P < 4.24×10^−4^. Candidate genes at each locus are named above each signal, with newly discovered genetic loci in blue, and previously described loci in black.

We tested the top 118 signals at the 109 risk loci in the 23andMe dataset (113,041 cases and 295,928 controls). Again, the LDSC intercept demonstrated moderate inflation (λ = 1.13, S.E. = 0.01) consistent with polygenicity and large sample size. Forty-nine of 118 variants demonstrated significant association at a conservative Bonferroni-corrected threshold of P < 4.24×10^−4^. Thus, we identified 49 independent significant associations at 46 risk loci (Figure 2; regional association plots for all 49 signals can be found in Supplementary Figure 2). Allelic effects were concordant across both cohorts at all 49 replicated variants, with minimal evidence of heterogeneity between the two GWAS at all loci (Q-statistic > 0.05). Eighteen loci were previously reported, and 28 are novel (Table 1). Sixty-nine variants at 63 risk loci did not replicate (Supplementary Table 2).

### Genetic risk score

As expected, we found the weighted genetic risk score (wGRS) for VV cases (5.179) was higher than in controls (4.986; P = 9.90×10^−324^; Table 2). We hypothesized that VV cases that had required surgery would have a higher wGRS. In keeping with this, we discovered a higher wGRS in the operation group (5.185) compared with the non-operation group (5.076; P = 2.46×10^−13^). These results strongly suggest that the wGRS derived from our susceptibility loci correlate with more severe VVs.

**Table 2.** Weighted genetic risk score for varicose veins in the UK Biobank cohort.

| Group | VV |  | P-Value<sup>†</sup> | VV cases<br>with<br>operation<br>code | VV cases<br>without<br>operation<br>code | P-Value<sup>§</sup> |
| --- | --- | --- | --- | --- | --- | --- |
|  | cases | Controls |  |  |  |  |
| **N** | 22,473 | 379,183 |  | 21,407 | 1,066 |  |
| **Mean wGRS\*** | 5.179 | 4.986 | 9.90×10<sup>-324</sup> | 5.185 (0.453) | 5.076 (0.468) | 2.46×10<sup>-13</sup> |
| **(standard deviation)** | (0.454) | (0.451) |  |  |  |  |

\*wGRS: weighted genetic risk score. <sup>†</sup>Unpaired two-tailed t-test between varicose veins cases and controls. <sup>§</sup>Unpaired two-tailed t-test between varicose veins cases with an operation code and varicose veins cases without an operation code.

### *In silico* annotation

FUMA identified 5,315 genome-wide significant candidate SNPs from our discovery cohort associated with VVs at 45 of the 46 replicated loci (Figure 3; Supplementary Data 2). Around 2% of candidate SNPs (n = 103) were exonic, of which 56 were non-synonymous (52 missense, two stop gain, one splice site variant, and one frameshift variant). Of the non-synonymous variants, four missense variants were predicted to affect protein structure or function, and were in moderate linkage disequilibrium (R^2^ ≥ 0.22 and D’ ≥ 0.75) with the index SNP at three loci – 12q13.12 (index SNP: rs7308356), 16q24.3 (index SNP: rs2002833) and 17q24.3 (index SNP: rs9895127) (Supplementary Table 3). This includes rs7184427 (A/G) (P = 9.10×10^−40^, OR = 1.19, R^2^_index_ = 0.22, D’_index_ = 0.75), which causes a predicted deleterious p.Val250Ala substitution within *PIEZO1* (SIFT: 0). *PIEZO1* encodes a mechanically active ion channel involved in the detection of vascular shear stress, and was previously associated with VVs and lymphoedema.^17,43^ Furthermore, rs12303082 (P = 2×10^−9^, OR = 1.06, R^2^_index_ = 0.27, D’_index_ = 0.92) resides within a highly conserved region (Genomic evolutionary rate profiling (GERP) score: 0.83) in *FAM186A*, and causes a non-conservative p.Lys187Gln amino acid substitution that is predicted to be deleterious and alter protein function (SIFT: 0.01, PolyPhen-2: 0.91).

**Figure 3.**
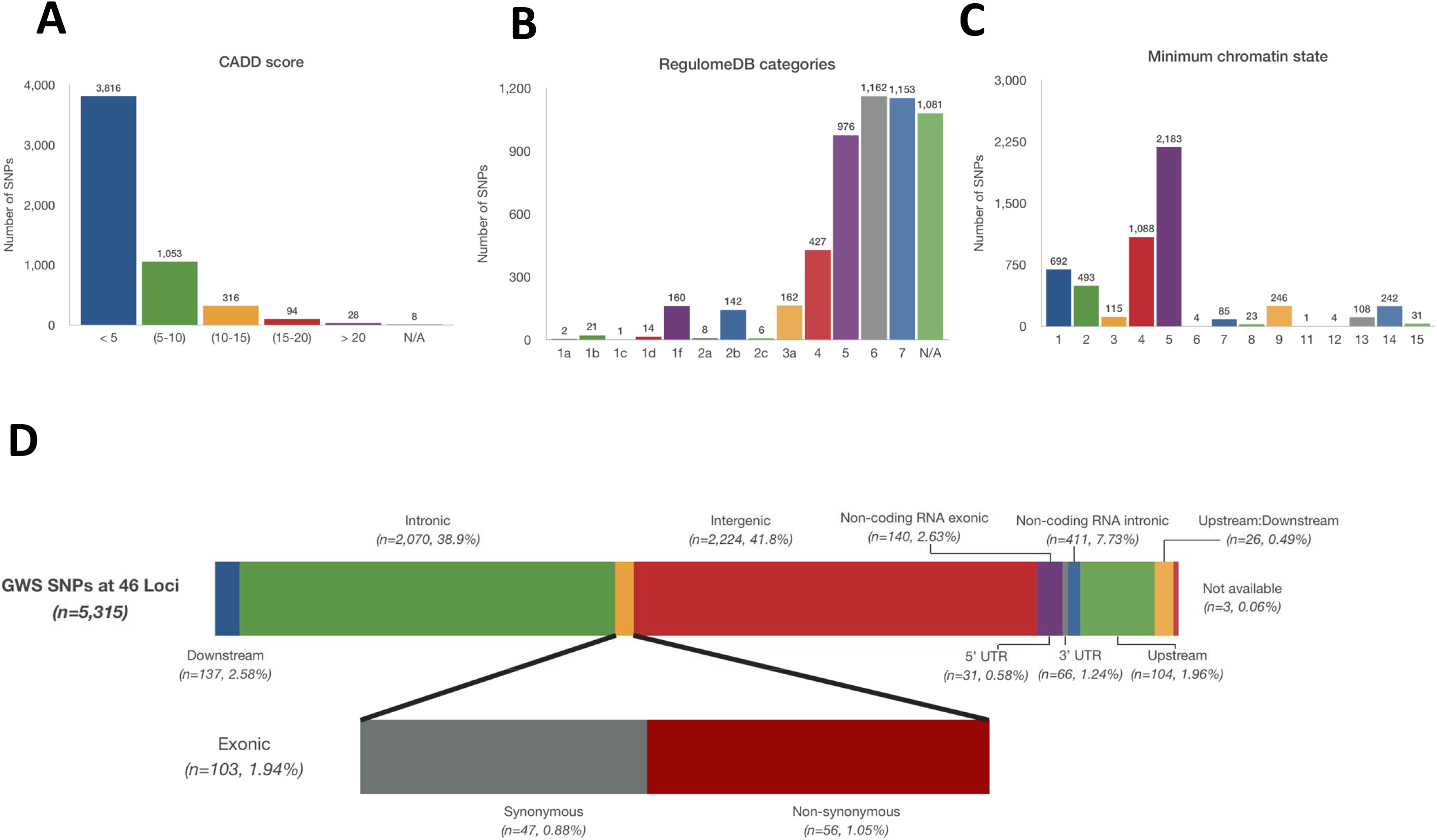
Functional annotation of the 5,315 genome-wide significant SNPs at our 46 replicated loci. Functional consequences of the SNPs on genes were obtained by performing ANNOVAR gene-based annotation using Ensembl genes (build 85) in FUMA. A) CADD scores, B) RegulomeDB scores and C) 15-core chromatin state were annotated to all 5,315 SNPs in 1000G phase 3 by FUMA through matching chromosome, position, reference, and alternative alleles. D) the location of the 5,315 SNPs are shown.

Of the 4,294 intronic and intergenic variants identified by FUMA (Figure 3), 3,735 (87.0%) resided in open chromatin regions (Supplementary Data 2), and 163 demonstrated evidence of functionality with CADD (Combined Annotation-Dependent Depletion^32^) score ≥ 12.37, the threshold suggested for deleterious variants (Supplementary Table 4). Using RegulomeDB (RDB^33^) to investigate their regulatory functions, 17 had a RDB score of at least 2b (likely to affect binding) and eight had an RDB score of at least 1f (likely to affect binding and linked to expression of a gene target) (Supplementary Table 4).

### Gene mapping

Positional mapping in FUMA SNP2GENE highlighted 204 genes based on genomic position at 38 loci (Supplementary Table 5).^31^ eQTL mapping, based on GTEx v8 tibial artery tissue^38^, mapped a total of 80 genes (Supplementary Table 5). Genome-wide, gene-based association analysis implemented in MAGMA v1.07^35^ identified 248 protein-coding genes significantly associated with VVs (P < 2.67×10^−6^); 117 were within our replicated loci (Supplementary Table 6; Supplementary Figure 3).

In the summary-based mendelian randomisation (SMR) analysis^36^, 44 genes passed the set significance threshold (P_SMR_ < 1.01×10^−5^). To exclude SMR associations due to linkage, we performed HEIDI analysis across 44 significant genes — 27 passed the HEIDI test (P_HEIDI_ ≥ 1.12×10^−3^; Supplementary Table 7), 14 of which were within our VVs susceptibility loci, highlighting an association through pleiotropy rather than linkage disequilibrium and co-localisation.

In summary, 237 unique genes were mapped by at least one mapping approach. Significant overlap between the mapping strategies was seen, with the majority of genes (54.9%, n = 130) being prioritised by two or more mapping approaches. Thirty-six genes were prioritised by three mapping approaches, and six genes (*ATF1, AP1M1, DNAH10OS, FBLN7, LBH, WDR92*) were prioritised by all four approaches (Supplementary Table 5).

### Gene set, tissue-specific and pathway enrichment

Following MAGMA gene set enrichment analysis, four Gene Ontology (GO) gene sets were significantly over-represented in our data: Cardiovascular Development (P = 1.56×10^−8^, n = 666); Tube Morphogenesis (P = 9.35×10^−8^, n = 778); Blood Vessel Morphogenesis (P = 9.39×10^−7^, n = 555); and Tube Development (P = 1.68×10^−6^, n = 956) (Supplementary Table 8). Further, tissue-specific gene property analysis demonstrated significant gene expression in all three vascular tissue types present in GTEx 54 tissue types: coronary artery (P = 6.23×10^−7^, 2^nd^ most enriched), tibial artery (P = 1.05×10^−6^, 3^rd^ most enriched) and aorta (P = 3.92×10^−5^, 8^th^ most enriched; Figure 4). MAGMA analysis of GTEx 30 general tissue types also demonstrated blood vessels to be a highly enriched tissue (P = 3.8×10^−4^, 3^rd^ most enriched; Figure 4). Using XGR^39^, six canonical pathways were significantly enriched, including genes in pathways related to extracellular matrix biology, the VEGF and VEGFR signalling network, and intracellular Ca^2+^ signalling in the T-Cell Receptor (TCR) Pathway (Table 3).

**Figure 4.**
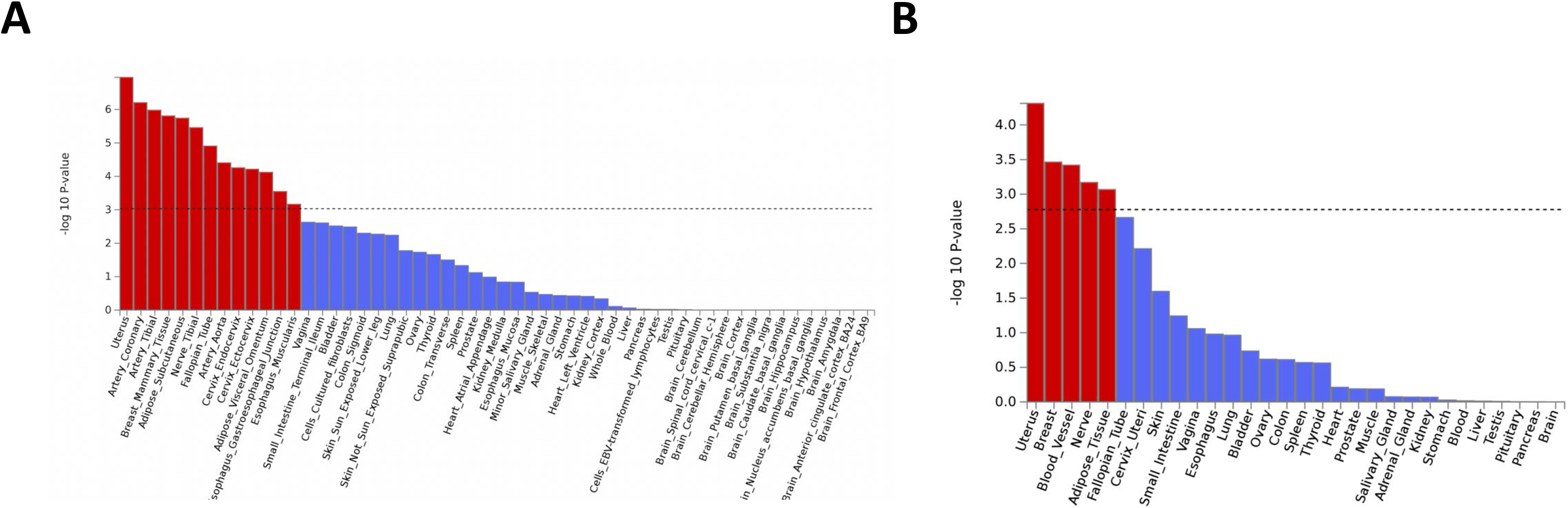
MAGMA tissue expression analysis. MAGMA Tissue Expression Analysis of varicose veins GWAS-summary data, implemented in FUMA in A) 54 specific and B) 30 general tissue types. This analysis tests the relationship between highly-expressed genes in a specific tissue and the genetic associations from the GWAS. Gene-property analysis is performed using average expression of genes per tissue type as a gene covariate. Gene expression values are log2 transformed average RPKM (Read Per Kilobase Per Million) per tissue type after winsorization at 50, and are based on GTEx v8 RNA-Seq data across 54 specific tissue types and 30 general tissue types. The dotted line indicates the Bonferroni-corrected α level, and the tissues that meet this significance threshold are highlighted in red.

**Table 3.** Gene-based enrichment analysis.

| Biological Process | Z-Score | P-Value | FDR | Number of overlapped genes | Genes |
| --- | --- | --- | --- | --- | --- |
| **Alpha9 beta1 integrin signalling events** | 3.85 | 0.0012 | 0.032 | 2 | *TNC, VEGFA* |
| **Genes encoding structural ECM glycoproteins** | 3.39 | 0.0012 | 0.032 | 6 | *EFEMP1, FBLN7, FBN2, IGFBP7, RSPO3, TNC* |
| **Calcium signalling in the CD4+ TCR pathway** | 3.5 | 0.0019 | 0.032 | 2 | *NFATC2, PPP3R1* |
| **Ensemble of genes encoding core extracellular matrix including ECM glycoproteins, collagens and proteoglycans** | 3.11 | 0.0019 | 0.032 | 7 | *COL27A1, EFEMP1, FBLN7, FBN2, IGFBP7, RSPO3, TNC* |
| **Non-canonical WNT signalling pathway** | 3.29 | 0.0025 | 0.035 | 2 | *MAPK10, NFATC2* |
| **VEGF and VEGFR signalling network** | 3.11 | 0.0032 | 0.036 | 1 | *VEGFA* |

### Genetic correlations with other phenotypes

Using LDSC regression^40^, we estimated the total SNP heritability (h^2^_g_) for VVs in UK Biobank to be 5.03% (S.E. = 0.30%) and in 23andMe to be 5.40% (S.E. = 0.30%). Of the nine trait categories tested for genetic correlation, two (anthropometric and autoimmune) contained traits that met our Bonferroni-corrected significance threshold (P < 5.56×10^−3^; Supplementary Table 9). All twelve significantly correlated traits were positively correlated with VVs (r_g_ range: 9-21%; Figure 5). Eleven traits belonged to the anthropometric category and pertained to height and weight phenotypes, which are well-known, established risk factors for VVs.^3,11,12^ In the autoimmune trait category, we discovered systemic lupus erythematosus (SLE) to be correlated with VVs, sharing ~19% genetic overlap (P = 4.2×10^−3^, r_g_ = 0.195). Of note, the C allele of variant rs17321999 at 2p23.1 (*LBH*), which is associated with an increased risk of SLE (P = 2.22×10^−16^, OR = 1.20)^44^, was also significantly associated with VVs in our discovery cohort (P_disc_ = 3.20×10^−14,^ OR = 1.09), and in high linkage disequilibrium with lead SNP at 2p23.1 in the meta-analysis (rs9967884; P_meta_ = 1.40×10^−14,^ OR = 1.11; r^2^ = 0.87).

**Figure 5.**
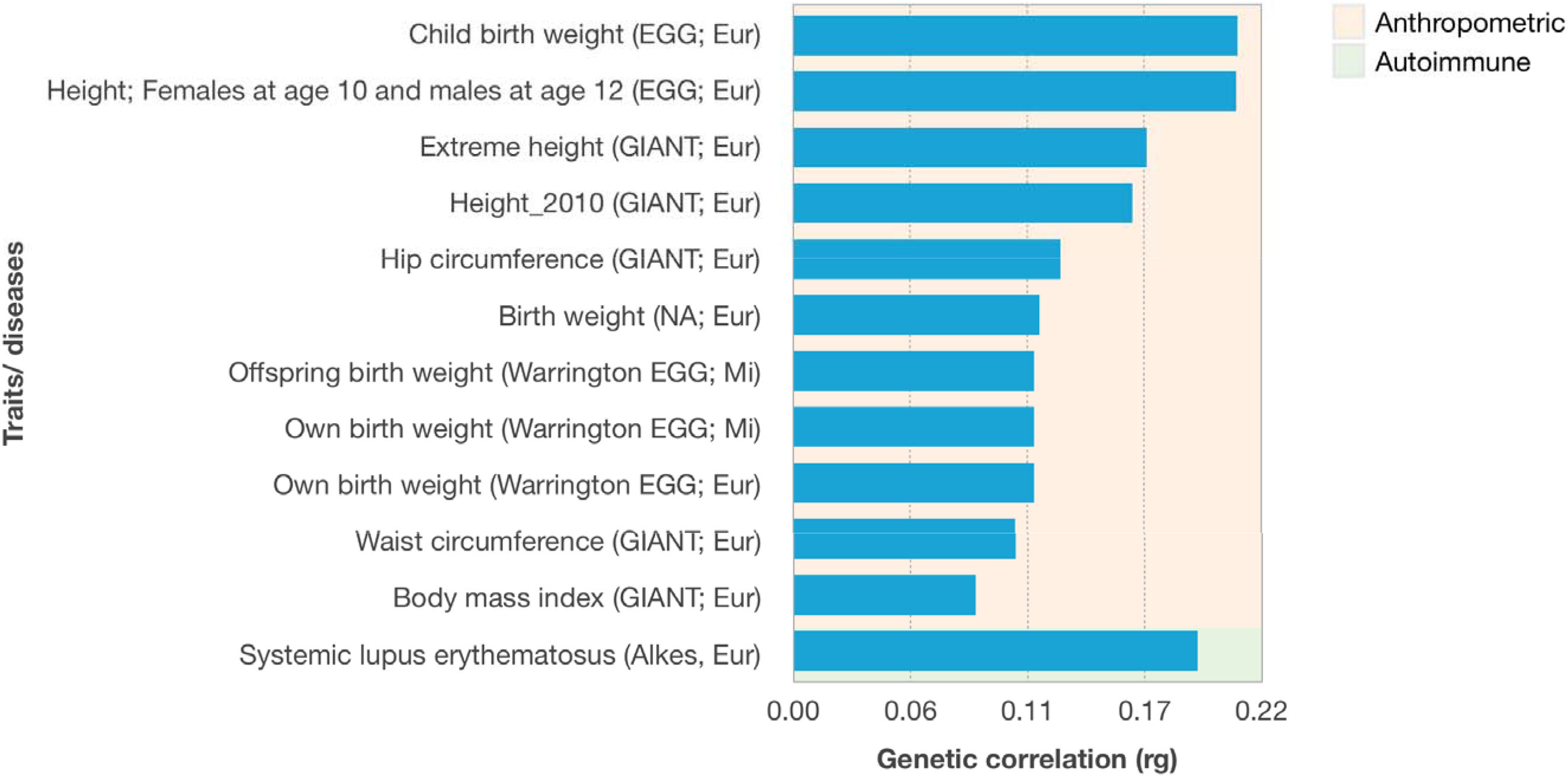
Genetic correlation between varicose veins and other traits and diseases. Genetic correlation (r_g_) between varicose veins and publicly-available phenotypes in LD Hub, using LD Score regression. All twelve traits have met a Bonferroni-corrected significance P < 5.56×10^−3^ and are positively correlated with varicose veins. The consortium and ethnicity for each study is provided in parentheses. EGG, Early Growth Genetics Consortium; GIANT, Genetic Investigation of ANthropometric Traits Consortium; Alkes, Alkes Group (Harvard T.H Chan School of Public Health); NA, Not Applicable. Eur, European ethnicity; Mi, Mixed ethnicity. Further details regarding the studies are provided in Supplementary Table 9.

### Drug target enrichment analysis

Of the 237 mapped genes, 200 genetic targets were identified.^42^ 42 drug pathways reached nominal significance (P < 0.05), with the Butyrophillin family interactions pathway being most enriched (P = 2.6×10^−7^, six targets), followed by the Calcineurin-NFAT pathway (P= 9.4×10^−4^, two targets), and the Transcriptional regulation by RUNX1 pathway, which possessed the highest number of targets (P = 1.6×10^−3^, 14 targets; Supplementary Table 10). Tractability information for 105 gene targets was available, with 65 of 237 genes predicted to be tractable to antibody targeting to a high confidence, and 26 genes predicted to be tractable to small molecule targeting (Supplementary Table 11). Eight of the 237 genes overlapped with pharmacologically active targets with known pharmaceutical interactions (*CDK10, COL27A1, GABBR1, KCNJ2, MAPK10, OPRL1, TNC and VEGFA*) (Supplementary Table 12). Of note, *VEGFA* is a target for several antibody, protein and oligosaccharide agents which are currently in phase 2, 3 and 4 clinical trials, including for several ocular vascular disorders (Supplementary Table 12).

## Discussion

VVs cause significant morbidity and large healthcare costs. There is an urgent need to understand the biology of VVs in order to develop new therapeutic strategies. Our analysis represents the largest and most comprehensive GWAS to date of VVs, including 135,514 patients and 675,111 controls. We discovered 28 new risk loci (29 new signals), and independently replicated 18 of 29 previously reported but non-replicated loci (20 of 32 known signals). Our weighted genetic risk score correlates with more severe prognosis – a first step in facilitating personalised medicine approaches to management. Furthermore, our *in silico* analyses demonstrated strong evidence of functional variants in VV-associated genomic regions. Our pathway analyses establish a strong enrichment for genes expressed in the extracellular matrix, immune cell signalling and circulatory system development. We have also identified a novel genetic correlation between VVs and systemic lupus erythematosus (SLE). Several prioritised genes demonstrate the potential for pharmacological targeting, and are currently under active investigation in other diseases.

### Genetic risk score

Over two million people in the USA have advanced chronic venous disease^2^, and around 500,000 per year require invasive surgical procedures. Our genetic risk score represents a proof-of-principle, demonstrating the feasibility of enhanced prognostication in enabling the subset of VVs patients that will require surgical intervention to be defined. Preventive strategies in high risk individuals could include prophylactic compression stocking use, or early ablation procedures to mitigate risk of venous ulceration. The benefits of early endovenous ablation in improving healing of venous leg ulcers has been demonstrated.^45^ Future research may also enable the identification of those at high risk of recurrence following surgery, a significant problem in current management of VVs patients.

### Biological drug targets

This study has identified several biological pathways as mediators of VV pathophysiology that may be of relevance to pharmacological targeting (Supplementary Table 13).

#### Extracellular matrix regulation

VVs demonstrate an increased luminal diameter and intimal hypertrophy, features closely related to disruption of the extracellular matrix (ECM).^46^ Deposition of ECM in the perivascular space in VVs is also recognised – possibly a compensatory mechanism to reinforce an already weakened wall.^47^ Venous dilatation and valve ring enlargement seen in VVs is thought to affect the ability of venous valves to co-apt, which in turn contributes to venous reflux and hypertension.^10^ Indeed, there is a noted imbalance of ECM proteins in VVs, specifically collagen and elastin - with a preponderance of collagen compared to normal vein.^48^ It is therefore possible that disruption in intrinsic connective tissue components of the vein wall or valves may contribute to VVs.

Our prioritised genes significantly overlapped with canonical pathways relating to ECM components, including Collagen Type XXVII Alpha 1 Chain (*COL27A1*) and EGF-containing Fibulin-like Extracellular Matrix Protein 1 (*EFEMP1)*.

rs753085 (P = 2.17×10^−11^, OR = 1.07) is in an intron of *COL27A1*. COL27A1 is a fibrillar collagen in the extracellular matrices of several tissues, including blood vessels.^49^ *COL27A1* expression has been demonstrated to be reduced in VV samples.^50^ Moreover, our drug enrichment analysis demonstrated COL27A1 to be a pharmacologically active target, with pharmaceutical agents currently being investigated in several clinical trials (Supplementary Table 12), demonstrating its potential candidacy as a therapeutic target for VV prevention or treatment.

In our GWAS, we replicated the previously described association between rs3791679 and VVs (P = 1.59×10^−13^, OR = 1.08).^16^ rs3791679 resides in an enhancer region of *EFEMP1*, which encodes the ECM glycoprotein fibulin-3.^51^ Fibulin-3 plays a key role in maintaining the integrity of elastic tissues^16^, and is highly expressed in vein endothelial cells. Fibulin-3 has been found to antagonise vascular development by reducing the expression of the matrix metalloproteases, MMP2 and MMP3, and increasing expression of tissue inhibitors of metalloproteases (TIMPs) in endothelial cells.^52^ The saphenofemoral junction in VVs demonstrates reduced expression of MMP2, and heightened expression of TIMP1 and MMP1 protein levels.^53^ Alterations in expression of these enzymes may therefore result in weakness in the venous wall which may predispose patients to VVs. Our drug-enrichment analysis found fibulin-3 to be tractable to antibody targeting to a high confidence; moreover, metformin has been demonstrated to down-regulate fibulin-3 through downregulation of MMP2.^54^ Fibulin-3 therefore represents another potential pharmacological target for VVs.

#### Immune response

Chronic Inflammation in the venous wall has been proposed to play a role in VV aetiopathology.^46^ When compared to normal veins, enhanced expression of inflammatory mediators has been observed.^10^ VVs contain increased mast cells, monocytes and macrophages compared to normal veins.^10^

We defined five inflammation-associated risk loci in our GWAS. Of particular note is rs78216177 (P_meta_ = 5.80×10^−14^, OR = 1.10), in an intron of *DOCK8* that has roles in both innate and adaptive immune systems. Deletion of *DOCK8* is strongly associated with Hyper-IgE syndrome, a type of primary immunodeficiency that affects multiple systems, including the vasculature.^55^ Vascular abnormalities in hyper-IgE syndrome include aneurysmal changes and abnormalities in great vessels.

We also discovered a significant genetic overlap between VVs and SLE. Lead variant rs1471251 (P = 8.33×10^−11^, OR = 1.06) is a known eQTL of *AFF1*, which is associated with SLE.^56^ Reinforcing this shared genetic basis, an associated variant at *LBH,* rs17321999 (P_disc_ = 3.20×10^−14,^ OR = 1.09), also increases the risk of SLE in a GWAS by Morris *et al* (P = 2.22×10^−16^, OR = 1.20).^44^

XGR analysis demonstrated enrichment of “intracellular calcium signalling in the CD4+ T-Cell Receptor (TCR) pathway” (P = 1.9×10^−3^, Z = 3.5), specifically highlighting genes *NFATC2* and *PPP3R1* which are intimately involved in this pathway. rs3787184 (P_meta_ = 2.51×10^−36^, OR = 1.16) is in an intron of *NFATC2,* and rs2861819 (P_meta_ = 2.65×10^−77^, OR = 1.20) is in an intergenic region ~19kb upstream of *PPP3R1*. *PPP3R1* encodes calcium binding B (CnB), a subunit of calcineurin, a Ca^2+^ influx-activated serine/threonine-specific phosphatase that interacts with NFAT transcription factors in the regulation of naive T-cell activation.^57^ VVs are defined by clustering and infiltration of T lymphocytes^47^, which are predominantly distributed close to the venous valve agger^58^, a fibroelastic structure located at the base of venous valves where media meets adventitia. Therefore, we hypothesize that aberrant *PPP3R1* and *NFATC2* expression could alter calcium signalling in T-cells which may contribute to the valvular pathology seen in VVs. The Calcineurin-NFAT pathway was also the second most enriched in our drug-target enrichment analysis, and may represent a novel therapeutic avenue in VVs.

#### Angiogenesis

Disruption in normal angiogenic processes can lead to VVs^59^, potentially because of a failure to develop properly-formed venous walls and valves, or to repair defects following vascular stress injury. MAGMA gene set analysis revealed enrichment of several gene sets relating to tube formation and morphology, with the VEGF/VEGFR signalling network also being enriched in the XGR analysis.

rs11967262 at 6p21.1 (P_meta_ = 1.45×10^−19^, OR = 1.09) lies in an intergenic region ~7kb upstream of vascular endothelial growth factor A (*VEGFA*). VEGFA is a critical regulator of angiogenesis, and is fundamental to maintaining the integrity and functionality of the vessel wall.^60^ VEGFA is a selective endothelial mitogen, binding to its receptor VEGFR2 to induce endothelial cell proliferation, migration and differentiation. VEGFA and VEGFR2 expression are significantly enhanced in the wall of VVs compared to normal veins, particularly in VVs complicated by thrombophlebitis.^61^ Plasma levels of VEGFA have also been demonstrated to be significantly increased in patients with VVs.^62^ VEGFA also causes vasodilatation, which may decrease vessel tone, and lead to stasis and the release of oxygen free radicals, which contribute to vein wall weakness.^63^ Additionally, VEGFA functions as a potent vascular permeability factor^64^, which may lead to VV progression and complications.^63^ Intriguingly, VEGFA also promotes inflammation via expression of intercellular and vascular cell adhesion molecules, linking angiogenesis to immune dysregulation.^65^ Of note, anti-VEGFA agents are currently being investigated in several clinical trials for the treatment of retinal vein occlusion (Supplementary Table 12). Furthermore, rs2713575 (P_meta_ = 1.82×10^−36^, OR = 1.12) at 3q21.3 lies in an intronic region near *GATA2*. *GATA2* is a haematopoietic transcription factor necessary for vascular integrity that acts downstream of VEGF, and has been demonstrated to regulate VEGF-induced angiogenesis and lymphangiogenesis.^66^ The VEGF axis may therefore be a promising candidate for therapeutic targeting in the treatment of VVs.

### Strengths and limitations

Certain limitations of this study must be acknowledged. While the discovery GWAS in UK Biobank used a combination of hospital diagnostic codes, operation codes and self-report codes, VV cases in 23andMe were identified based on self-report alone, meaning that the phenotyping for the replication GWAS was necessarily less stringent. Moreover, rather than undertake a formal meta-analysis between the discovery and replication GWAS, we tested the association only for the 118 independent lead SNPs that were genome-wide significant in the UK Biobank discovery GWAS. Thus, sub-threshold signals in the discovery GWAS that may have reached the significance threshold in the replication GWAS, or under meta-analysis, were not identified. Moreover, not having access to the full summary statistics for the replication GWAS also meant that our *in silico* analyses were performed on the summary statistics from the discovery GWAS alone. Finally, whilst we aimed to show the proof-of-principle behind genetic risk scoring and its clinical utility in the management of VVs, an independent validation cohort would enable improvement in its predictive capabilities.

However, several strengths of the paper mitigate these limitations. We have performed the largest GWAS of VVs to date by a considerable margin, with 135,514 cases and 675,111 controls. We have stringently controlled our false positive rate by reporting only the loci that were genome-wide significant in the discovery GWAS *and* that subsequently replicated, so we can be confident in the veracity of these 49 signals. This is reflected in the plethora of biologically plausible genes, gene clusters and biological pathways that were associated with these loci. This has also highlighted several novel therapeutic avenues that are currently under investigation. Furthermore, by including surgical codes for phenotyping in the UK Biobank discovery GWAS, we were able to identify a considerably greater number of cases than a previous GWAS that also used the UK Biobank resource but relied on ICD diagnostic codes alone (22,473 vs 9,577 cases).^17^ As a general principle of case ascertainment, we believe there is much to be gained by seeking out individuals who have undergone surgery for a disease: given the inevitable risk of complications, surgery is generally reserved for individuals at the more phenotypically severe end of the spectrum who have failed non-surgical treatment. We have demonstrated through our genetic risk score that the individuals who had undergone surgery for VVs were also at the *genotypically* more severe end of the spectrum. This finding lends further credence to the validity of the 49 identified loci, and opens the door to the future use of genetic risk scores in improving prognostication and guiding decision-making in the management of VVs patients.

## Conclusion

In conclusion, we have described the largest GWAS to date of VVs, a highly prevalent disease with a substantial health and socioeconomic cost. We discovered 49 variants at 46 loci that predispose to VVs. Importantly, our genetic risk score was higher in VV patients requiring surgery and, more immediately, represents a first step towards better prognostication. Furthermore, we have identified new associated pathways and genes involved in extracellular matrix regulation, inflammation, vascular and lymphatic development, smooth muscle cell activity, and apoptosis. These genes and pathways are biologically plausible contributors to the pathobiology of VVs, and provide excellent candidates for further investigation of venous biology. Several genes appear tractable to pharmacological targeting (notably *VEGFA*, *COL27A1*, *EFEMP1*, *PPP3R1* and *NFATC2*), and may represent viable therapeutic targets in the future management of VV patients.

## URLs

ANNOVAR, www.annovar.openbioinformatics.org/en/latest/; BOLT-LMM, www.data.broadinstitute.org/alkesgroup/BOLT-LMM/; CADD, cadd.gs.washington.edu/; ENSEMBL, www.ensembl.org/index.html; flashpca, github.com/gabraham/flashpca/; FUMA, www.fuma.ctglab.nl/; GERP, http://mendel.stanford.edu/SidowLab/downloads/gerp/; GTEx Portal, www.gtexportal.org/home/; GWAMA, https://genomics.ut.ee/en/tools/gwama; Human Genome Variation Society (HGVS), http://varnomen.hgvs.org/; HRC, www.haplotype-reference-consortium.org/; LD Hub, www.ldsc.broadinstitute.org/ldhub/; LD Link, www.ldlink.nci.nih.gov/; MAGMA, www.ctg.cncr.nl/software/magma; Open Targets Platform, www.targetvalidation.org/; PLINK, http://www.pngu.mgh.harvard.edu/~purcell/plink/; Polyphen-2, www.genetics.bwh.harvard.edu/pph2; QCTOOL,www.well.ox.ac.uk/~gav/qctool_v2/#overview; R, www.r-project.org; RegulomeDB, www.regulomedb.org/; SHAPEIT3, jmarchini.org/shapeit3/; SIFT, www.sift.bii.a-star.edu.sg/; UK Biobank, www.ukbiobank.ac.uk/; XGR, www.galahad.well.ox.ac.uk:3040; 1000 Genomes Project, www.1000genomes.org; 23andMe, https://research.23andme.com/

## Data availability

Full UK Biobank data can be accessed by direct application (see URLs). Discovery GWAS summary statistics from UK Biobank can be found in Supplementary Data 4.

## Acknowledgements

The data analysed in the present study was in part provided by the UK Biobank (www.ukbiobank.ac.uk), received under UK Biobank application no. 10948. The replication cohort was provided by 23andMe, Inc. (California).

D.F, A.W and K.Z. contributed to the conception, study design and supervision of this work. W.A, A.W and M.N contributed to data analysis. A.W and M.N. contributed to QC and imputation. W.W., A.A., and the 23andMe Research Team contributed to the replication study and analysis. R.L and A.H contributed to case ascertainment. W.A, A.W, D.F prepared the first draft of the manuscript. All co-authors made substantial contributions to data acquisition, data interpretation, and revised the work critically for important intellectual content.

The following members of the 23andMe Research Team contributed to this study: Michelle Agee, Stella Aslibekyan, Adam Auton, Robert K. Bell, Katarzyna Bryc, Sarah K. Clark, Sarah L. Elson, Kipper Fletez-Brant, Pierre Fontanillas, Nicholas A. Furlotte, Pooja M. Gandhi, Karl Heilbron, Barry Hicks, David A. Hinds, Karen E. Huber, Ethan M. Jewett, Yunxuan Jiang, Aaron Kleinman, Keng-Han Lin, Nadia K. Litterman, Marie K. Luff, Jennifer C. McCreight, Matthew H. McIntyre, Kimberly F. McManus, Joanna L. Mountain, Sahar V. Mozaffari, Priyanka Nandakumar, Elizabeth S. Noblin, Carrie A.M. Northover, Jared O’Connell, Aaron A. Petrakovitz, Steven J. Pitts, G. David Poznik, J. Fah Sathirapongsasuti, Anjali J. Shastri, Janie F. Shelton, Suyash Shringarpure, Chao Tian, Joyce Y. Tung, Robert J. Tunney, Vladimir Vacic, Xin Wang, Amir S. Zare.

## Sources of Funding

W.A. is supported by the Aziz Foundation, Wolfson Foundation, Royal College Surgeons of England and Oxford NIHR Biomedical Research Centre (Musculoskeletal theme). W.A was previously supported by grants from the British Association of Plastic and Reconstructive Surgeons (BAPRAS) and Royal College Surgeons of Edinburgh to complete part of this work. A.W. is an NIHR Academic Clinical Lecturer and was previously supported by an MRC Clinical Research Training Fellowship (MR/N001524/1). M.N and D.F are supported by the Oxford NIHR Biomedical Research Centre.

## Disclosures

W.W., A.A., and members of the 23andMe Research Team disclose a ‘significant’ relationship with 23andMe, Inc. as employees, and hold stock or stock options in 23andMe, Inc. All other authors have no relationships to disclose.

## Supplemental Materials (Online Only)

Supplementary Tables 1-13

Supplementary Figures 1-3

Supplementary Data 1-4

References 51-66

### 1. Supplementary Tables

**Supplementary Table 1. Codes used for varicose veins case definition in UK Biobank.** The total number of individuals with each of the codes is shown below. A total of 27,165 individuals possessed at least one of the diagnostic codes for varicose veins.

**Supplementary Table 2. Additional variants associated with varicose veins in discovery cohort.** 69 independent variants at 63 of the 109 loci were genome-wide significant (P < 5×10^−8^) in the UK Biobank cohort, however did not meet the Bonferroni-corrected threshold of P < 4.24×10^−4^ in the 23andMe replication cohort or replication data was not available (for ten tested variants). 59 of the 63 loci have not been previously reported. Using the four mapping strategies implemented in this study (see Methods), 127 genes were mapped to 51 of the 63 loci.

**Supplementary Table 3. Varicose veins associated exonic variants at the replicated loci.** 103 genome-wide significant exonic SNPs at seventeen of the replicated varicose veins susceptibility loci were identified by FUMA SNP2GENE.

**Supplementary Table 4. Predicted functional intronic and intergenic variants at the replicated loci.** 163 genome-wide significant intronic and intergenic variants predicted to be deleterious according to a CADD ≥ 12.37, as identified by FUMA SNP2GENE.

**Supplementary Table 5. Genes mapped to the varicose veins associated loci using the four mapping strategies.** 237 unique genes were mapped to 39 of 46 replicated loci by one or more gene mapping strategies (see Methods). 204 genes were mapped via positional mapping in FUMA, 80 genes were mapped via eQTL mapping in FUMA, 117 genes were mapped using MAGMA and 14 genes were mapped using summary-based mendelian randomisation. In total, 61 unique genes were mapped to novel loci. Overlap between the four different mapping strategies is shown.

**Supplementary Table 6. Genome-wide gene-based association analysis in MAGMA.** 248 protein-coding genes met the threshold for genome-wide significance (P < 2.67×10^−6^, 0.05/18,733) in this analysis. 117 of the 248 genes lay within our replicated loci and are highlighted in red.

**Supplementary Table 7. Summary-based Mendelian Randomisation (SMR) using eQTL data from GTEx v7 tibial artery.** The twenty-seven probes (genes) that met the Bonferroni-corrected significance threshold P_SMR_ < 1.01×10^−5^ (0.05/4,946) and passed the HEIDI test (P_HEIDI_ ≥ 1.12×10^−3^) (0.05/44)) are shown. Fourteen probes mapped to our replicated loci.

**Supplementary Table 8. Enriched gene sets from genome-wide gene-based enrichment analysis in MAGMA v1.07.** The convergence of 15,496 gene sets (15,381from MSigDB v7.0) were tested (See Supplementary Data 3 for all tested gene sets). A Bonferroni-corrected threshold of P < 3.23×10^−6^ (0.05/15,496) was set, resulting in four significant Gene Ontology (GO) gene sets and two curated gene sets. This analysis was performed using the SNP2GENE tool in FUMA.

**Supplementary Table 9. Genetic correlation between varicose veins and other phenotypes.** This analysis was performed using LD score (LDSC) regression, implemented in LD Hub. The traits are shown along with the consortia name, sample size, ethnicity, and PMID of the study from which the LDSC data were derived, the trait category, and the correlation coefficient (r_g_). Traits are ranked by P-value, and the twelve traits meeting a Bonferroni-corrected significant threshold of P < 5.56×10^−3^ are shown.

**Supplementary Table 10. Enriched drug pathways from the drug target enrichment analysis.** Mapped genes were interrogated with the Open Targets Platform to enrich drug pathways relating to the identified gene targets. The 200 gene targets identified by the Open Targets Platform mapped to 622 drug pathways, of which, 42 reached a nominal significance P < 0.05 and are shown below.

**Supplementary Table 11. Tractability information for targets in the drug-target enrichment analysis.** The 237 mapped genes were interrogated within the Open Targets Platform to determine their tractability to small molecule and antibody targeting. 200 genes targets were identified by the Open Targets Platform of which, tractability information was available for 105 gene targets (left column).

**Supplementary Table 12. Pharmacologically-active targets identified in drug-target enrichment** Of the 200 varicose-veins associated genes targets identified by the Open Targets Platform, eight gene targets have known pharmaceutical interactions and are presently, or in the past have been, investigated in clinical trials in different phases for the treatment of several diseases. Diseases shown below are those relating to vascular disorders.

**Supplementary Table 13. Functional categories of the gene clusters.** The varicose veins susceptibility loci map to genes implicated in five functional categories. Several genes map to more than one category. Genes not associated with varicose veins previously are highlighted in bold.

### 2. Supplementary Figures

**Supplementary Figure 1. Overview of Quality Control (QC). a**, Flowchart summarising QC protocol. Excluded SNPs are in blue panels on the left and excluded individuals are in green panels on the right. ^1^Pre-QC exclusions: 3 individuals with invalid IDs and sex, and 8 individuals who have withdrawn from UK Biobank were excluded prior to QC. ^2^Pre-association exclusions: 11 individuals who were not present in UK Biobank’s sample file accompanying the BGEN files were excluded prior to association. **b**, Principal Component Analysis (PCA) for demonstration of ethnicity of UK Biobank individuals. The UK Biobank cohort was merged with publicly available data from the 1000 Genomes Project and PCA was performed using flashpca. Individuals identified by UK Biobank as having white British ancestry are coloured in lime green, and the remaining UK Biobank individuals are in grey. In this graph of principal component 1 vs principal component 2, a near-perfect overlap can be seen between the UK Biobank “white British” individuals and both GBR (British in England and Scotland - light brown) and CEU (Utah residents with Northern and Western European ancestry - magenta) individuals from the 1000 Genomes Project.

**Supplementary Figure 2. Regional Locus Zoom plots of all varicose veins associated loci.** LocusZoom plots of the 49 independent genome-wide significant SNPs at the 46 replicated varicose veins associated susceptibility loci. Plots are ordered by chromosome number and genomic position. SNP position is shown on the x-axis, and strength of association on the y-axis (-log10 P-value). The linkage disequilibrium (LD) relationship between the lead SNP and the surrounding SNPs is indicated by the r^2^ legend. In the lower panel of each sub-figure, genes within 500kb of the index SNP are shown. The position on each chromosome is shown in relation to Human Genome build hg19.

**Supplementary Figure 3. MAGMA Gene-based association analysis Manhattan plot.** Association results for varicose veins in MAGMA gene-based association analysis. The dotted red line indicates the threshold for genome-wide significance (P < 2.68×10^−6^). 248 genes reached genome-wide significance in this analysis, with the top-ten genes highlighted in this figure.

### 3. Supplementary Data

**Supplementary Data 1. Genome-wide significant variants at all significant discovery loci.** 12,391 genome-wide significant variants (P < 5×10^−8^) at all 109 genomic risk loci identified in the discovery GWAS. Where the column headings pertain to: the SNP ID (RsID), chromosome, base position within NCBI Genome Build 37 (hg19), the effect allele, the alternate (non-effect) allele, the effect allele frequency in the study population, the imputation quality score; where 1 = genotyped SNP, the SNP effect size (BETA), the standard error of the BETA, and the GWAS association P-value. All variants are presented in ascending order according to chromosome and base position.

**Supplementary Data 2. Genome-wide significant variants at the replicated susceptibility loci.** 5,315 genome-wide significant variants (P < 5×10^−8^) identified by FUMA SNP2GENE at 45 of 46 replicated genomic risk associated with varicose veins. Where the column headings pertain to: the unique ID of the variant, the SNP ID (RsID), chromosome, base position within NCBI Genome Build 37 (hg19), the effect allele, the alternate (non-effect) allele, the effect allele frequency in the study population, the SNP effect size (BETA), the standard error of the BETA, the nearest gene according to positional mapping in FUMA, the functionality of the SNP as derived from ANNOVAR, the Combined Annotation Dependent Depletion (CADD) score of the variant, the RegulomeDB (RDB) score of the variant, the minimum chromatin state of the variant.

**Supplementary Data 3. MAGMA gene set analysis.** Gene sets were obtained from MsigDB v7.0 and total of 15,496 gene sets (Curated gene sets: 5,500, GO terms: 9,996) were tested. Curated gene sets consists of 9 data resources including KEGG, Reactome and BioCarta (http://software.broadinstitute.org/gsea/msigdb/collection_details.jsp#C2 for details). GO terms consists of three categories, biological processes (bp), cellular components (cc) and molecular functions (mf). All parameters were set as default (competitive test). Gene sets are arranged in order of ascending P-value, with a P-value < 3.23×10^−6^ indicative of a significant gene set, accounting for multiple testing (0.05/15,496).

**Supplementary Data 4. UK Biobank Summary Statistics**. The full summary statistics for the discovery GWAS of varicose veins in UK Biobank can be found on the Oxford Research Archive (ORA). URL: TBA.

### 4. Supplementary References

Supplementary References 51-66

