## Supplementary Materials for "Genome-wide association analysis and replication in 810,625 individuals identifies novel therapeutic targets for varicose veins"

### Table of Contents

#### 1. Supplementary Tables

Supplementary Table 1. Codes used for varicose veins case definition in UK Biobank

Supplementary Table 2. Additional variants associated with varicose veins in discovery cohort

Supplementary Table 3. Varicose veins associated exonic variants at the replicated loci

Supplementary Table 4. Predicted functional intronic and intergenic variants at the replicated loci

Supplementary Table 5. Genes mapped to the varicose veins associated loci using the four mapping strategies.

Supplementary Table 6. Genome-wide gene-based association analysis in MAGMA

Supplementary Table 7. Summary-based Mendelian Randomisation (SMR) using eQTL data from GTEx v7 tibial artery

Supplementary Table 8 Enriched gene sets from genome-wide gene-based enrichment analysis in MAGMA v1.07

Supplementary Table 9. Genetic correlation between varicose veins and other phenotypes

Supplementary Table 10. Enriched drug pathways from the drug target enrichment analysis

Supplementary Table 11. Tractability information for targets in the drug-target enrichment analysis

Supplementary Table 12. Pharmacologically-active targets identified in the drug-target enrichment analysis

Supplementary Table 13. Functional categories of the gene clusters

#### 2. Supplementary Figures

Supplementary Figure 1. Overview of Quality Control (QC)

Supplementary Figure 2. Regional Locus Zoom plots of all varicose veins associated loci

Supplementary Figure 3. MAGMA gene-based association analysis Manhattan plot

#### 3. Supplementary Data

Supplementary Data 1. Genome-wide significant variants at all significant discovery loci.

Supplementary Data 2. Genome-wide significant variants at the replicated susceptibility loci.

Supplementary Data 3. MAGMA gene set analysis.

Supplementary Data 4. UK Biobank Summary Statistics

#### 4. Supplementary References

Supplementary References 51-66

### 1. Supplementary Tables

**Supplementary Table 1. Codes used for varicose veins case definition in UK Biobank.** The total number of individuals with each of the diagnostic codes is shown below. A total of 27,165 individuals possessed at least one of the diagnostic codes for varicose veins.

##### Varicose veins

| Source of Data | UK Biobank Data Field | Code | Description | N |
|---|---|---|---|---|
| Primary ICD-10 | 41202 | I83 | Varicose veins of lower extremities | 12195 |
| Secondary ICD-10 | 41204 | As above | As above | 1168 |
| Primary OPCS | 41200 | L84<br>L85<br>L86<br>L87<br>L88 | Combined operations on varicose vein of leg<br>Ligation of varicose vein of leg<br>Injection into varicose vein of leg<br>Other operations on varicose vein of leg<br>Transluminal operations on varicose vein of leg | 12528 |
| Secondary OPCS | 41210 | As above | As above | 8116 |
| Non-cancer illness (self-report) | 20002 | 1494 | Varicose veins | 2266 |
| **Operation (self-report)** | 20004 | 1479 | Varicose vein surgery | 20115 |
| Total (excluding overlaps) | | | | **27165** |

**Supplementary Table 2. Additional variants associated with varicose veins in discovery cohort.** 69 independent variants at 63 of the 109 loci were genome-wide significant ($P < 5\times10^{-8}$) in the UK Biobank cohort, however did not meet the Bonferroni-corrected threshold of $P < 4.24\times10^{-4}$ in the 23andMe replication cohort or replication data was not available (for ten tested variants). 59 of the 63 loci have not been previously reported. Using the four mapping strategies implemented in this study (see Methods), 127 genes were mapped to 51 of the 63 loci.

| **SNP** | | | **Discovery GWAS in UK Biobank** | | | | | | **Replication Study in 23&Me** | | | | **Meta-Analysis** | | |
|---|---|---|---|---|---|---|---|---|---|---|---|---|---|---|---|
| **Chr** | **RsID** | **Position[a]** | **EA[b]** | **NEA[c]** | **EAF[d]** | **INFO[e]** | **OR** | **P-value** | **EAF[d]** | **INFO[e]** | **OR** | **P-value** | **OR** | **P-value** | **Mapped Genes** |
| **1** | **rs9793778** | **6554213** | **A** | **G** | **0.02** | **0.959** | **1.21 (1.13-1.29)** | **2.80x $10^{-8}$** | **0.03** | **0.849** | **1.08 (0.90-1.30)** | **4.25x $10^{-1}$** | **1.19 (1.12-1.27)** | **4.14x $10^{-8}$** | ***PLEKHG5*** |
| **1** | **rs12124756** | **8177028** | **C** | **T** | **0.46** | **0.992** | **1.07 (1.05-1.09)** | **3.10x $10^{-11}$** | **-** | **-** | **-** | **-** | **-** | **-** | ***ERRFI1, PARK7, RERE, TNFRSF9, VAMP3*** |
| **1** | **rs2046888** | **89136497** | **C** | **T** | **0.57** | **0.996** | **1.06 (1.04-1.08)** | **3.40x $10^{-10}$** | **0.59** | **0.995** | **1.00 (0.94-1.05)** | **9.02x $10^{-1}$** | **1.06 (1.04-1.07)** | **4.00x $10^{-9}$** | ***CCBL2, GBP3, GTF2B, PKN2*** |
| **1** | **rs11589479** | **155033308** | **A** | **G** | **0.16** | **G** | **1.09 (1.06-1.12)** | **1.50x $10^{-11}$** | **-** | **-** | **-** | **-** | **-** | **-** | ***ADAM15, CLK2, DCST1, EFNA3, EFNA4, FAM189B, GBA, HCN3, KRTCAP2, MTX1, MUC1, PKLR, RP11-201K10.3, THBS3*** |
| **1** | **rs551765662** | **182183969** | **G** | **T** | **0.03** | **0.976** | **1.20 (1.13-1.27)** | **1.20x $10^{-10}$** | **0.03** | **0.977** | **1.13 (0.96-1.33)** | **1.33x $10^{-1}$** | **1.19 (1.13-1.25)** | **4.81x $10^{-11}$** | ***GLUL*** |
| **1** | **rs7530318** | **185413233** | **A** | **C** | **0.66** | **0.983** | **1.06 (1.04-1.08)** | **1.50x $10^{-8}$** | **0.66** | **0.973** | **1.06 (1.00-1.12)** | **5.95x $10^{-2}$** | **1.06 (1.04-1.08)** | **2.57x $10^{-9}$** | ***HMCN1*** |
| **1** | **rs200800739** | **196813017** | **A** | **C** | **0.13** | **0.912** | **1.09 (1.06-1.13)** | **2.20x $10^{-9}$** | **-** | **-** | **-** | **-** | **-** | **-** | ***CFHR2*** |
| **2** | **rs17532799** | **15773339** | **G** | **A** | **0.25** | **0.998** | **1.06 (1.04-1.09)** | **3.60x $10^{-8}$** | **0.26** | **0.958** | **1.11 (1.04-1.18)** | **1.29x $10^{-3}$** | **1.07 (1.05-1.09)** | **3.83x $10^{-10}$** | ***DDX1*** |

| | | | | | | | | | | | | | | | |
|---|---|---|---|---|---|---|---|---|---|---|---|---|---|---|---|
| **2** | **rs4832010** | **85992648** | **T** | **C** | **0.50** | **0.994** | **1.06 (1.04-1.08)** | **$7.00\times10^{-9}$** | **0.48** | **0.973** | **1.09 (1.03-1.15)** | **$2.15\times10^{-3}$** | **1.06 (1.04-1.08)** | **$9.63\times10^{-11}$** | ***ATOH8*** |
| **2** | **rs11679159** | **218627794** | **T** | **C** | **0.07** | **0.996** | **1.11 (1.07-1.15)** | **$2.00\times10^{-8}$** | **0.07** | **0.994** | **1.10 (0.99-1.23)** | **$6.68\times10^{-2}$** | **1.11 (1.07-1.15)** | **$3.68\times10^{-9}$** | ***DIRC3*** |
| **2** | **rs61744217** | **242163827** | **T** | **C** | **0.14** | **0.975** | **1.08 (1.05-1.11)** | **$4.70\times10^{-8}$** | **0.14** | **0.943** | **1.12 (1.04-1.22)** | **$4.18\times10^{-3}$** | **1.08 (1.06-1.11)** | **$1.13\times10^{-9}$** | ***AC104841.2, ANO7, HDLBP*** |
| **3** | **rs2167037** | **8550363** | **C** | **A** | **0.47** | **0.982** | **1.06 (1.04-1.08)** | **$9.70\times10^{-9}$** | **0.48** | **0.997** | **1.07 (1.02-1.13)** | **$1.22\times10^{-2}$** | **1.06 (1.04-1.08)** | **$4.40\times10^{-10}$** | ***LMCD1*** |
| **3** | **rs322689** | **25347602** | **G** | **A** | **0.27** | **0.947** | **1.06 (1.04-1.09)** | **$4.40\times10^{-8}$** | **0.25** | **0.963** | **1.01 (0.95-1.07)** | **$7.72\times10^{-1}$** | **1.06 (1.04-1.08)** | **$1.46\times10^{-7}$** | **-** |
| **3** | **rs7614922[#]** | **128238222** | **C** | **G** | **0.14** | **0.998** | **1.13 (1.10-1.16)** | **$6.80\times10^{-19}$** | **0.15** | **0.975** | **1.10 (1.02-1.19)** | **$1.22\times10^{-2}$** | **1.13 (1.10-1.16)** | **$3.61\times10^{-20}$** | ***C3orf27, GATA2, MCM2, PLXND1, RAB7A, RPN1*** |
| **3** | **rs9842113** | **134192338** | **A** | **G** | **0.51** | **0.992** | **1.06 (1.04-1.08)** | **$1.10\times10^{-8}$** | **0.51** | **0.969** | **1.09 (1.03-1.15)** | **$3.09\times10^{-3}$** | **1.06 (1.04-1.08)** | **$2.03\times10^{-10}$** | ***ANAPC13, CEP63*** |
| **3** | **rs13322435** | **156795468** | **A** | **G** | **0.60** | **0.990** | **1.06 (1.04-1.08)** | **$8.30\times10^{-10}$** | **0.57** | **0.968** | **0.93 (0.88-0.98)** | **$6.80\times10^{-3}$** | **1.05 (1.03-1.07)** | **$9.59\times10^{-7}$** | ***TIPARP*** |
| **4** | **rs3135842** | **1796636** | **C** | **G** | **0.27** | **0.991** | **1.08 (1.05-1.10)** | **$2.50\times10^{-11}$** | **0.29** | **0.917** | **1.10 (1.03-1.17)** | **$2.51\times10^{-3}$** | **1.08 (1.06-1.10)** | **$3.15\times10^{-13}$** | ***FGFR3, LETM1, TACC3*** |
| **4** | **rs3775070** | **2663663** | **T** | **C** | **0.40** | **0.997** | **1.06 (1.04-1.08)** | **$3.50\times10^{-9}$** | **0.39** | **0.965** | **1.04 (0.98-1.10)** | **$2.11\times10^{-1}$** | **1.06 (1.04-1.08)** | **$2.15\times10^{-9}$** | ***FAM193A, RNF4, TNIP2*** |
| **4** | **rs2135479** | **125705699** | **G** | **A** | **0.67** | **0.997** | **1.06 (1.04-1.08)** | **$3.40\times10^{-8}$** | **0.66** | **0.997** | **1.04 (0.98-1.10)** | **$1.58\times10^{-1}$** | **1.06 (1.04-1.08)** | **$1.44\times10^{-8}$** | ***ANKRD50*** |
| **4** | **rs1246927** | **166543852** | **C** | **T** | **0.22** | **0.951** | **1.07 (1.04-1.09)** | **$3.60\times10^{-8}$** | **0.21** | **0.862** | **1.09 (1.01-1.17)** | **$1.99\times10^{-2}$** | **1.07 (1.05-1.09)** | **$2.53\times10^{-9}$** | **-** |

| | | | | | | | | | | | | | | | |
|---|---|---|---|---|---|---|---|---|---|---|---|---|---|---|---|
| **5** | **rs158669** | **55596849** | **A** | **G** | **0.63** | **0.995** | **1.06 (1.04-1.08)** | **$1.50 \times 10^{-8}$** | **0.62** | **0.980** | **1.03 (0.97-1.09)** | **$2.85 \times 10^{-1}$** | **1.06 (1.04-1.07)** | **$1.23 \times 10^{-8}$** | **-** |
| **5** | **rs10080063** | **79210083** | **A** | **G** | **0.57** | **0.982** | **1.06 (1.04-1.08)** | **$5.80 \times 10^{-9}$** | **0.58** | **0.981** | **1.06 (1.00-1.12)** | **$4.61 \times 10^{-2}$** | **1.06 (1.04-1.08)** | **$7.79 \times 10^{-10}$** | ***SERINC5*** |
| **6** | **rs111797764** | **12463572** | **T** | **C** | **0.05** | **G** | **1.13 (1.08-1.18)** | **$8.60 \times 10^{-9}$** | **-** | **-** | **-** | **-** | **-** | **-** | **-** |
| **6** | **rs62401797** | **46957391** | **C** | **G** | **0.92** | **0.962** | **1.12 (1.08-1.16)** | **$6.30 \times 10^{-10}$** | **-** | **-** | **-** | **-** | **-** | **-** | ***GPR110*** |
| 6 | rs440170 | 134904442 | T | C | 0.64 | 0.995 | 1.09 (1.07-1.11) | $1.50 \times 10^{-16}$ | 0.62 | 0.964 | 1.10 (1.04-1.17) | $5.70 \times 10^{-4}$ | 1.09 (1.07-1.11) | $4.64 \times 10^{-19}$ | - |
| 7 | rs9719461 | 632715 | T | C | 0.66 | 0.988 | 1.07 (1.05-1.09) | $1.80 \times 10^{-11}$ | 0.67 | 0.791 | 1.07 (0.99-1.15) | $8.32 \times 10^{-2}$ | 1.07 (1.05-1.09) | $4.15 \times 10^{-12}$ | *PRKAR1B* |
| **7** | **rs1184662** | **2758158** | **G** | **A** | **0.44** | **0.993** | **1.06 (1.04-1.08)** | **$1.20 \times 10^{-8}$** | **0.45** | **0.968** | **1.10 (1.04-1.16)** | **$6.87 \times 10^{-4}$** | **1.06 (1.04-1.08)** | **$8.70 \times 10^{-11}$** | ***AMZ1, GNA12*** |
| **7** | **rs10233387** | **27243106** | **G** | **A** | **0.55** | **0.986** | **1.06 (1.04-1.08)** | **$4.20 \times 10^{-9}$** | **0.55** | **0.954** | **1.09 (1.03-1.15)** | **$3.12 \times 10^{-3}$** | **1.06 (1.04-1.08)** | **$7.41 \times 10^{-11}$** | ***HOXA9, HOXA10, HOXA13, RP1-170O19.20*** |
| **7** | **rs62467929** | **114440716** | **G** | **A** | **0.27** | **0.992** | **1.08 (1.05-1.10)** | **$2.10 \times 10^{-11}$** | **0.24** | **0.939** | **1.10 (1.03-1.17)** | **$5.42 \times 10^{-3}$** | **1.08 (1.06-1.10)** | **$4.87 \times 10^{-13}$** | ***FOXP2, MDFIC*** |
| **7** | **rs80264508** | **129468492** | **T** | **G** | **0.91** | **0.998** | **1.11 (1.07-1.14)** | **$6.30 \times 10^{-10}$** | **0.91** | **0.995** | **1.00 (0.91-1.09)** | **$9.32 \times 10^{-1}$** | **1.09 (1.06-1.13)** | **$6.15 \times 10^{-9}$** | ***UBE2H*** |
| **8** | **rs2948294** | **8094961** | **G** | **A** | **0.44** | **0.998** | **1.05 (1.03-1.08)** | **$3.70 \times 10^{-8}$** | **0.44** | **0.816** | **0.93 (0.88-0.99)** | **$1.74 \times 10^{-2}$** | **1.04 (1.02-1.06)** | **$6.48 \times 10^{-6}$** | ***PPP1R3B*** |
| **8** | **rs6993121** | **10690124** | **C** | **T** | **0.68** | **0.995** | **1.07 (1.05-1.09)** | **$9.90 \times 10^{-12}$** | **0.67** | **0.997** | **1.02 (0.96-1.08)** | **$5.81 \times 10^{-1}$** | **1.07 (1.05-1.09)** | **$4.11 \times 10^{-11}$** | ***C8orf74, MSRA, PINX1, RP1L1, SOX7, XKR6*** |

| | | | | | | | | | | | | | | | |
|---|---|---|---|---|---|---|---|---|---|---|---|---|---|---|---|
| 8 | rs76364830 | 13372120 | G | A | 0.94 | 0.976 | 1.16 (1.12-1.21) | 1.50x $10^{-13}$ | - | - | - | - | - | - | *DLC1* |
| 8 | rs71526955 | 89604999 | C | T | 0.86 | 0.992 | 1.08 (1.05-1.11) | 2.80x $10^{-8}$ | 0.84 | 0.985 | 1.03 (0.95-1.11) | 4.80x $10^{-1}$ | 1.07 (1.05-1.10) | 4.80x $10^{-8}$ | - |
| 8 | rs7003629 | 129017012 | C | G | 0.59 | 0.995 | 1.06 (1.04-1.08) | 4.20x $10^{-9}$ | 0.60 | 0.991 | 1.07 (1.02-1.13) | 1.11x $10^{-2}$ | 1.06 (1.04-1.08) | 1.75x $10^{-10}$ | - |
| 9 | rs34919930 | 16895222 | G | A | 0.92 | 0.990 | 1.15 (1.10-1.19) | 9.70x $10^{-14}$ | 0.91 | 0.993 | 1.08 (0.98-1.19) | 1.23x $10^{-1}$ | 1.14 (1.10-1.18) | 5.79x $10^{-14}$ | *BNC2* |
| 9 | rs4978858 | 112684483 | G | T | 0.20 | 0.996 | 1.08 (1.05-1.10) | 1.60x $10^{-9}$ | 0.18 | 0.993 | 1.11 (1.03-1.19) | 3.95x $10^{-3}$ | 1.08 (1.05-1.10) | 3.19x $10^{-11}$ | *AKAP2, PALM2, PALM2-AKAP2* |
| 9 | rs72775784 | 139406496 | T | C | 0.08 | 0.999 | 1.12 (1.08-1.16) | 3.30x $10^{-10}$ | 0.08 | 0.879 | 1.10 (0.99-1.23) | 7.13x $10^{-2}$ | 1.12 (1.08-1.16) | 6.67x $10^{-11}$ | *NOTCH1* |
| 10 | rs7893671 | 5524502 | A | C | 0.25 | 0.991 | 1.07 (1.05-1.10) | 5.90x $10^{-10}$ | 0.24 | 0.993 | 1.10 (1.04-1.17) | 2.17x $10^{-3}$ | 1.07 (1.05-1.10) | 7.24x $10^{-12}$ | *CALML5, NET1, TUBAL3* |
| 10 | rs2474723 | 33485962 | T | C | 0.75 | 0.986 | 1.07 (1.04-1.09) | 1.00x $10^{-8}$ | 0.73 | 0.985 | 1.06 (1.00-1.13) | 4.73x $10^{-2}$ | 1.07 (1.04-1.09) | 1.43x $10^{-9}$ | *NRP1* |
| 10 | rs10761604 | 63819903 | C | T | 0.57 | 0.982 | 1.06 (1.04-1.08) | 3.00x $10^{-10}$ | 0.55 | 0.973 | 1.10 (1.04-1.16) | 5.74x $10^{-4}$ | 1.07 (1.05-1.09) | 1.46x $10^{-12}$ | *ARID5B* |
| 10 | rs4415677 | 73039860 | T | C | 0.78 | 0.994 | 1.07 (1.05-1.10) | 8.10x $10^{-10}$ | 0.77 | 0.994 | 1.01 (0.95-1.08) | 7.52x $10^{-1}$ | 1.07 (1.04-1.09) | 3.82x $10^{-9}$ | *UNC5B* |
| 10 | 10:79677281_CA_C | 73039860 | CA | C | 0.73 | 0.967 | 1.06 (1.04-1.09) | 1.30x $10^{-8}$ | - | - | - | - | - | - | - |
| 11 | rs11603111 | 1568192 | C | A | 0.45 | 0.998 | 1.07 (1.05-1.09) | 4.90x $10^{-12}$ | 0.47 | 0.987 | 1.07 (1.02-1.13) | 8.96x $10^{-3}$ | 1.07 (1.05-1.09) | 1.59x $10^{-13}$ | *BRSK2, DUSP8, KRTAP5-1, KRTAP5-2, KRTAP5-3, KRTAP5-5, KRTAP5-AS1, MOB2* |

| | | | | | | | | | | | | | | | |
|---|---|---|---|---|---|---|---|---|---|---|---|---|---|---|---|
| 11 | rs7111987 | 10353717 | A | G | 0.36 | 0.996 | 1.09 (1.07-1.11) | $1.10 \times 10^{-16}$ | - | - | - | - | - | - | *ADM, AMPD3, SBF2, SWAP70* |
| 12 | rs2159408 | 3202368 | A | C | 0.51 | 0.996 | 1.07 (1.05-1.09) | $4.50 \times 10^{-11}$ | 0.52 | 0.965 | 1.09 (1.03-1.15) | $1.64 \times 10^{-3}$ | 1.07 (1.05-1.09) | $4.24 \times 10^{-13}$ | *TSPAN9* |
| 12 | rs11168250 | 48209165 | T | G | 0.24 | G | 1.10 (1.08-1.13) | $2.40 \times 10^{-18}$ | 0.23 | 0.962 | 1.11 (1.04-1.19) | $1.09 \times 10^{-3}$ | 1.10 (1.08-1.13) | $1.18 \times 10^{-20}$ | *AC004466.1, ENDOU, HDAC7, RAPGEF3, RP1-228P16.5, SLC48A1* |
| 13 | rs7331874 | 38114420 | T | G | 0.51 | 0.995 | 1.06 (1.04-1.08) | $7.90 \times 10^{-10}$ | 0.52 | 0.985 | 1.02 (0.96-1.07) | $5.07 \times 10^{-1}$ | 1.06 (1.04-1.08) | $1.80 \times 10^{-9}$ | - |
| 13 | rs8002731 | 40362229 | C | A | 0.35 | 0.996 | 1.06 (1.04-1.08) | $3.90 \times 10^{-9}$ | 0.34 | 0.986 | 1.06 (1.01-1.13) | $3.22 \times 10^{-2}$ | 1.06 (1.04-1.08) | $3.87 \times 10^{-10}$ | *COG6* |
| 13 | rs1549062 | 106264061 | A | G | 0.36 | 0.999 | 1.09 (1.07-1.11) | $3.60 \times 10^{-18}$ | 0.37 | 0.793 | 1.04 (0.98-1.10) | $2.44 \times 10^{-1}$ | 1.09 (1.07-1.11) | $6.89 \times 10^{-18}$ | - |
| 14 | rs2899888 | 39889759 | C | T | 0.24 | 0.990 | 1.07 (1.05-1.10) | $9.10 \times 10^{-10}$ | 0.26 | 0.978 | 1.01 (0.95-1.08) | $6.64 \times 10^{-1}$ | 1.07 (1.04-1.09) | $3.37 \times 10^{-9}$ | *CTAGE5, FBXO33, GEMIN2, MIA2, RP11-407N17.3* |
| 15 | rs36160775 | 86167378 | G | A | 0.78 | 0.996 | 1.07 (1.04-1.09) | $3.70 \times 10^{-8}$ | 0.79 | 0.989 | 1.06 (0.99-1.13) | $9.83 \times 10^{-2}$ | 1.07 (1.04-1.09) | $9.50 \times 10^{-9}$ | *AKAP13* |
| 15 | rs10520785# | 96017545 | T | C | 0.75 | G | 1.07 (1.04-1.09) | $3.70 \times 10^{-9}$ | 0.74 | 0.977 | 1.07 (1.01-1.14) | $3.04 \times 10^{-2}$ | 1.07 (1.05-1.09) | $3.45 \times 10^{-10}$ | - |
| 15 | rs7167309 | 100003407 | T | G | 0.70 | 0.995 | 1.06 (1.04-1.09) | $3.40 \times 10^{-9}$ | 0.70 | 0.991 | 1.10 (1.04-1.17) | $1.18 \times 10^{-3}$ | 1.07 (1.05-1.09) | $2.95 \times 10^{-11}$ | *MEF2A* |
| 15 | rs1626645 | 101244572 | C | T | 0.22 | 0.989 | 1.07 (1.04-1.09) | $1.90 \times 10^{-8}$ | 0.21 | 0.956 | 1.04 (0.98-1.11) | $2.14 \times 10^{-1}$ | 1.07 (1.04-1.09) | $1.10 \times 10^{-8}$ | - |
| 17 | rs12451961 | 7351479 | A | G | 0.64 | 0.996 | 1.06 (1.04-1.08) | $1.50 \times 10^{-8}$ | 0.64 | 0.986 | 1.07 (1.01-1.13) | $1.78 \times 10^{-2}$ | 1.06 (1.04-1.08) | $9.25 \times 10^{-10}$ | *CHRNB1, FGF11, POLR2A, SLC35G6, ZBTB4* |

| | | | | | | | | | | | | | | | |
|---|---|---|---|---|---|---|---|---|---|---|---|---|---|---|---|
| 17 | rs2656417 | 18475630 | T | C | 0.53 | 0.953 | 1.06 (1.04-1.08) | 4.90x 10$^{-9}$ | - | - | - | - | - | - | *CTD-2303H24.2, FAM106A, SHMT1, TBC1D28, TVP23B, USP32P2, ZNF286B* |
| 17 | rs7209761 | 21207916 | A | G | 0.51 | 0.919 | 1.08 (1.06-1.10) | 5.50x 10$^{-15}$ | - | - | - | - | - | - | *KCNJ12, MAP2K3* |
| 17 | rs4968801# | 68159800 | A | G | 0.89 | 0.998 | 1.11 (1.08-1.15) | 1.10x 10$^{-12}$ | 0.89 | 0.977 | 1.15 (1.05-1.25) | 1.53x 10$^{-3}$ | 1.12 (1.09-1.15) | 8.60x 10$^{-15}$ | *KCNJ2, KCNJ16* |
| 18 | rs390439 | 77249122 | T | C | 0.51 | 0.999 | 1.05 (1.03-1.07) | 4.90x 10$^{-8}$ | 0.53 | 0.924 | 0.99 (0.93-1.04) | 6.21x 10$^{-1}$ | 1.05 (1.03-1.07) | 5.75x 10$^{-7}$ | *NFATC1* |
| 19 | rs451367* | 16412842 | T | C | 0.20 | 0.990 | 1.08 (1.05-1.11) | 2.10x 10$^{-10}$ | 0.19 | 0.864 | 1.03 (0.95-1.11) | 4.69x 10$^{-1}$ | 1.07 (1.05-1.10) | 3.64x 10$^{-10}$ | *AP1M1, FAM32A, KLF2* |
| 19 | rs76378167 | 18433714 | G | A | 0.94 | G | 1.12 (1.07-1.16) | 4.30x 10$^{-8}$ | 0.94 | 0.912 | 1.21 (1.08-1.36) | 8.97x 10$^{-4}$ | 1.13 (1.09-1.17) | 4.00x 10$^{-10}$ | *LSM4, PGPEP1* |
| 19 | rs12975407# | 18443650 | C | T | 0.45 | 0.993 | 1.04 (1.02-1.06) | 1.40x 10$^{-5}$ | 0.46 | 0.963 | 1.05 (1.00-1.11) | 5.48x 10$^{-2}$ | 1.04 (1.03-1.06) | 2.18x 10$^{-6}$ | *LSM4, PGPEP1* |
| 19 | rs4805891 | 33970989 | T | C | 0.65 | 0.992 | 1.06 (1.04-1.08) | 4.30x 10$^{-8}$ | 0.64 | 0.987 | 1.10 (1.04-1.16) | 1.01x 10$^{-3}$ | 1.06 (1.04-1.08) | 4.00x 10$^{-10}$ | *PEPD* |
| 20 | rs2235856# | 50158473 | A | C | 0.64 | G | 1.01 (0.99-1.03) | 1.60x 10$^{-1}$ | 0.64 | 0.961 | 1.07 (1.01-1.13) | 2.47x 10$^{-2}$ | 1.02 (1.00-1.04) | 4.04x 10$^{-2}$ | *NFATC2* |
| 21 | rs2836400 | 39815212 | C | T | 0.49 | 0.987 | 1.06 (1.04-1.08) | 1.30x 10$^{-9}$ | 0.51 | 0.963 | 1.09 (1.03-1.15) | 2.11x 10$^{-3}$ | 1.06 (1.04-1.08) | 1.64x 10$^{-11}$ | *ERG* |
| 22 | rs176155 | 24499204 | A | G | 0.15 | 0.999 | 1.09 (1.06-1.12) | 3.10x 10$^{-10}$ | 0.16 | 0.955 | 1.13 (1.05-1.21) | 1.88x 10$^{-3}$ | 1.09 (1.07-1.12) | 3.31x 10$^{-12}$ | *CABIN1, SUSD2* |
| 22 | rs13058589 | 39646737 | G | T | 0.09 | G | 1.13 (1.09-1.17) | 3.80x 10$^{-12}$ | 0.08 | 0.833 | 1.19 (1.07-1.32) | 1.08x 10$^{-3}$ | 1.13 (1.10-1.17) | 2.78x 10$^{-14}$ | *AL031590.1, PDGFB* |

| | | | | | | | | | | | | | | |
|---|---|---|---|---|---|---|---|---|---|---|---|---|---|---|
| **22** | **rs13054155** | **51118478** | **T** | **C** | **0.51** | **0.986** | **1.07 (1.05-1.09)** | **8.70x $10^{-13}$** | **0.49** | **0.968** | **1.08 (1.02-1.14)** | **6.06x $10^{-3}$** | **1.07 (1.05-1.09)** | **2.02x $10^{-14}$** | ***PLXNB2, SHANK3*** |

[a]Based on NCBI Genome Build 37 (hg19). [b]The effect allele. [c]The alternate (non-effect) allele. [d]The effect allele frequency in the study population. [e]The imputation quality score; G= genotyped SNP. [f]Odds ratio (95% confidence intervals). OR > 1 indicative of increased risk with effect allele. [#]denotes five additional residual significant signals following conditional regression analysis at the lead SNP at the locus. *At this locus, 19p13.11, the lead SNP in the UK Biobank cohort, rs451367, did not replicate in the 23andMe cohort, however, an independent residual signal at the locus, rs12609241($P_{discovery}$ = 3.00x$10^{-10}$, $P_{replication}$ = 1.84x$10^{-18}$, $P_{meta}$ = 1.32x$10^{-18}$ did replicate in the 23andMe Cohort (Table 1). Bold variants represent loci not previously reported.

**Supplementary Table 3. Varicose veins associated exonic variants at the replicated loci.** 103 genome-wide significant exonic SNPs at seventeen of the replicated varicose veins susceptibility loci were identified by FUMA SNP2GENE.

| rsID | Chr | Pos[a] | EA[b] | NEA[c] | EAF[d] | P-Value | OR (95% CI)[e] | Index SNP[f] | $r^2$ [g] | D' [g] | Nearest Gene[h] | CADD | RDB | Functionality[i] | HGVSp[j] | GERP[k] | Polyphen[l] | SIFT[l] |
|---|---|---|---|---|---|---|---|---|---|---|---|---|---|---|---|---|---|---|
| rs5821892 | 17 | 70036479 | C | CG | 0.41 | $3.00x10^{-21}$ | 1.10 (1.08-1.12) | rs9895127 | 0.93 | 1.00 | RP11-84E24.2:SOX9-AS1:AC007461.1 | 17.15 | NA | Frameshift | p.Pro6AlafsTer31 | - | - | - |
| rs10876023 | 12 | 50746917 | A | G | 0.35 | $1.60x10^{-9}$ | 1.06 (1.04-1.08) | rs7308356 | 0.26 | 0.91 | FAM186A | 1.80 | 6 | Missense | p.Leu1233Pro | -1.54 | Benign | Tolerated |
| rs10876024 | 12 | 50747005 | T | C | 0.35 | $1.60x10^{-9}$ | 1.06 (1.04-1.08) | rs7308356 | 0.27 | 0.92 | FAM186A | 9.73 | 6 | Missense | p.Arg1204Gly | -3 | Benign | Tolerated |
| rs11169360 | 12 | 50691474 | T | A | 0.35 | $2.10x10^{-9}$ | 0.94 (0.92-0.96) | rs7308356 | 0.27 | 0.92 | AC140061.12 | 8.58 | 6 | Missense | p.Phe42Tyr | -0.63 | Unknown | - |
| rs11549835 | 16 | 88781073 | G | A | 0.15 | $5.20x10^{-31}$ | 1.17 (1.14-1.20) | rs2002833 | 0.04 | 0.60 | CTU2 | 9.48 | NA | Missense | p.Arg427Gln | -2.64 | Benign | Tolerated |
| rs11549837 | 16 | 88779739 | A | G | 0.33 | $4.10x10^{-22}$ | 1.10 (1.08-1.13) | rs2002833 | 0.08 | 0.57 | CTU2 | 17.65 | 4 | Missense | p.Met253Val | 0.02 | Benign | Tolerated |
| **rs12303082**# | 12 | 50754563 | T | G | 0.35 | $2.00x10^{-9}$ | 1.06 (1.04-1.08) | rs7308356 | 0.27 | 0.92 | FAM186A | 21.20 | 6 | Missense | p.Lys187Gln | 0.83 | Probably damaging | Deleterious |
| rs12924185 | 16 | 89016884 | C | T | 0.36 | $4.50x10^{-13}$ | 1.08 (1.05-1.10) | rs2002833 | 0.02 | 0.23 | CBFA2T3:RP11-830F9.6 | 12.67 | 6 | Missense | p.Arg120Trp | - | Benign | Tolerated Low Confidence |
| rs13009282 | 2 | 68364478 | T | C | 0.27 | $1.50x10^{-14}$ | 0.92 (0.90-0.94) | rs2861819 | 0.22 | 1.00 | WDR92:RP11-474G23.1 | 20.20 | 3a | Missense | p.Met241Val | 2.96 | Benign | Tolerated |
| rs13195509 | 6 | 26463660 | G | A | 0.12 | $1.70x10^{-8}$ | 1.09 (1.06-1.12) | rs7773004 | 0.08 | 0.81 | BTN2A1 | 22.50 | 1f | Missense | p.Val207Met | - | Possibly damaging | Tolerated |

| | | | | | | | | | | | | | | | | | | |
|---|---|---|---|---|---|---|---|---|---|---|---|---|---|---|---|---|---|---|
| rs139332558 | 6 | 26637724 | T | C | 0.10 | 2.30x $10^{-9}$ | 1.10 (1.07-1.13) | rs7773004 | 0.07 | 0.78 | ZNF322 | 19.82 | 5 | Missense | - | - | - | - |
| rs16891235 | 6 | 26017542 | T | C | 0.12 | 4.00x $10^{-11}$ | 1.10 (1.07-1.13) | rs7773004 | 0.08 | 0.75 | HIST1H1A | 11.55 | 1f | Missense | p.Lys140Arg | - | Unknown | Tolerated |
| rs198845 | 6 | 26107790 | G | T | 0.37 | 7.70x $10^{-9}$ | 0.94 (0.93-0.96) | rs7773004 | 0.13 | 0.55 | HIST1H1T | 14.07 | 6 | Missense | p.Gln178Lys | 0.33 | Benign | Tolerated |
| rs200484 | 6 | 27775674 | A | G | 0.13 | 2.50x $10^{-9}$ | 1.09 (1.06-1.12) | rs7773004 | 0.05 | 0.67 | HIST1H2BL | 16.23 | 1f | Missense | Leu4Pro | 0.67 | Benign | Tolerated Low Confidence |
| rs200833298 | 16 | 89017386 | T | C | 0.05 | 4.30x $10^{-9}$ | 1.13 (1.09-1.18) | rs2002833 | 0.02 | 1.00 | CBFA2T3:RP11-830F9.6 | 13.20 | 6 | Missense | p.Val287Ala | - | Unknown | Deleterious Low Confidence |
| rs2044693 | 2 | 68385097 | A | G | 0.37 | 8.10x $10^{-53}$ | 0.86 (0.84-0.87) | rs2861819 | 0.91 | 0.98 | RP11-474G23.1:PNO1 | 14.88 | NA | Missense | p.Arg11Gly | 1.74 | Benign | Tolerated |
| rs2071790 | 6 | 28911802 | G | A | 0.49 | 3.20x $10^{-10}$ | 1.06 (1.04-1.08) | rs7773004 | 0.04 | 0.21 | C6orf100 | 6.34 | NA | Missense | p.Gly41Glu | -2.59 | Benign | Tolerated Low Confidence |
| rs2073152 | 6 | 29364838 | C | G | 0.29 | 3.60x $10^{-8}$ | 0.94 (0.92-0.96) | rs7773004 | 0.01 | 0.14 | OR5V1:OR12D2 | 13.64 | 6 | Missense | p.Ser121Cys | -3.33 | Benign | Deleterious |
| rs2232423 | 6 | 28366151 | A | G | 0.11 | 1.60x $10^{-9}$ | 1.09 (1.06-1.13) | rs7773004 | 0.06 | 0.77 | ZSCAN12 | 0.36 | NA | Missense | p.Met11Thr | - | Benign | Tolerated |
| rs2269700 | 9 | 118163563 | C | T | 0.39 | 4.40x $10^{-15}$ | 1.08 (1.06-1.10) | rs10817762 | 0.59 | 1.00 | DEC1 | 0.51 | 5 | Missense | p.Ala60Val | -2.7 | Benign | Tolerated Low Confidence |
| rs2304789 | 8 | 87567193 | C | T | 0.33 | 2.70x $10^{-11}$ | 1.07 (1.05-1.09) | rs10504825 | 0.32 | 1.00 | CPNE3:CNGB3 | 23.20 | NA | Missense | p.Thr412Met | - | Benign | Tolerated |
| **rs2342302#** | 17 | 70036546 | G | A | 0.40 | 1.30x $10^{-21}$ | 1.10 (1.08-1.12) | rs9895127 | 0.87 | 1.00 | RP11-84E24.2:SOX9- | 14.87 | 6 | Missense | p.Gly28Arg | - | Probably damaging | - |

| | | | | | | | | | | | | | | | | | | |
|---|---|---|---|---|---|---|---|---|---|---|---|---|---|---|---|---|---|---|
| | | | | | | | | | | | AS1:AC007461.1 | | | | | | | |
| rs3116855 | 6 | 29141632 | T | C | 0.49 | $1.40 \times 10^{-8}$ | 0.95 (0.93-0.96) | rs7773004 | 0.03 | 0.18 | OR2J2 | 23.10 | 7 | Missense | p.Tyr74His | - | Probably damaging | Deleterious |
| rs3130743 | 6 | 29142064 | A | G | 0.49 | $1.50 \times 10^{-8}$ | 0.95 (0.93-0.96) | rs7773004 | 0.03 | 0.18 | OR2J2 | 15.59 | 7 | Missense | p.Thr218Ala | - | Benign | Deleterious |
| rs33932084 | 6 | 28268824 | A | G | 0.11 | $8.50 \times 10^{-10}$ | 1.10 (1.06-1.13) | rs7773004 | 0.06 | 0.77 | PGBD1 | 14.75 | 3a | Missense | p.Asn398Ser | -1.27 | Benign | Tolerated |
| rs34525648 | 6 | 25914853 | G | A | 0.12 | $3.50 \times 10^{-10}$ | 1.10 (1.07-1.13) | rs7773004 | 0.08 | 0.75 | SLC17A2 | 15.35 | 6 | Missense | p.Ser370Leu | -3.23 | Possibly damaging | Tolerated |
| rs34788973 | 6 | 27879200 | C | A | 0.11 | $1.20 \times 10^{-9}$ | 1.10 (1.06-1.13) | rs7773004 | 0.05 | 0.67 | OR2B2 | 23.20 | 7 | Missense | Ala300Ser | 2.26 | Benign | Deleterious |
| rs35265318 | 16 | 88800785 | C | T | 0.14 | $1.50 \times 10^{-31}$ | 1.17 (1.14-1.21) | rs2002833 | 0.05 | 0.76 | PIEZO1:RP5-1142A6.2 | 9.23 | NA | Missense | p.Arg720His | -1.98 | Benign | Tolerated |
| rs35555795 | 6 | 26509382 | C | T | 0.12 | $3.50 \times 10^{-8}$ | 1.08 (1.05-1.12) | rs7773004 | 0.08 | 0.80 | BTN1A1 | 0.92 | 7 | Missense | p.Pro521Ser | -4.58 | Benign | Tolerated |
| rs3734542 | 6 | 26468326 | G | A | 0.12 | $3.40 \times 10^{-8}$ | 1.08 (1.05-1.12) | rs7773004 | 0.08 | 0.81 | BTN2A1 | 5.38 | 5 | Missense | p.Arg378Gln | -4.39 | Benign | Tolerated |
| rs3734543 | 6 | 26468545 | G | C | 0.12 | $2.80 \times 10^{-8}$ | 1.09 (1.05-1.12) | rs7773004 | 0.08 | 0.81 | BTN2A1 | 11.89 | 5 | Missense | p.Gly451Ala | -2.86 | Benign | Tolerated |
| rs3745318 | 19 | 16436262 | T | C | 0.25 | $2.50 \times 10^{-8}$ | 1.06 (1.04-1.09) | rs12609241 | 0.01 | 0.21 | KLF2 | 15.95 | 4 | Missense | p.Leu104Pro | -3.45 | Benign | Tolerated |
| rs3749971 | 6 | 29342775 | G | A | 0.12 | $9.10 \times 10^{-10}$ | 1.09 (1.06-1.12) | rs7773004 | 0.06 | 0.76 | OR5V1:OR12D3 | 11.39 | NA | Missense | p.Thr97Ile | -1.48 | Benign | Tolerated |

| | | | | | | | | | | | | | | | | | | |
|---|---|---|---|---|---|---|---|---|---|---|---|---|---|---|---|---|---|---|
| rs3796529 | 4 | 57797414 | C | T | 0.19 | $1.90 \times 10^{-12}$ | 0.92 (0.89-0.94) | rs56155140 | 1.00 | 1.00 | REST | 3.93 | NA | Missense | p.Pro797Leu | -2.68 | Benign | Tolerated Low Confidence |
| rs41266839 | 6 | 26409890 | G | C | 0.10 | $2.90 \times 10^{-9}$ | 1.10 (1.06-1.13) | rs7773004 | 0.10 | 1.00 | BTN3A1 | 0.50 | 4 | Missense | p.Arg282Thr | - | Benign | Tolerated |
| rs4782321 | 16 | 88780175 | G | A | 0.15 | $1.50 \times 10^{-31}$ | 1.17 (1.14-1.20) | rs2002833 | 0.05 | 0.69 | CTU2 | 3.72 | 4 | Missense | p.Val332Ile | -7.01 | Benign | Tolerated |
| rs540473 | 9 | 214706 | G | C | 0.42 | $6.60 \times 10^{-9}$ | 0.94 (0.93-0.96) | rs78216177 | 0.16 | 0.81 | C9orf66 | 10.06 | 4 | Missense | p.Arg231Gly | - | Benign | Tolerated Low Confidence |
| rs61734177 | 16 | 88964557 | C | G | 0.09 | $2.80 \times 10^{-9}$ | 1.10 (1.07-1.14) | rs2002833 | 0.01 | 0.50 | CBFA2T3 | 21.90 | NA | Missense | p.Arg103Pro | - | Benign | Tolerated Low Confidence |
| rs61742093 | 6 | 27879982 | A | G | 0.11 | $7.00 \times 10^{-10}$ | 1.10 (1.07-1.13) | rs7773004 | 0.05 | 0.67 | OR2B2 | 22.60 | 3a | Missense | p.Ile39Thr | - | Benign | Deleterious |
| rs62070084 | 17 | 70036573 | A | G | 0.27 | $1.40 \times 10^{-9}$ | 1.07 (1.05-1.09) | rs9895127 | 0.01 | 0.94 | RP11-84E24.2:SOX9-AS1:AC007461.1 | 15.14 | 7 | Missense | p.Asn37Asp | | Possibly damaging | - |
| rs6500486 | 16 | 88723629 | C | T | 0.13 | $2.00 \times 10^{-29}$ | 0.85 (0.83-0.88) | rs2002833 | 0.12 | 0.70 | MVD | 2.91 | 5 | Missense | p.Val144Ile | - | Unknown | Tolerated Low Confidence |
| rs6500493 | 16 | 88803982 | C | G | 0.33 | $1.30 \times 10^{-8}$ | 1.06 (1.04-1.08) | rs2002833 | 0.00 | 0.04 | PIEZO1:RP5-1142A6.2 | 0.54 | 1f | Missense | p.Val394Leu | -6.73 | Benign | Tolerated |
| rs6500495 | 16 | 88808743 | A | G | 0.13 | $1.00 \times 10^{-43}$ | 1.22 (1.19-1.26) | rs2002833 | 0.25 | 0.84 | PIEZO1:RP5-1142A6.7 | 14.04 | NA | Missense | p.Ile83Thr | 1.36 | Benign | Tolerated |
| **rs6580741**[#] | 12 | 50727706 | G | C | 0.35 | $2.50 \times 10^{-9}$ | 0.94 (0.92-0.96) | rs7308356 | 0.27 | 0.92 | FAM186A | 0.26 | 5 | Missense | p.His61Gln | - | Possibly damaging | Tolerated Low Confidence |
| rs706792 | 12 | 50467644 | G | T | 0.41 | $1.50 \times 10^{-12}$ | 1.07 (1.05-1.09) | rs7308356 | 0.71 | 0.90 | ASIC1 | 19.00 | NA | Missense | p.Ala93Ser | 2.8 | Benign | Tolerated |

| | | | | | | | | | | | | | | | | | | |
|---|---|---|---|---|---|---|---|---|---|---|---|---|---|---|---|---|---|---|
| **rs7184427**# | 16 | 88804734 | A | G | 0.15 | $9.10 \times 10^{-40}$ | 1.19 (1.16-1.23) | rs2002833 | 0.22 | 0.75 | PIEZO1:RP5-1142A6.2 | 17.39 | 5 | Missense | p.Val250Ala | 0.49 | Benign | Deleterious |
| rs7296291 | 12 | 50744119 | G | A | 0.35 | $1.50 \times 10^{-9}$ | 1.06 (1.04-1.08) | rs7308356 | 0.27 | 0.92 | FAM186A | 10.64 | 6 | Missense | p.His2166Tyr | - | Benign | Tolerated |
| rs7302981 | 12 | 50537815 | A | G | 0.37 | $5.90 \times 10^{-16}$ | 0.92 (0.90-0.94) | rs7308356 | 1.00 | 1.00 | RP4-605O3.4:CERS5 | 16.30 | 6 | Missense | p.Cys75Arg | -1.47 | Benign | Tolerated |
| rs7404939 | 16 | 88807896 | G | A | 0.12 | $1.20 \times 10^{-38}$ | 0.82 (0.80-0.85) | rs2002833 | 0.23 | 0.83 | PIEZO1 | 9.01 | 5 | Missense | p.Pro152Leu | -0.09 | Benign | Tolerated |
| rs76267236 | 16 | 88931274 | C | T | 0.09 | $1.40 \times 10^{-12}$ | 1.13 (1.09-1.17) | rs2002833 | 0.04 | 1.00 | PABPN1L | 24.10 | 5 | Missense | p.Gly214Asp | - | Benign | Tolerated |
| rs7766641 | 6 | 26184102 | G | A | 0.26 | $4.10 \times 10^{-9}$ | 1.07 (1.04-1.09) | rs7773004 | 0.31 | 0.85 | HIST1H2BE | 22.40 | 4 | Missense | p.Gly27Ser | 1.65 | Unknown | Tolerated Low Confidence |
| rs8337 | 6 | 29523676 | C | G | 0.27 | $6.00 \times 10^{-9}$ | 1.07 (1.04-1.09) | rs7773004 | - | - | UBD:GABBR1 | 0.01 | 5 | Missense | p.Cys160Ser | -4.12 | Benign | Tolerated |
| rs853684 | 6 | 28294550 | T | C | 0.39 | $1.20 \times 10^{-9}$ | 1.06 (1.04-1.08) | rs7773004 | 0.07 | 0.34 | ZSCAN31 | 0.18 | NA | Missense | p.Lys205Arg | - | Benign | Tolerated |
| rs1129406 | 12 | 51203371 | C | T | 0.41 | $1.70 \times 10^{-13}$ | 0.93 (0.91-0.95) | rs7308356 | 0.28 | 0.85 | ATF1 | 20.70 | 5 | Splice Region | c.327C>T | 2.59 | - | - |
| rs13195401 | 6 | 26463574 | G | T | 0.10 | $3.80 \times 10^{-9}$ | 1.09 (1.06-1.13) | rs7773004 | 0.07 | 0.79 | BTN2A1 | 22.80 | 5 | Stop gained | p.Trp178Leu | - | Probably damaging | Deleterious |
| rs13195402 | 6 | 26463575 | G | T | 0.11 | $6.40 \times 10^{-9}$ | 1.09 (1.06-1.13) | rs7773004 | 0.07 | 0.79 | BTN2A1 | 23.70 | 5 | Stop gained | p.Trp178Cys | - | Probably damaging | Deleterious |
| rs10506292 | 12 | 50744753 | T | C | 0.35 | $1.60 \times 10^{-9}$ | 1.06 (1.04-1.08) | rs7308356 | 0.27 | 0.92 | FAM186A | 0.81 | 1f | Synonymous | - | - | - | - |

| | | | | | | | | | | | | | | | | | | |
|---|---|---|---|---|---|---|---|---|---|---|---|---|---|---|---|---|---|---|
| rs10776795 | 1 | 115604798 | C | T | 0.29 | $3.20\times 10^{-10}$ | 0.94 (0.92-0.96) | rs7518191 | 1.00 | 1.00 | TSPAN2 | 10.45 | 4 | Synonymous | - | - | - | - |
| rs11169524 | 12 | 51089734 | T | A | 0.36 | $4.30\times 10^{-11}$ | 0.94 (0.92-0.95) | rs7308356 | 0.22 | 0.83 | DIP2B | 11.43 | 6 | Synonymous | - | - | - | - |
| rs1137930 | 2 | 68388823 | A | G | 0.29 | $2.80\times 10^{-12}$ | 0.93 (0.91-0.95) | rs2861819 | 0.23 | 0.95 | RP11-474G23.1:PNO1 | 19.09 | 7 | Synonymous | - | - | - | - |
| rs1368298 | 5 | 158204425 | A | G | 0.46 | $8.70\times 10^{-32}$ | 0.89 (0.88-0.91) | rs11135046 | 0.94 | 0.98 | EBF1 | 9.25 | NA | Synonymous | - | - | - | - |
| rs1697 | 3 | 128356852 | C | T | 0.39 | $4.60\times 10^{-12}$ | 0.93 (0.92-0.95) | rs2713575 | 0.45 | 0.81 | RPN1 | 3.49 | NA | Synonymous | - | - | - | - |
| rs173135 | 17 | 68172326 | C | T | 0.11 | $5.80\times 10^{-11}$ | 1.10 (1.07-1.14) | rs638538 | 0.01 | 0.41 | KCNJ2 | 10.52 | NA | Synonymous | - | - | - | - |
| rs17528178 | 6 | 25983777 | C | T | 0.09 | $3.50\times 10^{-9}$ | 1.10 (1.07-1.13) | rs7773004 | 0.09 | 0.89 | TRIM38 | 0.63 | NA | Synonymous | - | - | - | - |
| rs17763089 | 6 | 27835218 | G | A | 0.11 | $3.80\times 10^{-9}$ | 1.09 (1.06-1.12) | rs7773004 | 0.05 | 0.67 | HIST1H1B | 7.41 | 4 | Synonymous | - | - | - | - |
| rs17774663 | 6 | 28120898 | G | A | 0.24 | $3.80\times 10^{-10}$ | 1.07 (1.05-1.10) | rs7773004 | 0.10 | 0.69 | ZKSCAN8 | 0.10 | 6 | Synonymous | - | - | - | - |
| rs1892251 | 6 | 25769349 | T | C | 0.12 | $3.40\times 10^{-9}$ | 0.92 (0.89-0.94) | rs7773004 | 0.08 | 0.75 | SLC17A4 | 5.92 | 7 | Synonymous | - | - | - | - |
| rs198852 | 6 | 26104448 | A | G | 0.37 | $4.50\times 10^{-9}$ | 0.94 (0.93-0.96) | rs7773004 | 0.13 | 0.55 | HIST1H4C | 14.95 | 4 | Synonymous | - | - | - | - |
| rs200948 | 6 | 27835272 | T | C | 0.13 | $1.80\times 10^{-9}$ | 1.09 (1.06-1.12) | rs7773004 | 0.05 | 0.67 | HIST1H1B | 12.12 | 1f | Synonymous | - | - | - | - |

| | | | | | | | | | | | | | | | | | | |
|---|---|---|---|---|---|---|---|---|---|---|---|---|---|---|---|---|---|---|
| rs200973 | 6 | 27858421 | A | G | 0.19 | $6.90 \times 10^{-13}$ | 1.09 (1.07-1.12) | rs7773004 | 0.07 | 0.61 | HIST1H3J | 21.50 | 1b | Synonymous | - | - | - | - |
| rs200981 | 6 | 27833174 | A | G | 0.13 | $1.70 \times 10^{-9}$ | 1.09 (1.06-1.12) | rs7773004 | 0.06 | 0.69 | HIST1H2AL | 15.90 | NA | Synonymous | - | - | - | - |
| rs2227901 | 4 | 57798189 | G | A | 0.19 | $2.10 \times 10^{-12}$ | 0.92 (0.89-0.94) | rs56155140 | 1.00 | 1.00 | REST | 0.35 | 1f | Synonymous | - | - | - | - |
| rs2230683 | 6 | 28891176 | T | C | 0.11 | $7.80 \times 10^{-10}$ | 1.10 (1.06-1.13) | rs7773004 | 0.06 | 0.76 | TRIM27 | 18.69 | NA | Synonymous | - | - | - | - |
| rs2279258 | 16 | 88724347 | G | T | 0.34 | $2.40 \times 10^{-14}$ | 1.08 (1.06-1.10) | rs2002833 | 0.01 | 0.12 | MVD | 7.13 | 5 | Synonymous | - | - | - | - |
| rs2290902 | 16 | 88782676 | A | G | 0.11 | $2.70 \times 10^{-46}$ | 1.25 (1.21-1.29) | rs2002833 | 0.13 | 0.67 | PIEZO1 | 5.72 | NA | Synonymous | - | - | - | - |
| rs2859348 | 6 | 28359170 | A | G | 0.44 | $1.70 \times 10^{-11}$ | 1.07 (1.05-1.09) | rs7773004 | 0.07 | 0.29 | ZSCAN12 | 6.68 | 7 | Synonymous | - | - | - | - |
| rs34908386 | 16 | 88800913 | C | T | 0.14 | $8.30 \times 10^{-32}$ | 1.17 (1.14-1.21) | rs2002833 | 0.05 | 0.76 | PIEZO1:RP5-1142A6.2 | 20.90 | NA | Synonymous | - | - | - | - |
| rs34961555 | 6 | 26199903 | C | T | 0.10 | $3.30 \times 10^{-8}$ | 1.09 (1.06-1.13) | rs7773004 | 0.09 | 0.90 | HIST1H2BF | 4.34 | 4 | Synonymous | - | - | - | - |
| rs35353288 | 16 | 88800814 | G | A | 0.14 | $1.80 \times 10^{-31}$ | 1.17 (1.14-1.20) | rs2002833 | 0.05 | 0.76 | PIEZO1:RP5-1142A6.2 | 15.24 | NA | Synonymous | - | - | - | - |
| rs370520 | 6 | 28542520 | C | T | 0.46 | $5.00 \times 10^{-10}$ | 1.06 (1.04-1.08) | rs7773004 | 0.05 | 0.25 | SCAND3 | 9.28 | 6 | Synonymous | - | - | - | - |
| rs3746420 | 20 | 50140627 | G | C | 0.06 | $7.60 \times 10^{-10}$ | 1.13 (1.09-1.18) | rs3787184 | 0.16 | 0.80 | NFATC2 | 8.97 | 5 | Synonymous | - | - | - | - |

| | | | | | | | | | | | | | | | | | | |
|---|---|---|---|---|---|---|---|---|---|---|---|---|---|---|---|---|---|---|
| rs3749970 | 6 | 29342825 | G | A | 0.47 | 6.40x $10^{-9}$ | 0.95 (0.93-0.96) | rs7773004 | 0.08 | 0.31 | OR5V1:OR12D3 | 0.26 | NA | Synonymous | - | - | - | - |
| rs3752417 | 6 | 26045905 | G | C | 0.12 | 2.00x $10^{-10}$ | 1.10 (1.07-1.13) | rs7773004 | 0.09 | 0.82 | HIST1H3C | 14.99 | 4 | Synonymous | - | - | - | - |
| rs3752797 | 19 | 16339715 | C | T | 0.28 | 3.10x $10^{-8}$ | 0.94 (0.92-0.96) | rs12609241 | 0.82 | 1.00 | AP1M1 | 10.56 | NA | Synonymous | - | - | - | - |
| rs404240 | 6 | 29523957 | A | G | 0.12 | 8.40x $10^{-11}$ | 1.10 (1.07-1.13) | rs7773004 | 0.06 | 0.76 | UBD:GABBR1 | 14.02 | NA | Synonymous | - | - | - | - |
| rs4421818 | 12 | 50749294 | A | G | 0.35 | 2.20x $10^{-9}$ | 0.94 (0.92-0.96) | rs7308356 | 0.27 | 0.92 | FAM186A | 0.35 | 6 | Synonymous | - | - | - | - |
| rs450630 | 6 | 28542424 | G | A | 0.46 | 4.60x $10^{-10}$ | 1.06 (1.04-1.08) | rs7773004 | 0.05 | 0.25 | SCAND3 | 8.19 | 7 | Synonymous | - | - | - | - |
| rs4692549 | 4 | 26585881 | C | T | 0.42 | 5.40x $10^{-13}$ | 0.93 (0.91-0.95) | rs28558138 | 0.09 | 0.49 | TBC1D19 | 16.43 | 2b | Synonymous | - | - | - | - |
| rs4782430 | 16 | 88792047 | A | G | 0.12 | 4.50x $10^{-47}$ | 1.24 (1.20-1.28) | rs2002833 | 0.07 | 0.46 | PIEZO1 | 2.84 | NA | Synonymous | - | - | - | - |
| rs61742623 | 16 | 88802748 | C | T | 0.14 | 1.40x $10^{-30}$ | 1.17 (1.14-1.20) | rs2002833 | 0.05 | 0.76 | PIEZO1:RP5-1142A6.2 | 12.20 | 4 | Synonymous | - | - | - | - |
| rs6500491 | 16 | 88783521 | T | C | 0.10 | 1.50x $10^{-45}$ | 1.25 (1.21-1.29) | rs2002833 | 0.13 | 0.67 | PIEZO1 | 0.59 | 2b | Synonymous | - | - | - | - |
| rs687 | 2 | 68415767 | G | A | 0.38 | 7.30x $10^{-16}$ | 0.92 (0.91-0.94) | rs2861819 | 0.38 | 0.96 | RP11-474G23.1:PPP3R1 | 11.89 | NA | Synonymous | - | - | - | - |
| rs706793 | 12 | 50467769 | G | A | 0.41 | 1.40x $10^{-12}$ | 1.07 (1.05-1.09) | rs7308356 | 0.71 | 0.90 | ASIC1 | 11.00 | NA | Synonymous | - | - | - | - |

| | | | | | | | | | | | | | | | | | | |
|---|---|---|---|---|---|---|---|---|---|---|---|---|---|---|---|---|---|---|
| rs72683923 | 14 | 50735947 | T | C | 0.02 | 1.40x $10^{-8}$ | 0.82 (0.77-0.88) | rs72683923 | 1.00 | 1.00 | L2HGDH | 10.75 | 5 | Synonymous | - | - | - | - |
| rs7312252 | 12 | 50744171 | T | C | 0.35 | 1.50x $10^{-9}$ | 1.06 (1.04-1.08) | rs7308356 | 0.27 | 0.92 | FAM186A | 0.82 | 6 | Synonymous | - | - | - | - |
| rs735815 | 2 | 68511584 | A | G | 0.45 | 8.80x $10^{-12}$ | 0.94 (0.92-0.95) | rs2861819 | 0.26 | 0.74 | CNRIP1 | 1.38 | NA | Synonymous | - | - | - | - |
| rs759493 | 17 | 70036558 | C | T | 0.40 | 1.30x $10^{-21}$ | 1.10 (1.08-1.12) | rs9895127 | 0.87 | 1.00 | RP11-84E24.2:SOX9-AS1:AC007461.1 | 4.27 | NA | Synonymous | - | - | - | - |
| rs78905828 | 16 | 88802573 | G | A | 0.14 | 1.10x $10^{-31}$ | 1.17 (1.14-1.21) | rs2002833 | 0.05 | 0.76 | PIEZO1:RP5-1142A6.2 | 13.44 | 4 | Synonymous | - | - | - | - |
| rs8043637 | 16 | 88781125 | T | C | 0.11 | 1.70x $10^{-46}$ | 1.25 (1.21-1.29) | rs2002833 | 0.11 | 0.61 | CTU2 | 1.39 | 4 | Synonymous | - | - | - | - |
| rs8043924 | 16 | 88787704 | G | C | 0.12 | 1.10x $10^{-48}$ | 1.24 (1.21-1.28) | rs2002833 | 0.07 | 0.44 | PIEZO1 | 2.95 | 5 | Synonymous | - | - | - | - |
| rs8057031 | 16 | 88783449 | C | G | 0.44 | 1.70x $10^{-39}$ | 1.14 (1.11-1.16) | rs2002833 | 0.04 | 0.25 | PIEZO1 | 13.11 | 2b | Synonymous | - | - | - | - |
| rs836180 | 12 | 50503269 | C | T | 0.37 | 9.70x $10^{-16}$ | 1.08 (1.06-1.10) | rs7308356 | 0.95 | 1.00 | GPD1 | 15.32 | 5 | Synonymous | - | - | - | - |
| rs9379860 | 6 | 26370605 | T | C | 0.41 | 2.90x $10^{-10}$ | 1.06 (1.04-1.08) | rs7773004 | 0.11 | 0.38 | BTN3A2 | 0.65 | 7 | Synonymous | - | - | - | - |

[a]Based on NCBI Genome Build 37 (hg19). [b]The effect allele. [c]The alternate (non-effect) allele. [d]The effect allele frequency in the study population. [e]Odds ratio (95% confidence intervals), where OR > 1 indicative of increased risk with effect allele and OR < 1 a decreased risk with effect allele. [f]The top index signal at each replicated locus (as highlighted in Table 1). [g]The linkage disequilibrium score (R-square and D-prime) as calculated using the LDpair tool in LD Link (See URLs), in a study population of GBR (British in England and Scotland).[h]The nearest gene at which the SNP maps as calculated by positional mapping in FUMA. iThe functional consequences of SNPs was obtained by performing ANNOVAR gene-based annotation using Ensembl genes (build 85) in FUMA. [j]Human Genome Variation Society (HGVS) nomenclature on the description of

sequence variants (See URLs). [k]The GERP conservation score as calculated by Ensembl.[l]The SIFT and Polyphen-2 score of each variant as calculated by the gnomAD browser, red denotes damaging or deleterious missense variants. #denotes the four missense variants that are predicted to affect protein structure or function and are in linkage ($R2 \geq 0.22$ and $D' \geq 0.75$) with the top index SNP at each locus.

**Supplementary Table 4. Predicted functional intronic and intergenic variants at the replicated loci.** 163 genome-wide significant intronic and intergenic variants predicted to be deleterious according to a CADD ≥ 12.37, as identified by FUMA SNP2GENE.

| rsID | Chr | Position[a] | EA[b] | NEA[c] | EAF[d] | P-value | OR (95% CI)[e] | Functionality[f] | CADD[g] | RDB[h] |
|---|---|---|---|---|---|---|---|---|---|---|
| 1:10831584_GA_G | 1 | 10831584 | GA | G | 0.28 | 2.90E-123 | 1.29 (1.26-1.32) | intronic | 15.95 | NA |
| 1:10831714_TTCC_T | 1 | 10831714 | TTC C | T | 0.21 | 4.80E-91 | 1.27 (1.24-1.30) | intronic | 14.88 | NA |
| rs11121624 | 1 | 10850371 | G | A | 0.13 | 2.70E-29 | 0.85 (0.83-0.87) | intronic | 16.73 | 5 |
| rs11577258 | 1 | 10855315 | C | T | 0.05 | 5.70E-18 | 0.83 (0.80-0.87) | intronic | 13.44 | 4 |
| rs11577305 | 1 | 10855638 | C | G | 0.06 | 1.50E-17 | 0.84 (0.80-0.87) | intronic | 16.3 | 4 |
| rs12728592 | 1 | 10854073 | C | A | 0.02 | 2.50E-13 | 0.79 (0.74-0.84) | intronic | 21.2 | 2a |
| rs205475 | 1 | 10833105 | G | A | 0.07 | 9.60E-21 | 0.84 (0.80-0.87) | intronic | 13.24 | 5 |
| rs41307747 | 1 | 10836393 | C | A | 0.06 | 1.30E-23 | 0.82 (0.79-0.85) | intronic | 18.27 | 4 |
| rs58064215 | 1 | 10833232 | C | G | 0.29 | 4.50E-118 | 1.28 (1.25-1.30) | intronic | 18.68 | 5 |
| rs60694428 | 1 | 10822546 | G | A | 0.03 | 2.40E-19 | 0.78 (0.73-0.82) | intronic | 14.78 | 5 |
| rs61227177 | 1 | 10842769 | G | A | 0.04 | 8.90E-10 | 0.86 (0.82-0.90) | intronic | 14.88 | 5 |

| | | | | | | | | | | |
|---|---|---|---|---|---|---|---|---|---|---|
| rs761124 | 1 | 10856053 | G | T | 0.20 | 4.20E-24 | 1.13 (1.10-1.16) | intronic | 14.52 | 5 |
| rs776907 | 1 | 10845538 | T | A | 0.18 | 6.80E-33 | 1.16 (1.13-1.19) | intronic | 16.98 | 4 |
| rs776912 | 1 | 10847784 | T | A | 0.16 | 2.90E-31 | 1.17 (1.14-1.20) | intronic | 15.9 | 5 |
| rs11392514 | 1 | 115606679 | G | GA | 0.33 | 9.20E-10 | 0.94 (0.92-0.96) | intronic | 14.12 | NA |
| rs2798660 | 1 | 115359607 | A | G | 0.21 | 1.00E-08 | 0.94 (0.91-0.96) | intergenic | 13.82 | 6 |
| rs4839406 | 1 | 115604175 | A | C | 0.29 | 3.30E-10 | 0.94 (0.92-0.96) | intronic | 12.38 | 3a |
| rs340835 | 1 | 214163675 | G | A | 0.49 | 1.20E-10 | 1.06 (1.04-1.08) | intronic | 14.67 | NA |
| rs2785986 | 1 | 219706327 | A | G | 0.35 | 8.00E-09 | 0.94 (0.92-0.96) | intergenic | 14.7 | NA |
| rs2362541 | 2 | 30478453 | T | G | 0.49 | 2.30E-13 | 1.07 (1.05-1.09) | intronic | 19.95 | 5 |
| rs3813658 | 2 | 30473522 | T | C | 0.48 | 1.50E-11 | 1.07 (1.05-1.09) | intronic | 14.88 | NA |
| rs4952115 | 2 | 30478386 | G | T | 0.21 | 3.80E-14 | 1.09 (1.07-1.12) | intronic | 21.5 | 5 |
| rs3791679 | 2 | 56096892 | A | G | 0.22 | 1.90E-08 | 1.07 (1.04-1.09) | intronic | 17.76 | NA |
| 2:68423451_AG_A | 2 | 68423451 | AG | A | 0.26 | 2.60E-15 | 0.92 (0.90-0.94) | intronic | 16.87 | NA |

| | | | | | | | | | | |
|---|---|---|---|---|---|---|---|---|---|---|
| rs4347819 | 2 | 68479671 | A | G | 0.35 | 1.30E-54 | 0.85 (0.84-0.87) | intronic | 16.62 | 2b |
| rs4671894 | 2 | 68517936 | A | G | 0.31 | 7.20E-31 | 0.89 (0.87-0.90) | intronic | 13.95 | 5 |
| rs5831921 | 2 | 68478782 | G | GT | 0.38 | 7.40E-16 | 0.92 (0.91-0.94) | intronic | 14.48 | NA |
| rs7577500 | 2 | 68431296 | C | T | 0.26 | 4.50E-15 | 0.92 (0.90-0.94) | intronic | 17.82 | 7 |
| rs7593613 | 2 | 68483396 | A | T | 0.38 | 1.80E-16 | 0.92 (0.90-0.94) | intronic | 15.2 | 2b |
| rs7600326 | 2 | 68518280 | C | T | 0.39 | 5.30E-22 | 0.91 (0.89-0.93) | intronic | 16.26 | 5 |
| rs62164905 | 2 | 118879828 | A | G | 0.04 | 5.20E-09 | 1.15 (1.10-1.21) | intergenic | 12.78 | 4 |
| rs55889669 | 2 | 173194747 | G | A | 0.19 | 3.60E-10 | 0.93 (0.90-0.95) | intergenic | 20.7 | 4 |
| rs6775011 | 3 | 128396483 | T | C | 0.28 | 3.40E-09 | 1.07 (1.04-1.09) | intronic | 15.83 | 7 |
| rs13076750 | 3 | 188059443 | A | G | 0.25 | 8.00E-11 | 0.93 (0.91-0.95) | intronic | 17.26 | 5 |
| rs4132172 | 3 | 188061714 | C | T | 0.25 | 1.70E-10 | 0.93 (0.91-0.95) | intronic | 16.49 | NA |
| rs6787621 | 3 | 188066953 | T | G | 0.25 | 9.90E-11 | 1.07 (1.05-1.10) | intronic | 19.14 | 3a |
| rs59761494 | 4 | 26813959 | C | T | 0.35 | 2.40E-17 | 0.92 (0.90-0.94) | intergenic | 15.61 | 5 |

| | | | | | | | | | | |
|---|---|---|---|---|---|---|---|---|---|---|
| rs7672267 | 4 | 26728855 | A | G | 0.47 | 4.10E-11 | 0.94 (0.92-0.96) | intronic | 13.41 | 3a |
| rs57265257 | 4 | 57839280 | A | T | 0.19 | 2.00E-12 | 0.92 (0.89-0.94) | intronic | 16.2 | 3a |
| rs59708338 | 4 | 57725321 | G | A | 0.15 | 1.20E-09 | 0.92 (0.90-0.95) | intergenic | 13.54 | 7 |
| rs6852182 | 4 | 57775116 | G | C | 0.19 | 2.40E-12 | 0.92 (0.89-0.94) | intronic | 20 | 3a |
| rs6853156 | 4 | 57774843 | C | T | 0.19 | 1.50E-12 | 0.92 (0.89-0.94) | intronic | 18.46 | 4 |
| rs251216 | 5 | 127543280 | A | G | 0.25 | 6.20E-26 | 1.12 (1.10-1.15) | intergenic | 15.06 | NA |
| rs36702 | 5 | 127552723 | T | A | 0.26 | 1.50E-22 | 1.11 (1.09-1.14) | intergenic | 18.1 | 1f |
| rs36712 | 5 | 127561133 | G | C | 0.26 | 2.90E-23 | 1.12 (1.09-1.14) | intergenic | 12.38 | 5 |
| rs72794380 | 5 | 127466406 | A | G | 0.03 | 8.50E-09 | 0.85 (0.81-0.90) | intronic | 15.13 | 7 |
| rs10447201 | 5 | 158418735 | A | C | 0.30 | 1.70E-08 | 1.06 (1.04-1.08) | intronic | 13.22 | 5 |
| rs11135046 | 5 | 158230013 | G | T | 0.46 | 1.60E-32 | 1.12 (1.10-1.14) | intronic | 15.46 | 5 |
| rs11745795 | 5 | 158202920 | A | G | 0.18 | 3.80E-10 | 0.92 (0.90-0.95) | intronic | 21.5 | 3a |
| rs1422797 | 5 | 158320876 | T | A | 0.37 | 4.30E-17 | 1.09 (1.07-1.11) | intronic | 14.62 | 7 |

| | | | | | | | | | | |
|---|---|---|---|---|---|---|---|---|---|---|
| rs1422798 | 5 | 158320877 | C | G | 0.37 | 4.30E-17 | 1.09 (1.07-1.11) | intronic | 15.64 | 7 |
| rs1544754 | 5 | 158414305 | T | C | 0.19 | 3.40E-09 | 1.08 (1.05-1.10) | intronic | 13.57 | 5 |
| rs17543752 | 5 | 158332380 | T | C | 0.37 | 9.50E-17 | 1.09 (1.06-1.11) | intronic | 16.47 | 5 |
| rs17715000 | 5 | 158259583 | T | C | 0.40 | 3.20E-27 | 1.11 (1.09-1.13) | intronic | 12.68 | 6 |
| rs1864940 | 5 | 158196503 | T | A | 0.18 | 4.70E-10 | 0.93 (0.90-0.95) | intronic | 15.42 | 6 |
| rs1978706 | 5 | 158274071 | T | C | 0.35 | 7.20E-09 | 1.06 (1.04-1.08) | intronic | 20.7 | 6 |
| rs2072495 | 5 | 158296996 | C | T | 0.38 | 1.30E-16 | 1.08 (1.06-1.11) | intronic | 18.59 | NA |
| rs4704959 | 5 | 158202291 | A | C | 0.18 | 4.00E-10 | 0.92 (0.90-0.95) | intronic | 12.96 | 2b |
| rs66666124 | 5 | 158392702 | G | A | 0.18 | 2.50E-08 | 0.93 (0.91-0.96) | intronic | 16.77 | 5 |
| rs6867332 | 5 | 158380961 | A | T | 0.39 | 9.30E-09 | 0.94 (0.93-0.96) | intronic | 15.15 | 4 |
| rs6887211 | 5 | 158446223 | C | T | 0.29 | 2.70E-08 | 1.06 (1.04-1.08) | intronic | 17.48 | 4 |
| rs72643433 | 5 | 158364449 | G | A | 0.24 | 4.80E-09 | 1.07 (1.04-1.09) | intronic | 16.92 | 5 |
| rs7707918 | 5 | 158270225 | A | C | 0.38 | 2.50E-16 | 1.08 (1.06-1.11) | intronic | 14.32 | 5 |

| | | | | | | | | | | |
|---|---|---|---|---|---|---|---|---|---|---|
| rs7732733 | 5 | 158167192 | A | G | 0.21 | 3.60E-10 | 0.93 (0.91-0.95) | intronic | 18.4 | 2b |
| rs7736883 | 5 | 158343969 | A | G | 0.37 | 1.00E-16 | 1.09 (1.06-1.11) | intronic | 15.34 | 7 |
| rs7737500 | 5 | 158317804 | G | C | 0.37 | 1.70E-16 | 1.08 (1.06-1.11) | intronic | 13.7 | 5 |
| rs7737883 | 5 | 158340872 | G | C | 0.22 | 2.20E-11 | 1.08 (1.06-1.11) | intronic | 16.91 | 5 |
| rs891903 | 5 | 158279638 | G | A | 0.24 | 1.20E-08 | 1.07 (1.04-1.09) | intronic | 15.76 | NA |
| 6:27914359_AAATGAACTGAAGGAGAGGTCCC_A | 6 | 27914359 | AAATGAACTGAAGGAGAGGTCCC | A | 0.11 | 6.70E-10 | 1.10 (1.07-1.13) | intergenic | 13.54 | NA |
| 6:28277346_AT_A | 6 | 28277346 | AT | A | 0.11 | 9.00E-10 | 1.10 (1.06-1.13) | intergenic | 12.91 | NA |
| 6:28315542_CCAAT_C | 6 | 28315542 | CCAAT | C | 0.24 | 1.80E-08 | 1.07 (1.04-1.09) | intronic | 12.96 | NA |
| rs113517046 | 6 | 28770596 | A | AC | 0.49 | 1.10E-09 | 1.06 (1.04-1.08) | intergenic | 14.58 | NA |
| rs11967609 | 6 | 28656109 | T | A | 0.29 | 4.60E-08 | 1.06 (1.04-1.08) | intergenic | 13.29 | 6 |
| rs12215773 | 6 | 27039233 | T | C | 0.22 | 6.70E-10 | 1.07 (1.05-1.10) | intergenic | 13.57 | 6 |

| | | | | | | | | | | |
|---|---|---|---|---|---|---|---|---|---|---|
| rs1233572 | 6 | 28703244 | T | C | 0.49 | 4.40E-09 | 1.06 (1.04-1.08) | intergenic | 16.07 | 4 |
| rs1233583 | 6 | 28715566 | G | A | 0.11 | 1.30E-09 | 1.09 (1.06-1.13) | intergenic | 14.21 | 2b |
| rs13193480 | 6 | 27702561 | A | G | 0.11 | 1.30E-08 | 1.09 (1.06-1.12) | intergenic | 15.29 | 5 |
| rs13198716 | 6 | 26582035 | C | T | 0.10 | 5.10E-09 | 1.09 (1.06-1.13) | intergenic | 12.54 | 1f |
| rs13203358 | 6 | 26590578 | A | T | 0.13 | 4.60E-09 | 1.09 (1.06-1.12) | intergenic | 13.55 | 5 |
| rs13211434 | 6 | 26743531 | G | C | 0.11 | 2.60E-09 | 1.09 (1.06-1.13) | intergenic | 13.07 | 6 |
| rs13211507 | 6 | 28257377 | T | C | 0.11 | 8.20E-10 | 1.10 (1.06-1.13) | intronic | 13.79 | 6 |
| rs149947 | 6 | 27972433 | A | T | 0.24 | 6.40E-10 | 1.07 (1.05-1.09) | intergenic | 13.59 | 5 |
| rs149974 | 6 | 27985096 | T | C | 0.39 | 4.10E-10 | 1.06 (1.04-1.08) | intergenic | 13.71 | 6 |
| rs188105 | 6 | 28071393 | C | T | 0.29 | 9.70E-10 | 1.07 (1.04-1.09) | intergenic | 15.23 | 6 |
| rs209181 | 6 | 28792477 | G | A | 0.17 | 5.50E-09 | 1.08 (1.05-1.10) | intergenic | 12.38 | NA |
| rs2207699 | 6 | 29331819 | A | T | 0.47 | 7.00E-10 | 0.94 (0.92-0.96) | intronic | 12.69 | 7 |
| rs28749522 | 6 | 29338563 | G | C | 0.47 | 2.30E-09 | 0.94 (0.93-0.96) | intronic | 13.66 | NA |

| | | | | | | | | | | |
|---|---|---|---|---|---|---|---|---|---|---|
| rs28751297 | 6 | 29338562 | T | TAAA | 0.47 | 2.30E-09 | 0.94 (0.93-0.96) | intronic | 13.23 | NA |
| rs3130730 | 6 | 29126943 | T | C | 0.49 | 1.50E-08 | 0.95 (0.93-0.97) | intergenic | 14.99 | 6 |
| rs3130746 | 6 | 29153155 | A | G | 0.11 | 8.40E-10 | 1.09 (1.06-1.13) | intergenic | 12.86 | 7 |
| rs3130810 | 6 | 29190181 | A | T | 0.49 | 1.30E-08 | 0.95 (0.93-0.96) | intergenic | 12.72 | 6 |
| rs3132392 | 6 | 28838629 | T | C | 0.11 | 1.50E-09 | 1.09 (1.06-1.13) | intergenic | 13.32 | 5 |
| rs3135300 | 6 | 28824397 | T | A | 0.11 | 1.40E-09 | 1.09 (1.06-1.13) | intergenic | 13.51 | 6 |
| rs34104395 | 6 | 26478252 | C | T | 0.12 | 2.70E-08 | 1.09 (1.05-1.12) | intergenic | 13.34 | 4 |
| rs34573979 | 6 | 27480526 | C | T | 0.11 | 2.90E-08 | 1.09 (1.06-1.12) | intergenic | 15.35 | 5 |
| rs35193936 | 6 | 28076270 | A | G | 0.24 | 1.40E-09 | 1.07 (1.05-1.09) | intergenic | 12.49 | 6 |
| rs35657082 | 6 | 27067657 | A | T | 0.09 | 2.80E-08 | 0.91 (0.88-0.94) | intergenic | 13.36 | 7 |
| rs35744819 | 6 | 28318331 | G | T | 0.11 | 3.10E-09 | 1.09 (1.06-1.13) | intronic | 13.57 | 3a |
| rs56075693 | 6 | 28290328 | T | G | 0.11 | 8.70E-10 | 1.10 (1.06-1.13) | intergenic | 12.71 | 5 |
| rs56405707 | 6 | 27640246 | G | A | 0.11 | 9.40E-09 | 1.09 (1.06-1.12) | intergenic | 13.34 | 4 |

| | | | | | | | | | | |
|---|---|---|---|---|---|---|---|---|---|---|
| rs67878650 | 6 | 28142587 | G | C | 0.24 | 3.30E-10 | 1.07 (1.05-1.10) | intergenic | 12.53 | 7 |
| rs71559054 | 6 | 27896799 | A | C | 0.11 | 5.60E-10 | 1.10 (1.07-1.13) | intergenic | 14.09 | 7 |
| rs7765989 | 6 | 28400295 | T | C | 0.23 | 4.00E-09 | 1.07 (1.05-1.09) | intronic | 12.95 | 6 |
| rs9257189 | 6 | 28757555 | C | G | 0.11 | 9.80E-10 | 1.09 (1.06-1.13) | intergenic | 14.92 | 4 |
| rs9380052 | 6 | 28064623 | T | C | 0.24 | 1.40E-09 | 1.07 (1.05-1.09) | intergenic | 12.61 | 6 |
| rs9468195 | 6 | 27676309 | A | G | 0.28 | 3.80E-09 | 1.07 (1.04-1.09) | intergenic | 15.14 | 6 |
| rs11967262 | 6 | 43760327 | C | G | 0.48 | 5.20E-09 | 1.06 (1.04-1.08) | intergenic | 14.05 | 7 |
| rs4711750 | 6 | 43757082 | T | A | 0.49 | 1.20E-08 | 1.06 (1.04-1.08) | intergenic | 12.87 | 5 |
| rs998584 | 6 | 43757896 | C | A | 0.48 | 6.30E-09 | 1.06 (1.04-1.08) | intergenic | 12.4 | NA |
| rs1936800 | 6 | 127436064 | C | T | 0.48 | 2.60E-09 | 1.06 (1.04-1.08) | intergenic | 17.77 | 7 |
| rs1936807 | 6 | 127448249 | C | G | 0.49 | 2.60E-08 | 1.06 (1.04-1.08) | intronic | 16.5 | 6 |
| rs1441249 | 8 | 87614243 | T | C | 0.48 | 2.50E-10 | 0.94 (0.92-0.96) | intronic | 20.2 | NA |
| rs4740661 | 9 | 227621 | C | T | 0.14 | 4.10E-11 | 0.91 (0.89-0.94) | intronic | 14.94 | 4 |

| | | | | | | | | | | |
|---|---|---|---|---|---|---|---|---|---|---|
| rs10117339 | 9 | 118327941 | C | T | 0.48 | 2.90E-09 | 0.94 (0.93-0.96) | intergenic | 12.61 | 6 |
| rs10817747 | 9 | 118095937 | T | A | 0.49 | 2.10E-11 | 0.94 (0.92-0.96) | intronic | 15.33 | 5 |
| rs10982705 | 9 | 118119365 | C | T | 0.46 | 2.90E-11 | 0.94 (0.92-0.96) | intronic | 12.68 | 3a |
| rs10982846 | 9 | 118378191 | G | T | 0.25 | 1.10E-13 | 0.92 (0.90-0.94) | intergenic | 20.8 | 4 |
| rs12686630 | 9 | 118199807 | A | C | 0.37 | 1.30E-14 | 0.93 (0.91-0.94) | intergenic | 13.97 | 4 |
| rs2027932 | 9 | 118374481 | T | C | 0.48 | 3.10E-09 | 0.94 (0.93-0.96) | intergenic | 15.39 | 5 |
| rs2188065 | 9 | 118370136 | T | C | 0.39 | 3.60E-09 | 0.94 (0.93-0.96) | intergenic | 14.55 | 6 |
| rs2188066 | 9 | 118370953 | T | C | 0.17 | 1.10E-09 | 0.92 (0.90-0.95) | intergenic | 13.26 | 6 |
| rs2188067 | 9 | 118371019 | A | C | 0.26 | 1.50E-13 | 0.92 (0.90-0.94) | intergenic | 12.37 | 7 |
| rs4997383 | 9 | 118378025 | T | C | 0.25 | 1.20E-13 | 0.92 (0.90-0.94) | intergenic | 18.87 | 3a |
| rs7024456 | 9 | 118373854 | T | G | 0.25 | 1.60E-13 | 0.92 (0.90-0.94) | intergenic | 12.58 | 5 |
| rs72748112 | 9 | 118118677 | C | T | 0.46 | 2.80E-11 | 0.94 (0.92-0.96) | intronic | 13.41 | 7 |
| rs76423767 | 9 | 118358637 | A | G | 0.10 | 1.70E-08 | 1.09 (1.06-1.13) | intergenic | 21.4 | 2b |

| | | | | | | | | | | |
|---|---|---|---|---|---|---|---|---|---|---|
| rs79333681 | 9 | 118113350 | T | A | 0.29 | 2.30E-09 | 0.94 (0.92-0.96) | intronic | 15.34 | 5 |
| rs10876055 | 12 | 50940983 | T | G | 0.36 | 2.10E-11 | 0.94 (0.92-0.95) | intronic | 16.26 | 1f |
| rs11169339 | 12 | 50651456 | T | A | 0.35 | 2.20E-09 | 0.94 (0.92-0.96) | intronic | 13.04 | 4 |
| rs11169348 | 12 | 50665946 | G | T | 0.35 | 1.50E-09 | 0.94 (0.92-0.96) | intronic | 14.57 | 1f |
| rs11169376 | 12 | 50713091 | G | C | 0.35 | 2.20E-09 | 0.94 (0.92-0.96) | intergenic | 12.6 | 6 |
| rs11169377 | 12 | 50713094 | G | A | 0.35 | 2.00E-09 | 0.94 (0.92-0.96) | intergenic | 12.89 | 6 |
| rs141978158 | 12 | 50605755 | A | AC | 0.35 | 1.00E-09 | 0.94 (0.92-0.96) | intronic | 17.22 | NA |
| rs2111988 | 12 | 50668538 | C | T | 0.35 | 2.20E-09 | 0.94 (0.92-0.96) | intronic | 13.46 | 1f |
| rs2251603 | 12 | 51049653 | G | A | 0.35 | 4.90E-14 | 1.08 (1.06-1.10) | intronic | 12.56 | 1f |
| rs4986838 | 12 | 51203376 | T | C | 0.41 | 1.70E-13 | 0.93 (0.91-0.95) | intronic | 15.18 | 1f |
| rs7132551 | 12 | 50646754 | G | A | 0.35 | 1.60E-09 | 0.94 (0.92-0.96) | intronic | 12.54 | 1f |
| rs7136648 | 12 | 50624822 | G | A | 0.46 | 1.00E-10 | 0.94 (0.92-0.96) | intronic | 14.58 | 6 |
| rs751859685 | 12 | 51008201 | AT | A | 0.36 | 5.90E-11 | 0.94 (0.92-0.96) | intronic | 15.63 | NA |

| | | | | | | | | | | |
|---|---|---|---|---|---|---|---|---|---|---|
| rs7960015 | 12 | 51145194 | G | A | 0.32 | 2.00E-08 | 1.06 (1.04-1.08) | intergenic | 15.45 | 6 |
| rs860698 | 12 | 50486322 | T | C | 0.41 | 4.00E-12 | 1.07 (1.05-1.09) | intronic | 14.3 | 6 |
| 13:73632830_AC_A | 13 | 73632830 | AC | A | 0.25 | 2.70E-10 | 0.93 (0.91-0.95) | intronic | 22.5 | NA |
| 15:96167440_TG_T | 15 | 96167440 | TG | T | 0.16 | 1.20E-17 | 1.12 (1.09-1.15) | intergenic | 15.81 | NA |
| 15:96190141_CA_C | 15 | 96190141 | CA | C | 0.07 | 2.80E-08 | 1.11 (1.07-1.16) | intergenic | 14.89 | NA |
| rs10520792 | 15 | 96151914 | C | T | 0.28 | 1.80E-08 | 1.06 (1.04-1.08) | intergenic | 17.44 | 5 |
| rs11852492 | 15 | 96167544 | T | C | 0.16 | 8.90E-18 | 1.12 (1.09-1.15) | intergenic | 13 | 5 |
| rs11857783 | 15 | 96170238 | C | G | 0.16 | 1.30E-17 | 1.12 (1.09-1.15) | intergenic | 17.55 | 5 |
| rs11857837 | 15 | 96170245 | G | A | 0.16 | 1.30E-17 | 1.12 (1.09-1.15) | intergenic | 16.11 | 5 |
| rs12593084 | 15 | 96161743 | G | A | 0.16 | 1.90E-17 | 1.12 (1.09-1.15) | intergenic | 13.57 | 7 |
| rs12593086 | 15 | 96154502 | G | C | 0.06 | 6.40E-10 | 1.13 (1.09-1.18) | intergenic | 13.17 | 7 |
| rs34720717 | 15 | 96167935 | G | GA | 0.16 | 1.00E-17 | 1.12 (1.09-1.15) | intergenic | 14.58 | NA |
| rs55790747 | 15 | 96169774 | G | C | 0.16 | 2.80E-17 | 1.12 (1.09-1.15) | intergenic | 16.42 | 4 |

| | | | | | | | | | | |
|---|---|---|---|---|---|---|---|---|---|---|
| rs59208367 | 15 | 96190148 | A | T | 0.07 | 2.70E-08 | 1.11 (1.07-1.16) | intergenic | 14.76 | 6 |
| rs7162535 | 15 | 96158598 | T | C | 0.28 | 1.60E-08 | 1.06 (1.04-1.09) | intergenic | 12.54 | 3a |
| rs7163129 | 15 | 96158885 | T | C | 0.28 | 1.60E-08 | 1.06 (1.04-1.09) | intergenic | 16.85 | 3a |
| rs74918319 | 15 | 96170043 | T | A | 0.07 | 3.60E-12 | 1.14 (1.10-1.18) | intergenic | 15.27 | 5 |
| rs113478686 | 16 | 88850897 | C | CT | 0.23 | 6.50E-35 | 1.15 (1.13-1.18) | intronic | 13.26 | NA |
| rs78579285 | 16 | 88813060 | C | T | 0.18 | 6.20E-30 | 1.15 (1.12-1.18) | intronic | 15.41 | 5 |
| rs8043839 | 16 | 88847437 | T | C | 0.45 | 2.50E-08 | 0.95 (0.93-0.97) | intronic | 12.79 | 3a |
| rs9935872 | 16 | 88788477 | A | G | 0.12 | 2.40E-48 | 1.24 (1.21-1.28) | intronic | 13.28 | 5 |
| rs236591 | 17 | 68206588 | T | C | 0.49 | 1.80E-08 | 1.06 (1.04-1.08) | intergenic | 12.54 | 6 |
| rs185799410 | 20 | 57466093 | G | T | 0.02 | 1.80E-12 | 1.24 (1.17-1.32) | intronic | 13.54 | 2b |
| rs6062340 | 20 | 62683275 | T | C | 0.10 | 2.10E-10 | 1.10 (1.07-1.14) | intergenic | 14.44 | 2b |

[a]Based on NCBI Genome Build 37 (hg19). [b]The effect allele. [c]The alternate (non-effect) allele. [d]The effect allele frequency in the study population. [e]Odds ratio (95% confidence intervals), where OR > 1 indicative of increased risk with effect allele and OR < 1 a decreased risk with effect allele. [f]The functional consequences of SNPs on the respective gene obtained by performing ANNOVAR gene-based annotation using Ensembl genes (build 85) in FUMA. [g]The combined annotation depletion score as calculated in FUMA. [h]The RegulomeDB score as calculated in FUMA, where blue variants are those that have an RDB score 2a or 2b, and red variants are those with an RDB score 1f or less.

**Supplementary Table 5. Genes mapped to the varicose veins associated loci using the four mapping strategies.** 237 unique genes were mapped to 39 of 46 replicated loci by one or more gene mapping strategies (see Methods). 204 genes were mapped via positional mapping in FUMA, 80 genes were mapped via eQTL mapping in FUMA, 117 genes were mapped using MAGMA and 14 genes were mapped using summary-based mendelian randomisation. In total, 61 unique genes were mapped to novel loci. Overlap between the four different mapping strategies is shown.

| Novel genes | Chromosome | Lead SNP[a] | Position[b] | FUMA Positionally Mapped Genes | FUMA eQTL Mapped Genes | MAGMA Mapped Genes | SMR Mapped Genes | Number of Gene Mapping Approaches |
|---|---|---|---|---|---|---|---|---|
| | 1 | rs11121615 | 10825577 | CASZ1 | | CASZ1 | | 2 |
| Yes | 1 | rs7518191 | 115603413 | TSPAN2 | | TSPAN2 | | 2 |
| Yes | 1 | rs7518191 | 115603413 | SYCP1 | | | | 1 |
| Yes | 1 | rs7518191 | 115603413 | | BCAS2 | | | 1 |
| Yes | 1 | rs7518191 | 115603413 | | TSHB | | | 1 |
| Yes | 1 | rs340875 | 214158986 | PROX1 | | PROX1 | | 2 |
| | 2 | rs9967884 | 30488183 | LBH | LBH | LBH | LBH | 4 |
| | 2 | rs3791679 | 56096892 | EFEMP1 | | | | 1 |
| | 2 | rs2861819 | 68489221 | CNRIP1 | CNRIP1 | CNRIP1 | | 3 |
| | 2 | rs2861819 | 68489221 | PNO1 | PNO1 | PNO1 | | 3 |
| | 2 | rs2861819 | 68489221 | PPP3R1 | PPP3R1 | PPP3R1 | | 3 |
| | 2 | rs2861819 | 68489221 | RP11-474G23.1 | | RP11-474G23.1 | | 2 |
| | 2 | rs2861819 | 68489221 | WDR92 | WDR92 | WDR92 | WDR92 | 4 |
| | 2 | rs2861819 | 68489221 | C1D | | | | 1 |
| Yes | 2 | rs4849044 | 112898933 | FBLN7 | FBLN7 | FBLN7 | FBLN7 | 4 |
| Yes | 2 | rs4849044 | 112898933 | TMEM87B | TMEM87B | TMEM87B | | 3 |
| Yes | 2 | rs4849044 | 112898933 | | CHCHD5 | | | 1 |
| Yes | 2 | rs4849044 | 112898933 | | POLR1B | | | 1 |

| | | | | | | | | |
|---|---|---|---|---|---|---|---|---|
| Yes | 2 | rs4849044 | 112898933 | | ZC3H8 | | | 1 |
| Yes | 2 | rs17819430 | 118886398 | | INSIG2 | | | 1 |
| Yes | 3 | rs844176 | 14827080 | FGD5 | | FGD5 | | 2 |
| Yes | 3 | rs844176 | 14827080 | C3orf20 | | | | 1 |
| | 3 | rs2713575 | 128294355 | C3orf27 | | C3orf27 | | 2 |
| | 3 | rs2713575 | 128294355 | GATA2 | | GATA2 | | 2 |
| | 3 | rs2713575 | 128294355 | RPN1 | RPN1 | RPN1 | | 3 |
| | 3 | rs2713575 | 128294355 | | MCM2 | | | 1 |
| | 3 | rs2713575 | 128294355 | | PLXND1 | | | 1 |
| | 3 | rs2713575 | 128294355 | | | RAB7A | | 1 |
| Yes | 3 | rs9877579 | 188058716 | LPP | | | | 1 |
| | 4 | rs28558138 | 26818080 | TBC1D19 | | TBC1D19 | | 2 |
| Yes | 4 | rs56155140 | 57824451 | NOA1 | | NOA1 | | 2 |
| Yes | 4 | rs56155140 | 57824451 | POLR2B | POLR2B | | | 2 |
| Yes | 4 | rs56155140 | 57824451 | REST | REST | REST | | 3 |
| Yes | 4 | rs56155140 | 57824451 | IGFBP7 | | | | 1 |
| Yes | 4 | rs1471251 | 87976359 | AFF1 | AFF1 | AFF1 | | 3 |
| Yes | 4 | rs1471251 | 87976359 | | MAPK10 | | | 1 |
| Yes | 4 | rs34154818 | 89726823 | FAM13A | FAM13A | FAM13A | | 3 |
| Yes | 4 | rs10007409 | 120142306 | RP11-455G16.1 | | RP11-455G16.1 | | 2 |
| Yes | 4 | rs10007409 | 120142306 | USP53 | | | | 1 |
| Yes | 4 | rs10007409 | 120142306 | | | MYOZ2 | | 1 |
| Yes | 4 | rs11728719 | 186696172 | SORBS2 | | | | 1 |
| | 5 | rs3749748 | 127350549 | CTXN3 | | CTXN3 | | 2 |
| | 5 | rs3749748 | 127350549 | FBN2 | FBN2 | | | 2 |
| | 5 | rs3749748 | 127350549 | SLC12A2 | SLC12A2 | SLC12A2 | | 3 |
| | 5 | rs3749748 | 127350549 | | | | CTC-228N24.3 | 1 |

| | | | | | | | |
|---|---|---|---|---|---|---|---|
| 5 | rs11135046 | 158230013 | EBF1 | | EBF1 | | 2 |
| 6 | rs7773004 | 26267755 | ABT1 | ABT1 | ABT1 | | 3 |
| 6 | rs7773004 | 26267755 | BTN1A1 | | BTN1A1 | | 2 |
| 6 | rs7773004 | 26267755 | BTN2A1 | | BTN2A1 | | 2 |
| 6 | rs7773004 | 26267755 | BTN2A2 | BTN2A2 | BTN2A2 | | 3 |
| 6 | rs7773004 | 26267755 | BTN3A1 | BTN3A1 | | | 2 |
| 6 | rs7773004 | 26267755 | BTN3A2 | BTN3A2 | BTN3A2 | | 3 |
| 6 | rs7773004 | 26267755 | BTN3A3 | BTN3A3 | BTN3A3 | | 3 |
| 6 | rs7773004 | 26267755 | C6orf100 | | C6orf100 | | 2 |
| 6 | rs7773004 | 26267755 | GPX6 | | GPX6 | | 2 |
| 6 | rs7773004 | 26267755 | HFE | HFE | HFE | | 3 |
| 6 | rs7773004 | 26267755 | HIST1H1A | | HIST1H1A | | 2 |
| 6 | rs7773004 | 26267755 | HIST1H1B | | HIST1H1B | | 2 |
| 6 | rs7773004 | 26267755 | HIST1H2AC | | HIST1H2AC | | 2 |
| 6 | rs7773004 | 26267755 | HIST1H2AJ | | HIST1H2AJ | | 2 |
| 6 | rs7773004 | 26267755 | HIST1H2AL | | HIST1H2AL | | 2 |
| 6 | rs7773004 | 26267755 | HIST1H2BC | | HIST1H2BC | | 2 |
| 6 | rs7773004 | 26267755 | HIST1H2BD | | HIST1H2BD | | 2 |
| 6 | rs7773004 | 26267755 | HIST1H2BE | | HIST1H2BE | | 2 |
| 6 | rs7773004 | 26267755 | HIST1H2BF | | HIST1H2BF | | 2 |
| 6 | rs7773004 | 26267755 | HIST1H2BJ | HIST1H2BJ | | | 2 |
| 6 | rs7773004 | 26267755 | HIST1H2BK | HIST1H2BK | | | 2 |
| 6 | rs7773004 | 26267755 | HIST1H2BL | | HIST1H2BL | | 2 |
| 6 | rs7773004 | 26267755 | HIST1H2BN | | HIST1H2BN | | 2 |
| 6 | rs7773004 | 26267755 | HIST1H3C | | HIST1H3C | | 2 |
| 6 | rs7773004 | 26267755 | HIST1H3I | | HIST1H3I | | 2 |
| 6 | rs7773004 | 26267755 | HIST1H3J | | HIST1H3J | | 2 |

| | | | | | | | |
|---|---|---|---|---|---|---|---|
| 6 | rs7773004 | 26267755 | HIST1H4A | | HIST1H4A | | 2 |
| 6 | rs7773004 | 26267755 | HIST1H4C | | HIST1H4C | | 2 |
| 6 | rs7773004 | 26267755 | HIST1H4L | | HIST1H4L | | 2 |
| 6 | rs7773004 | 26267755 | HMGN4 | HMGN4 | HMGN4 | | 3 |
| 6 | rs7773004 | 26267755 | LRRC16A | | LRRC16A | | 2 |
| 6 | rs7773004 | 26267755 | OR12D3 | | OR12D3 | | 2 |
| 6 | rs7773004 | 26267755 | OR2B2 | | OR2B2 | | 2 |
| 6 | rs7773004 | 26267755 | OR2H2 | OR2H2 | | | 2 |
| 6 | rs7773004 | 26267755 | OR2J2 | | OR2J2 | | 2 |
| 6 | rs7773004 | 26267755 | OR5V1 | | OR5V1 | | 2 |
| 6 | rs7773004 | 26267755 | PGBD1 | | PGBD1 | | 2 |
| 6 | rs7773004 | 26267755 | POM121L2 | | POM121L2 | | 2 |
| 6 | rs7773004 | 26267755 | PRSS16 | PRSS16 | | | 2 |
| 6 | rs7773004 | 26267755 | SCAND3 | SCAND3 | SCAND3 | | 3 |
| 6 | rs7773004 | 26267755 | SCGN | | SCGN | | 2 |
| 6 | rs7773004 | 26267755 | SLC17A2 | | SLC17A2 | | 2 |
| 6 | rs7773004 | 26267755 | SLC17A3 | | SLC17A3 | | 2 |
| 6 | rs7773004 | 26267755 | SLC17A4 | | SLC17A4 | | 2 |
| 6 | rs7773004 | 26267755 | TRIM27 | TRIM27 | TRIM27 | | 3 |
| 6 | rs7773004 | 26267755 | TRIM38 | TRIM38 | TRIM38 | | 3 |
| 6 | rs7773004 | 26267755 | ZKSCAN3 | ZKSCAN3 | ZKSCAN3 | | 3 |
| 6 | rs7773004 | 26267755 | ZKSCAN4 | | ZKSCAN4 | | 2 |
| 6 | rs7773004 | 26267755 | ZKSCAN8 | ZKSCAN8 | ZKSCAN8 | | 3 |
| 6 | rs7773004 | 26267755 | ZNF165 | ZNF165 | ZNF165 | | 3 |
| 6 | rs7773004 | 26267755 | ZNF311 | ZNF311 | | | 2 |
| 6 | rs7773004 | 26267755 | ZNF322 | ZNF322 | ZNF322 | | 3 |
| 6 | rs7773004 | 26267755 | ZNF391 | ZNF391 | | | 2 |

| | | | | | | | |
|---|---|---|---|---|---|---|---|
| 6 | rs7773004 | 26267755 | ZSCAN12 | ZSCAN12 | ZSCAN12 | | 3 |
| 6 | rs7773004 | 26267755 | ZSCAN16 | | ZSCAN16 | | 2 |
| 6 | rs7773004 | 26267755 | ZSCAN23 | ZSCAN23 | | | 2 |
| 6 | rs7773004 | 26267755 | ZSCAN31 | | ZSCAN31 | | 2 |
| 6 | rs7773004 | 26267755 | ZSCAN9 | ZSCAN9 | ZSCAN9 | | 3 |
| 6 | rs7773004 | 26267755 | GABBR1 | | | | 1 |
| 6 | rs7773004 | 26267755 | GPX5 | | | | 1 |
| 6 | rs7773004 | 26267755 | HIST1H1C | | | | 1 |
| 6 | rs7773004 | 26267755 | HIST1H1D | | | | 1 |
| 6 | rs7773004 | 26267755 | HIST1H1E | | | | 1 |
| 6 | rs7773004 | 26267755 | HIST1H1T | | | | 1 |
| 6 | rs7773004 | 26267755 | HIST1H2AB | | | | 1 |
| 6 | rs7773004 | 26267755 | HIST1H2AD | | | | 1 |
| 6 | rs7773004 | 26267755 | HIST1H2AE | | | | 1 |
| 6 | rs7773004 | 26267755 | HIST1H2AG | | | | 1 |
| 6 | rs7773004 | 26267755 | HIST1H2AH | | | | 1 |
| 6 | rs7773004 | 26267755 | HIST1H2AI | | | | 1 |
| 6 | rs7773004 | 26267755 | HIST1H2AK | | | | 1 |
| 6 | rs7773004 | 26267755 | HIST1H2AM | | | | 1 |
| 6 | rs7773004 | 26267755 | HIST1H2BB | | | | 1 |
| 6 | rs7773004 | 26267755 | HIST1H2BG | | | | 1 |
| 6 | rs7773004 | 26267755 | HIST1H2BH | | | | 1 |
| 6 | rs7773004 | 26267755 | HIST1H2BI | | | | 1 |
| 6 | rs7773004 | 26267755 | HIST1H2BM | | | | 1 |
| 6 | rs7773004 | 26267755 | HIST1H2BO | | | | 1 |
| 6 | rs7773004 | 26267755 | HIST1H3A | | | | 1 |
| 6 | rs7773004 | 26267755 | HIST1H3B | | | | 1 |

| | | | | |
|---|---|---|---|---|
| 6 | rs7773004 | 26267755 | HIST1H3D | 1 |
| 6 | rs7773004 | 26267755 | HIST1H3E | 1 |
| 6 | rs7773004 | 26267755 | HIST1H3F | 1 |
| 6 | rs7773004 | 26267755 | HIST1H3G | 1 |
| 6 | rs7773004 | 26267755 | HIST1H3H | 1 |
| 6 | rs7773004 | 26267755 | HIST1H4B | 1 |
| 6 | rs7773004 | 26267755 | HIST1H4D | 1 |
| 6 | rs7773004 | 26267755 | HIST1H4E | 1 |
| 6 | rs7773004 | 26267755 | HIST1H4F | 1 |
| 6 | rs7773004 | 26267755 | HIST1H4G | 1 |
| 6 | rs7773004 | 26267755 | HIST1H4I | 1 |
| 6 | rs7773004 | 26267755 | HIST1H4J | 1 |
| 6 | rs7773004 | 26267755 | HIST1H4K | 1 |
| 6 | rs7773004 | 26267755 | MAS1L | 1 |
| 6 | rs7773004 | 26267755 | NKAPL | 1 |
| 6 | rs7773004 | 26267755 | OR10C1 | 1 |
| 6 | rs7773004 | 26267755 | OR11A1 | 1 |
| 6 | rs7773004 | 26267755 | OR12D2 | 1 |
| 6 | rs7773004 | 26267755 | OR14J1 | 1 |
| 6 | rs7773004 | 26267755 | OR2B3 | 1 |
| 6 | rs7773004 | 26267755 | OR2B6 | 1 |
| 6 | rs7773004 | 26267755 | OR2H1 | 1 |
| 6 | rs7773004 | 26267755 | OR2J1 | 1 |
| 6 | rs7773004 | 26267755 | OR2J3 | 1 |
| 6 | rs7773004 | 26267755 | OR2W1 | 1 |
| 6 | rs7773004 | 26267755 | SLC17A1 | 1 |
| 6 | rs7773004 | 26267755 | UBD | 1 |

| | | | | | | | | |
|---|---|---|---|---|---|---|---|---|
| | 6 | rs7773004 | 26267755 | ZNF184 | | | | 1 |
| | 6 | rs7773004 | 26267755 | | | | MOG | 1 |
| | 6 | rs7773004 | 26267755 | | | | RP1-265C24.5 | 1 |
| | 6 | rs7773004 | 26267755 | | | | RP5-874C20.3 | 1 |
| Yes | 6 | rs11967262 | 43760327 | VEGFA | | | | 1 |
| | 6 | rs1936800 | 127436064 | RSPO3 | RSPO3 | RSPO3 | | 3 |
| | 6 | rs1936800 | 127436064 | | CENPW | | | 1 |
| | 8 | rs10504825 | 87567848 | CPNE3 | CPNE3 | | | 2 |
| | 8 | rs10504825 | 87567848 | CNGB3 | | | | 1 |
| | 8 | rs10504825 | 87567848 | | RMDN1 | | | 1 |
| Yes | 9 | rs78216177 | 232148 | C9orf66 | | C9orf66 | | 2 |
| Yes | 9 | rs78216177 | 232148 | | CBWD1 | | CBWD1 | 2 |
| Yes | 9 | rs78216177 | 232148 | DOCK8 | | DOCK8 | | 2 |
| Yes | 9 | rs78216177 | 232148 | | FOXD4 | | | 1 |
| Yes | 9 | rs753085 | 117045447 | COL27A1 | | COL27A1 | | 2 |
| | 9 | rs10817762 | 118161597 | DEC1 | | DEC1 | | 2 |
| | 9 | rs10817762 | 118161597 | | TNC | | | 1 |
| Yes | 12 | rs7308356 | 50539611 | AC140061.12 | | AC140061.12 | | 2 |
| Yes | 12 | rs7308356 | 50539611 | ASIC1 | | ASIC1 | | 2 |
| Yes | 12 | rs7308356 | 50539611 | ATF1 | ATF1 | ATF1 | ATF1 | 4 |
| Yes | 12 | rs7308356 | 50539611 | CERS5 | CERS5 | CERS5 | | 3 |
| Yes | 12 | rs7308356 | 50539611 | COX14 | COX14 | COX14 | | 3 |
| Yes | 12 | rs7308356 | 50539611 | DIP2B | | DIP2B | | 2 |
| Yes | 12 | rs7308356 | 50539611 | FAM186A | | FAM186A | | 2 |
| Yes | 12 | rs7308356 | 50539611 | GPD1 | | GPD1 | | 2 |
| Yes | 12 | rs7308356 | 50539611 | LIMA1 | LIMA1 | LIMA1 | | 3 |
| Yes | 12 | rs7308356 | 50539611 | SMARCD1 | | SMARCD1 | | 2 |

| | | | | | | | | |
|---|---|---|---|---|---|---|---|---|
| Yes | 12 | rs7308356 | 50539611 | LARP4 | | | | 1 |
| Yes | 12 | rs7308356 | 50539611 | | METTL7A | | | 1 |
| Yes | 12 | rs7308356 | 50539611 | | | | RP4-605O3.4 | 1 |
| Yes | 12 | rs1054852 | 124496316 | CCDC92 | CCDC92 | CCDC92 | | 3 |
| Yes | 12 | rs1054852 | 124496316 | DNAH10 | DNAH10 | | | 2 |
| Yes | 12 | rs1054852 | 124496316 | DNAH10OS | DNAH10OS | DNAH10OS | DNAH10OS | 4 |
| Yes | 12 | rs1054852 | 124496316 | | | FAM101A | | 1 |
| Yes | 12 | rs1054852 | 124496316 | | | ZNF664 | | 1 |
| Yes | 12 | rs1054852 | 124496316 | | | | RP11-380L11.4 | 1 |
| Yes | 13 | rs41286076 | 73634859 | KLF5 | KLF5 | KLF5 | | 3 |
| Yes | 14 | rs72683923 | 50735947 | CDKL1 | | | | 1 |
| Yes | 14 | rs72683923 | 50735947 | L2HGDH | | | | 1 |
| Yes | 16 | rs11076178 | 57146402 | CPNE2 | CPNE2 | | | 2 |
| | 16 | rs2002833 | 88842117 | APRT | | APRT | | 2 |
| | 16 | rs2002833 | 88842117 | CBFA2T3 | | CBFA2T3 | | 2 |
| | 16 | rs2002833 | 88842117 | CTU2 | CTU2 | CTU2 | | 3 |
| | 16 | rs2002833 | 88842117 | CYBA | CYBA | CYBA | | 3 |
| | 16 | rs2002833 | 88842117 | MVD | | MVD | | 2 |
| | 16 | rs2002833 | 88842117 | PABPN1L | | PABPN1L | | 2 |
| | 16 | rs2002833 | 88842117 | PIEZO1 | PIEZO1 | PIEZO1 | | 3 |
| | 16 | rs2002833 | 88842117 | RNF166 | RNF166 | RNF166 | | 3 |
| | 16 | rs2002833 | 88842117 | RP11-830F9.6 | | RP11-830F9.6 | | 2 |
| | 16 | rs2002833 | 88842117 | SLC22A31 | | SLC22A31 | | 2 |
| | 16 | rs2002833 | 88842117 | SNAI3 | SNAI3 | SNAI3 | | 3 |
| | 16 | rs2002833 | 88842117 | AC092384.1 | | | | 1 |
| | 16 | rs2002833 | 88842117 | CDH15 | | | | 1 |
| | 16 | rs2002833 | 88842117 | CDT1 | | | | 1 |

| | | | | | | | | |
|---|---|---|---|---|---|---|---|---|
| | 16 | rs2002833 | 88842117 | GALNS | | | | 1 |
| | 16 | rs2002833 | 88842117 | IL17C | | | | 1 |
| | 16 | rs2002833 | 88842117 | TRAPPC2L | | | | 1 |
| | 16 | rs2002833 | 88842117 | ZC3H18 | | | | 1 |
| | 16 | rs2002833 | 88842117 | ZFPM1 | | | | 1 |
| | 16 | rs2002833 | 88842117 | | ACSF3 | | | 1 |
| | 16 | rs2002833 | 88842117 | | CDK10 | | | 1 |
| | 16 | rs2002833 | 88842117 | | CHMP1A | | | 1 |
| | 16 | rs2002833 | 88842117 | | DBNDD1 | | | 1 |
| | 16 | rs2002833 | 88842117 | | | | SNAI3-AS1 | 1 |
| | 16 | rs2002833 | 88842117 | | | ANKRD11 | | 1 |
| Yes | 17 | rs6503321 | 2096580 | SMG6 | | SMG6 | | 2 |
| Yes | 17 | rs6503321 | 2096580 | | SRR | SRR | | 2 |
| Yes | 17 | rs6503321 | 2096580 | | SGSM2 | | | 1 |
| | 17 | rs638538 | 68216128 | KCNJ2 | | KCNJ2 | | 2 |
| | 17 | rs638538 | 68216128 | KCNJ16 | | | | 1 |
| | 17 | rs9895127 | 70029808 | AC007461.1 | | AC007461.1 | | 2 |
| Yes | 19 | rs12609241 | 16360926 | AP1M1 | AP1M1 | AP1M1 | AP1M1 | 4 |
| Yes | 19 | rs12609241 | 16360926 | FAM32A | | FAM32A | | 2 |
| Yes | 19 | rs12609241 | 16360926 | KLF2 | | KLF2 | | 2 |
| | 20 | rs3787184 | 50157837 | NFATC2 | | NFATC2 | | 2 |
| Yes | 20 | rs76602912 | 57459868 | GNAS | GNAS | | | 2 |
| | 20 | rs6062619 | 62683002 | OPRL1 | OPRL1 | | | 2 |
| | 20 | rs6062619 | 62683002 | PRPF6 | PRPF6 | | | 2 |
| | 20 | rs6062619 | 62683002 | RGS19 | RGS19 | | | 2 |
| | 20 | rs6062619 | 62683002 | TCEA2 | TCEA2 | TCEA2 | | 3 |
| | 20 | rs6062619 | 62683002 | C20orf201 | | | | 1 |

| | | | | |
|---|---|---|---|---|
| 20 | rs6062619 | 62683002 | SAMD10 | 1 |
| 20 | rs6062619 | 62683002 | SOX18 | 1 |
| 20 | rs6062619 | 62683002 | ZNF512B | 1 |

[a]Denotes the lead SNP at each replicated locus, to which the genes are mapped. [b]Based on NCBI Genome Build 37 (hg19). Highlighted genes (pink) are those that are mapped by more than one gene mapping strategies.

**Supplementary Table 6. Genome-wide gene-based association analysis in MAGMA.** 248 protein-coding genes met the threshold for genome-wide significance ($p<2.67x10^{-6}$, 0.05/18,733) in this analysis. 117 of the 248 genes lay within our replicated loci and are highlighted in red.

| Gene | Chromosome | Number of SNPs | Z-statistic | P-value |
|---|---|---|---|---|
| *PIEZO1* | 16 | 379 | 19.047 | 3.47E-81 |
| *CASZ1* | 1 | 412 | 16.485 | 2.36E-61 |
| *PNO1* | 2 | 51 | 13.673 | 7.33E-43 |
| *RP11-474G23.1* | 2 | 273 | 12.989 | 7.03E-39 |
| *CTU2* | 16 | 69 | 12.911 | 1.95E-38 |
| *PPP3R1* | 2 | 169 | 12.447 | 7.25E-36 |
| *SNAI3* | 16 | 20 | 12.218 | 1.24E-34 |
| *WDR92* | 2 | 66 | 12.123 | 3.97E-34 |
| *EBF1* | 5 | 860 | 10.386 | 1.43E-25 |
| *C3orf27* | 3 | 17 | 10.321 | 2.84E-25 |
| *HDAC7* | 12 | 155 | 9.1745 | 2.27E-20 |
| *MVD* | 16 | 39 | 9.1299 | 3.43E-20 |
| *RNF166* | 16 | 48 | 9.083 | 5.28E-20 |
| *SBF2* | 11 | 1913 | 8.3179 | 4.48E-17 |
| *SLC48A1* | 12 | 94 | 7.6023 | 1.46E-14 |
| *LBH* | 2 | 343 | 7.5722 | 1.84E-14 |
| *AC007461.1* | 17 | 5 | 7.515 | 2.85E-14 |
| *CNRIP1* | 2 | 79 | 7.5049 | 3.07E-14 |
| *NFATC2* | 20 | 499 | 7.5046 | 3.08E-14 |
| *RP11-830F9.6* | 16 | 21 | 7.4466 | 4.79E-14 |
| *DEC1* | 9 | 1014 | 7.3924 | 7.21E-14 |
| *CBFA2T3* | 16 | 482 | 7.3214 | 1.23E-13 |
| *RAPGEF3* | 12 | 132 | 7.2658 | 1.85E-13 |
| *RPN1* | 3 | 196 | 7.1561 | 4.15E-13 |

| | | | | |
|---|---|---|---|---|
| *AMPD3* | 11 | 844 | 7.0672 | 7.90E-13 |
| *GPD1* | 12 | 8 | 6.8552 | 3.56E-12 |
| *SMARCD1* | 12 | 22 | 6.8377 | 4.02E-12 |
| *ATF1* | 12 | 129 | 6.8203 | 4.54E-12 |
| *FGFR3* | 4 | 37 | 6.7879 | 5.69E-12 |
| *OR5V1* | 6 | 332 | 6.7621 | 6.80E-12 |
| *DUSP8* | 11 | 66 | 6.742 | 7.81E-12 |
| *LIMA1* | 12 | 144 | 6.7231 | 8.89E-12 |
| *ZKSCAN8* | 6 | 52 | 6.7014 | 1.03E-11 |
| *HIST1H3J* | 6 | 7 | 6.6892 | 1.12E-11 |
| *CERS5* | 12 | 70 | 6.6467 | 1.50E-11 |
| *UNC5B* | 10 | 314 | 6.6018 | 2.03E-11 |
| *HFE* | 6 | 29 | 6.5637 | 2.62E-11 |
| *TRIM38* | 6 | 46 | 6.5412 | 3.05E-11 |
| *HIST1H4A* | 6 | 1 | 6.5295 | 3.30E-11 |
| *ZNF165* | 6 | 23 | 6.5176 | 3.57E-11 |
| *TBC1D19* | 4 | 218 | 6.5155 | 3.62E-11 |
| *KCNJ2* | 17 | 9 | 6.5148 | 3.64E-11 |
| *SLC12A2* | 5 | 201 | 6.509 | 3.78E-11 |
| *ZKSCAN3* | 6 | 72 | 6.4546 | 5.43E-11 |
| *HIST1H2BN* | 6 | 30 | 6.4141 | 7.08E-11 |
| *ZSCAN9* | 6 | 29 | 6.3914 | 8.22E-11 |
| *UBE2H* | 7 | 470 | 6.3711 | 9.38E-11 |
| *FBLN7* | 2 | 153 | 6.3668 | 9.65E-11 |
| *PINX1* | 8 | 449 | 6.3318 | 1.21E-10 |
| *MSRA* | 8 | 1657 | 6.3275 | 1.25E-10 |
| *SOX7* | 8 | 618 | 6.2959 | 1.53E-10 |

| | | | | |
|---|---|---|---|---|
| *NFATC1* | 18 | 644 | 6.2719 | 1.78E-10 |
| *ZSCAN31* | 6 | 111 | 6.2519 | 2.03E-10 |
| *TRIM27* | 6 | 72 | 6.2462 | 2.10E-10 |
| *SOX7* | 8 | 612 | 6.2358 | 2.25E-10 |
| *PARK7* | 1 | 69 | 6.1989 | 2.84E-10 |
| *TSPAN2* | 1 | 170 | 6.1105 | 4.96E-10 |
| *TCEA2* | 20 | 72 | 6.107 | 5.07E-10 |
| *C9orf66* | 9 | 17 | 6.0944 | 5.49E-10 |
| *ERRFI1* | 1 | 36 | 6.0931 | 5.54E-10 |
| *ARID5B* | 10 | 498 | 6.0863 | 5.78E-10 |
| *PABPN1L* | 16 | 24 | 6.0835 | 5.88E-10 |
| *CABIN1* | 22 | 274 | 6.0765 | 6.14E-10 |
| *FBXO33* | 14 | 96 | 6.0675 | 6.50E-10 |
| *ZSCAN16* | 6 | 10 | 6.0652 | 6.59E-10 |
| *SCAND3* | 6 | 113 | 6.0554 | 7.00E-10 |
| *SHANK3* | 22 | 145 | 5.991 | 1.04E-09 |
| *ZKSCAN4* | 6 | 37 | 5.9893 | 1.05E-09 |
| *OR2B2* | 6 | 4 | 5.9846 | 1.08E-09 |
| *HIST1H4L* | 6 | 1 | 5.9823 | 1.10E-09 |
| *FAM186A* | 12 | 128 | 5.9776 | 1.13E-09 |
| *AFF1* | 4 | 581 | 5.9579 | 1.28E-09 |
| *HIST1H1B* | 6 | 4 | 5.9422 | 1.41E-09 |
| *DIP2B* | 12 | 479 | 5.9002 | 1.82E-09 |
| *COX14* | 12 | 24 | 5.8945 | 1.88E-09 |
| *ZSCAN12* | 6 | 72 | 5.8817 | 2.03E-09 |
| *PROX1* | 1 | 145 | 5.8574 | 2.35E-09 |
| *FAM193A* | 4 | 309 | 5.8466 | 2.51E-09 |

| | | | | |
|---|---|---|---|---|
| *POM121L2* | 6 | 111 | 5.834 | 2.71E-09 |
| *AL031590.1* | 22 | 1 | 5.8168 | 3.00E-09 |
| *GDAP2* | 1 | 53 | 5.8112 | 3.10E-09 |
| *AKAP2* | 9 | 1294 | 5.7984 | 3.35E-09 |
| *PALM2-AKAP2* | 9 | 1295 | 5.7928 | 3.46E-09 |
| *HIST1H2BF* | 6 | 10 | 5.7915 | 3.49E-09 |
| *PRKAR1B* | 7 | 567 | 5.7694 | 3.98E-09 |
| *AC140061.12* | 12 | 5 | 5.7669 | 4.04E-09 |
| *REST* | 4 | 86 | 5.7604 | 4.20E-09 |
| *RAD51B* | 14 | 2377 | 5.7548 | 4.34E-09 |
| *LRRC16A* | 6 | 1556 | 5.7501 | 4.46E-09 |
| *NET1* | 10 | 105 | 5.7492 | 4.48E-09 |
| *KLF5* | 13 | 62 | 5.7389 | 4.76E-09 |
| *ARHGEF26* | 3 | 386 | 5.7251 | 5.17E-09 |
| *CYBA* | 16 | 29 | 5.723 | 5.23E-09 |
| *SMG6* | 17 | 723 | 5.7046 | 5.83E-09 |
| *FAM32A* | 19 | 16 | 5.6733 | 7.00E-09 |
| *HIST1H2BC* | 6 | 32 | 5.6567 | 7.72E-09 |
| *SUSD2* | 22 | 22 | 5.6329 | 8.86E-09 |
| *COL4A2* | 13 | 993 | 5.6315 | 8.93E-09 |
| *BTN3A2* | 6 | 102 | 5.6288 | 9.08E-09 |
| *ZNF664* | 12 | 86 | 5.6287 | 9.08E-09 |
| *HIST1H2AJ* | 6 | 4 | 5.6108 | 1.01E-08 |
| *HIST1H2AL* | 6 | 2 | 5.5903 | 1.13E-08 |
| *ASIC1* | 12 | 50 | 5.5856 | 1.16E-08 |
| *KRTAP5-3* | 11 | 7 | 5.5789 | 1.21E-08 |
| *TSPAN9* | 12 | 986 | 5.5762 | 1.23E-08 |

| | | | | |
|---|---|---|---|---|
| *WDR3* | 1 | 42 | 5.5626 | 1.33E-08 |
| *AMZ1* | 7 | 397 | 5.5364 | 1.54E-08 |
| *BTN1A1* | 6 | 22 | 5.5342 | 1.56E-08 |
| *MOB2* | 11 | 106 | 5.5232 | 1.66E-08 |
| *OR2J2* | 6 | 9 | 5.5048 | 1.85E-08 |
| *HIST1H2BE* | 6 | 2 | 5.5 | 1.90E-08 |
| *HIST1H3C* | 6 | 3 | 5.4988 | 1.91E-08 |
| *SLC17A3* | 6 | 154 | 5.4879 | 2.03E-08 |
| *PGBD1* | 6 | 63 | 5.4846 | 2.07E-08 |
| *LMCD1* | 3 | 247 | 5.4838 | 2.08E-08 |
| *CCBL2* | 1 | 117 | 5.4181 | 3.01E-08 |
| *BTN3A3* | 6 | 36 | 5.411 | 3.13E-08 |
| *MTX1* | 1 | 7 | 5.3996 | 3.34E-08 |
| *FAM13A* | 4 | 1384 | 5.3958 | 3.41E-08 |
| *THBS2* | 6 | 241 | 5.3874 | 3.57E-08 |
| *CUX1* | 7 | 1310 | 5.3848 | 3.63E-08 |
| *CCDC92* | 12 | 124 | 5.3825 | 3.67E-08 |
| *HIST1H2BD* | 6 | 17 | 5.3701 | 3.94E-08 |
| *COG6* | 13 | 410 | 5.3637 | 4.08E-08 |
| *MFAP2* | 1 | 18 | 5.3539 | 4.31E-08 |
| *CTAGE5* | 14 | 362 | 5.3521 | 4.35E-08 |
| *KCNJ12* | 17 | 36 | 5.3512 | 4.37E-08 |
| *ZNF322* | 6 | 73 | 5.3476 | 4.46E-08 |
| *ADM* | 11 | 3 | 5.3421 | 4.59E-08 |
| *TMEM87B* | 2 | 186 | 5.3194 | 5.20E-08 |
| *COL27A1* | 9 | 596 | 5.3097 | 5.49E-08 |
| *RP11-407N17.3* | 14 | 325 | 5.3024 | 5.72E-08 |

| | | | | |
|---|---|---|---|---|
| *RARA* | 17 | 65 | 5.3014 | 5.74E-08 |
| *TBC1D28* | 17 | 23 | 5.2977 | 5.86E-08 |
| *TUBAL3* | 10 | 42 | 5.2903 | 6.10E-08 |
| *SLC17A2* | 6 | 62 | 5.289 | 6.15E-08 |
| *RP1L1* | 8 | 284 | 5.2833 | 6.35E-08 |
| *CA8* | 8 | 360 | 5.2789 | 6.50E-08 |
| *RP1-170O19.20* | 7 | 25 | 5.2752 | 6.63E-08 |
| *NOA1* | 4 | 49 | 5.2526 | 7.50E-08 |
| *ZBTB4* | 17 | 68 | 5.2467 | 7.74E-08 |
| *ANKRD11* | 16 | 472 | 5.2331 | 8.33E-08 |
| *RAPGEFL1* | 17 | 40 | 5.2161 | 9.14E-08 |
| *GOSR2* | 17 | 406 | 5.214 | 9.24E-08 |
| *PALM2* | 9 | 1258 | 5.1994 | 1.00E-07 |
| *HIST1H3I* | 6 | 1 | 5.1993 | 1.00E-07 |
| *RP11-156P1.2* | 17 | 455 | 5.1955 | 1.02E-07 |
| *RERE* | 1 | 804 | 5.1839 | 1.09E-07 |
| *RUNDC1* | 17 | 14 | 5.1838 | 1.09E-07 |
| *HIST1H2BL* | 6 | 2 | 5.1829 | 1.09E-07 |
| *RNF4* | 4 | 403 | 5.1817 | 1.10E-07 |
| *MYOZ2* | 4 | 190 | 5.1805 | 1.11E-07 |
| *BTN2A1* | 6 | 68 | 5.1763 | 1.13E-07 |
| *SNAP29* | 22 | 84 | 5.1668 | 1.19E-07 |
| *ANGPT1* | 8 | 914 | 5.1651 | 1.20E-07 |
| *TMEM106A* | 17 | 13 | 5.1497 | 1.30E-07 |
| *BTN2A2* | 6 | 45 | 5.1485 | 1.31E-07 |
| *ST3GAL4* | 11 | 237 | 5.1481 | 1.32E-07 |
| *GPX6* | 6 | 87 | 5.1471 | 1.32E-07 |

| | | | | |
|---|---|---|---|---|
| *RSPO3* | 6 | 158 | 5.1257 | 1.48E-07 |
| *KCNK13* | 14 | 508 | 5.1217 | 1.51E-07 |
| *SWAP70* | 11 | 351 | 5.1204 | 1.52E-07 |
| *DNAH10OS* | 12 | 15 | 5.1118 | 1.60E-07 |
| *FGD5* | 3 | 562 | 5.1118 | 1.60E-07 |
| *OR12D3* | 6 | 11 | 5.1095 | 1.61E-07 |
| *SRR* | 17 | 37 | 5.1048 | 1.66E-07 |
| *TACC3* | 4 | 155 | 5.1008 | 1.69E-07 |
| *PLXNB2* | 22 | 114 | 5.0811 | 1.88E-07 |
| *ENDOU* | 12 | 45 | 5.0751 | 1.94E-07 |
| *XKR6* | 8 | 1140 | 5.0713 | 1.98E-07 |
| *HMGN4* | 6 | 21 | 5.0702 | 1.99E-07 |
| *THBS3* | 1 | 20 | 5.0687 | 2.00E-07 |
| *HMCN1* | 1 | 700 | 5.0664 | 2.03E-07 |
| *RAB7A* | 3 | 257 | 5.0533 | 2.17E-07 |
| *FAM101A* | 12 | 1137 | 5.049 | 2.22E-07 |
| *RP11-455G16.1* | 4 | 67 | 5.0092 | 2.73E-07 |
| *VAT1* | 17 | 9 | 5.0025 | 2.83E-07 |
| *C8orf74* | 8 | 30 | 5.0021 | 2.84E-07 |
| *MEF2A* | 15 | 608 | 4.9788 | 3.20E-07 |
| *BRSK2* | 11 | 375 | 4.9701 | 3.35E-07 |
| *ARHGAP9* | 12 | 36 | 4.9573 | 3.57E-07 |
| *MSL1* | 17 | 31 | 4.9506 | 3.70E-07 |
| *DOCK10* | 2 | 832 | 4.9459 | 3.79E-07 |
| *BRCA1* | 17 | 182 | 4.9376 | 3.95E-07 |
| *RP11-201K10.3* | 1 | 32 | 4.9296 | 4.12E-07 |
| *C6orf100* | 6 | 4 | 4.9247 | 4.23E-07 |

| | | | | |
|---|---|---|---|---|
| *TNIP2* | 4 | 60 | 4.9221 | 4.28E-07 |
| *CASC3* | 17 | 60 | 4.9163 | 4.41E-07 |
| *ATP13A2* | 1 | 60 | 4.9117 | 4.52E-07 |
| *DNAL1* | 14 | 177 | 4.907 | 4.63E-07 |
| *RASIP1* | 19 | 49 | 4.9067 | 4.63E-07 |
| *PI4KA* | 22 | 471 | 4.9019 | 4.75E-07 |
| *ARL4D* | 17 | 5 | 4.8945 | 4.93E-07 |
| *HOXA9* | 7 | 14 | 4.8941 | 4.94E-07 |
| *PTGES3L* | 17 | 16 | 4.8938 | 4.94E-07 |
| *PTGES3L-AARSD1* | 17 | 39 | 4.8937 | 4.95E-07 |
| *CTD-2267D19.3* | 17 | 11 | 4.8469 | 6.27E-07 |
| *ATOH8* | 2 | 66 | 4.8365 | 6.61E-07 |
| *NBR1* | 17 | 80 | 4.8311 | 6.79E-07 |
| *DOCK8* | 9 | 1349 | 4.8247 | 7.01E-07 |
| *SERINC5* | 5 | 585 | 4.8208 | 7.15E-07 |
| *AARSD1* | 17 | 14 | 4.8196 | 7.19E-07 |
| *PARD3* | 10 | 1802 | 4.81 | 7.55E-07 |
| *RP1-228P16.5* | 12 | 30 | 4.8085 | 7.60E-07 |
| *TOP2A* | 17 | 54 | 4.799 | 7.97E-07 |
| *AP1M1* | 19 | 149 | 4.7902 | 8.33E-07 |
| *CHRNB1* | 17 | 37 | 4.7899 | 8.34E-07 |
| *CFHR2* | 1 | 323 | 4.766 | 9.40E-07 |
| *ABT1* | 6 | 14 | 4.7617 | 9.60E-07 |
| *KLF2* | 19 | 12 | 4.7598 | 9.69E-07 |
| *CDC6* | 17 | 17 | 4.7588 | 9.74E-07 |
| *SCGN* | 6 | 176 | 4.7583 | 9.76E-07 |

| | | | | |
|---|---|---|---|---|
| *MLLT1* | 19 | 212 | 4.743 | 1.05E-06 |
| *HIST1H4C* | 6 | 2 | 4.7365 | 1.09E-06 |
| *RP11-544M22.13* | 1 | 120 | 4.7312 | 1.12E-06 |
| *PKLR* | 1 | 17 | 4.7284 | 1.13E-06 |
| *MIA2* | 14 | 67 | 4.7182 | 1.19E-06 |
| *HIST1H2AC* | 6 | 49 | 4.7167 | 1.20E-06 |
| *ZNF219* | 14 | 31 | 4.7122 | 1.23E-06 |
| *ADAM15* | 1 | 22 | 4.7052 | 1.27E-06 |
| *FUT2* | 19 | 40 | 4.6952 | 1.33E-06 |
| *CAPRIN2* | 12 | 146 | 4.6948 | 1.33E-06 |
| *C15orf52* | 15 | 25 | 4.6931 | 1.35E-06 |
| *SLC17A4* | 6 | 113 | 4.6905 | 1.36E-06 |
| *RP11-321F6.1* | 15 | 467 | 4.6873 | 1.38E-06 |
| *GNA12* | 7 | 381 | 4.6812 | 1.43E-06 |
| *HOXA10* | 7 | 13 | 4.6791 | 1.44E-06 |
| *SERPIND1* | 22 | 49 | 4.6766 | 1.46E-06 |
| *IPO8* | 12 | 223 | 4.6763 | 1.46E-06 |
| *IFI35* | 17 | 14 | 4.6737 | 1.48E-06 |
| *CYSTM1* | 5 | 248 | 4.6721 | 1.49E-06 |
| *GATA2* | 3 | 46 | 4.669 | 1.51E-06 |
| *SLC22A31* | 16 | 22 | 4.6559 | 1.61E-06 |
| *LTBP3* | 11 | 30 | 4.6514 | 1.65E-06 |
| *GJD3* | 17 | 7 | 4.6474 | 1.68E-06 |
| *APRT* | 16 | 7 | 4.6385 | 1.75E-06 |
| *SBSPON* | 8 | 210 | 4.6211 | 1.91E-06 |
| *HIST1H1A* | 6 | 4 | 4.6165 | 1.95E-06 |
| *HCN3* | 1 | 28 | 4.601 | 2.10E-06 |

| | | | | |
|---|---|---|---|---|
| *THAP2* | 12 | 28 | 4.5936 | 2.18E-06 |
| *LGR4* | 11 | 227 | 4.5929 | 2.19E-06 |
| *CTXN3* | 5 | 28 | 4.5903 | 2.21E-06 |
| *AC004466.1* | 12 | 10 | 4.5829 | 2.29E-06 |
| *KRTAP5-5* | 11 | 15 | 4.5664 | 2.48E-06 |
| *SEMA3F* | 3 | 65 | 4.5652 | 2.50E-06 |
| *KRTCAP2* | 1 | 11 | 4.5599 | 2.56E-06 |
| *CLK2* | 1 | 23 | 4.5597 | 2.56E-06 |
| *WNT3* | 17 | 122 | 4.5559 | 2.61E-06 |

**Supplementary Table 7. Summary-based Mendelian Randomisation (SMR) using eQTL data from GTEx v7 tibial artery.** The twenty-seven probes (genes) that met the Bonferroni-corrected significance threshold $P_{SMR} < 1.01\times10^{-5}$ (0.05/4,946) and passed the HEIDI test ($P_{HEIDI} \geq 1.12x10^{-3}$) (0.05/44)) are shown. Fourteen probes mapped to our replicated loci.

| Probe ID[a] | Chr[b] | Gene[c] | Top SNP[d] | EA[e] | NEA[f] | EAF[g] | $P_{GWAS}$[h] | $P_{eQTL}$[i] | $P_{SMR}$[j] | $P_{HEIDI}$[k] |
|---|---|---|---|---|---|---|---|---|---|---|
| ENSG00000123268.4 | 12 | ATF1 | rs1129406 | T | C | 0.41 | $1.70x10^{-13}$ | $2.11x10^{-25}$ | $1.80x10^{-9}$ | $1.68x10^{-2}$ |
| ENSG00000233930.3 | 11 | KRTAP5-AS1 | rs6421044 | T | C | 0.55 | $2.40x10^{-11}$ | $4.50x10^{-30}$ | $8.22x10^{-9}$ | $5.24x10^{-1}$ |
| ENSG00000272368.1 | 12 | RP4-605O3.4 | rs7302981 | G | A | 0.63 | $5.90x10^{-16}$ | -0.0098 | $3.54x10^{-1}$ | 13 |
| ENSG00000245937.3 | 5 | CTC-228N24.3 | rs36694 | A | C | 0.74 | $2.10x10^{-23}$ | $2.81x10^{-11}$ | $3.11x10^{-8}$ | NA |
| ENSG00000244731.3 | 6 | C4A | rs2395149 | A | G | 0.13 | $5.10x10^{-11}$ | $6.64x10^{-17}$ | $2.42x10^{-7}$ | $9.67x10^{-3}$ |
| ENSG00000068078.13 | 4 | FGFR3 | rs3135848 | C | T | 0.27 | $7.60x10^{-11}$ | $1.52x10^{-16}$ | $3.20x10^{-7}$ | $6.14x10^{-1}$ |
| ENSG00000204655.7 | 6 | MOG | rs3129073 | G | A | 0.22 | $1.30x10^{-12}$ | $5.43x10^{-13}$ | $4.25x10^{-7}$ | $7.38x10^{-3}$ |
| ENSG00000198496.6 | 17 | NBR2 | rs1799949 | A | G | 0.33 | $2.00x10^{-7}$ | $1.81x10^{-93}$ | $4.62x10^{-7}$ | $5.44x10^{-2}$ |
| ENSG00000213626.7 | 2 | LBH | rs72787724 | C | T | 0.21 | $9.30x10^{-14}$ | $7.71x10^{-11}$ | $9.55x10^{-7}$ | $9.10x10^{-1}$ |
| ENSG00000228078.1 | 6 | HLA-U | rs1632933 | T | C | 0.46 | $1.20x10^{-9}$ | $2.00x10^{-16}$ | $1.01x10^{-6}$ | $1.65x10^{-2}$ |
| ENSG00000233327.6 | 17 | USP32P2 | rs9909256 | T | G | 0.52 | $3.10x10^{-7}$ | $1.69x10^{-53}$ | $1.20x10^{-6}$ | $5.19x10^{-1}$ |
| ENSG00000273018.1 | 17 | CTD-2303H24.2 | rs9909256 | T | G | 0.52 | $3.10x10^{-7}$ | $2.9x10^{-3}$ | $4.61x10^{-1}$ | 16 |
| ENSG00000137944.12 | 1 | CCBL2 | rs3820242 | G | A | 0.50 | $2.20x10^{-7}$ | $5.81x10^{-36}$ | $1.68x10^{-6}$ | $1.07x10^{-2}$ |
| ENSG00000260630.2 | 16 | SNAI3-AS1 | rs6500487 | T | C | 0.62 | $5.30x10^{-11}$ | $1.0x10^{-2}$ | $3.50x10^{-1}$ | 4 |
| ENSG00000250091.2 | 12 | DNAH10OS | rs4765127 | T | G | 0.33 | $1.30x10^{-7}$ | $6.4x10^{-3}$ | $9.89x10^{-2}$ | 9 |
| ENSG00000213077.5 | 17 | FAM106A | rs9909256 | T | G | 0.52 | $3.10x10^{-7}$ | $1.53x10^{-39}$ | $1.86x10^{-6}$ | $5.32x10^{-1}$ |
| ENSG00000219392.1 | 6 | RP1-265C24.5 | rs2275508 | C | T | 0.24 | $1.30x10^{-9}$ | $4.54x10^{-14}$ | $2.27x10^{-6}$ | $6.35x10^{-2}$ |
| ENSG00000144152.8 | 2 | FBLN7 | rs10175143 | G | A | 0.49 | $1.20x10^{-7}$ | $9.33x10^{-25}$ | $2.53x10^{-6}$ | $2.39x10^{-1}$ |
| ENSG00000197062.7 | 6 | RP5-874C20.3 | rs1531681 | A | G | 0.58 | $1.90x10^{-8}$ | $1.0x10^{-2}$ | $1.14x10^{-2}$ | 20 |
| ENSG00000243667.2 | 2 | WDR92 | rs12994861 | T | C | 0.27 | $6.70x10^{-15}$ | $5.85x10^{-9}$ | $3.11x10^{-6}$ | $9.63x10^{-3}$ |
| ENSG00000220161.4 | 17 | RP1-37N7.3 | rs9909256 | T | G | 0.52 | $3.10x10^{-7}$ | -0.0039 | $2.87x10^{-1}$ | 11 |
| ENSG00000172785.14 | 9 | CBWD1 | rs522747 | T | A | 0.41 | $5.00x10^{-7}$ | $2.83x10^{-27}$ | $5.12x10^{-6}$ | $8.21x10^{-2}$ |
| ENSG00000270028.1 | 12 | RP11-380L11.4 | rs4765127 | T | G | 0.33 | $1.30x10^{-7}$ | $6.7x10^{-3}$ | $5.12x10^{-2}$ | 7 |

| | | | | | | | | | | |
|---|---|---|---|---|---|---|---|---|---|---|
| ENSG00000228022.1 | 6 | HCG20 | rs3131043 | G | A | 0.42 | 3.60x10$^{-8}$ | 2.74x10$^{-14}$ | 8.12x10$^{-6}$ | 5.31x10$^{-2}$ |
| ENSG00000146112.7 | 6 | PPP1R18 | rs3129973 | T | C | 0.15 | 4.30x10$^{-10}$ | 2.40x10$^{-10}$ | 8.74x10$^{-6}$ | 8.59x10$^{-2}$ |
| ENSG00000072958.4 | 19 | AP1M1 | rs4808034 | A | G | 0.72 | 3.10x10$^{-8}$ | 1.68x10$^{-13}$ | 9.60x10$^{-6}$ | NA |
| ENSG00000133103.12 | 13 | COG6 | rs4368051 | T | C | 0.75 | 1.60x10$^{-6}$ | 7.71x10$^{-30}$ | 9.98x10$^{-6}$ | 2.94x10$^{-1}$ |

[a]Probe ID. [b]Probe chromosome. [c]Gene name. [d]SNP name. [e]Effect allele. [f]Non-effect allele. [g]Frequency of the effect allele in the study population. [h]GWAS P-value. [i]eQTL P-value. [j]SMR P-value. [k]HEIDI P-value. Probes mapped to replicated loci are highlighted in red.

**Supplementary Table 8. Enriched gene sets from genome-wide gene-based enrichment analysis in MAGMA v1.07.** The convergence of 15,496 gene sets (15,381from MSigDB v7.0) were tested (See Supplementary Data 3 for all tested gene sets). A Bonferroni-corrected threshold of $P < 3.23\times10^{-6}$ (0.05/15,496) was set, resulting in four significant Gene Ontology (GO) gene sets and two curated gene sets. This analysis was performed using the SNP2GENE tool in FUMA.

| Gene Set | $N_{genes}$ | Beta | $Beta_{STD}$ | SE | P-value | $P_{bon}$ |
|---|---|---|---|---|---|---|
| GO_bp:go_cardiovascular_system_development | 666 | 0.22331 | 0.041351 | 0.039951 | $1.16x10^{-8}$ | $1.79x10^{-4}$ |
| GO_bp:go_tube_morphogenesis | 778 | 0.1964 | 0.039186 | 0.037667 | $9.35x10^{-8}$ | $1.45x10^{-3}$ |
| GO_bp:go_blood_vessel_morphogenesis | 555 | 0.21137 | 0.03584 | 0.044333 | $9.39x10^{-7}$ | $1.45x10^{-2}$ |
| GO_bp:go_tube_development | 956 | 0.15781 | 0.03473 | 0.033944 | $1.68x10^{-6}$ | $2.60x10^{-2}$ |
| Curated_gene_sets:nikolsky_breast_cancer_16q24_amplicon | 53 | 1.7715 | 0.094095 | 0.25379 | $1.53x10^{-12}$ | $2.37x10^{-8}$ |
| Curated_gene_sets:cui_tcf21_targets_2_dn | 786 | 0.18957 | 0.038009 | 0.03709 | $1.62x10^{-7}$ | $2.51x10^{-3}$ |

**Supplementary Table 9. Genetic correlation between varicose veins and other phenotypes.** This analysis was performed using LD score (LDSC) regression, implemented in LD Hub. The traits are shown along with the consortia name, sample size, ethnicity, and PMID of the study from which the LDSC data were derived, the trait category, and the correlation coefficient ($r_g$). Traits are ranked by P-value, and the twelve traits meeting a Bonferroni-corrected significant threshold of $P < 5.56\times10^{-3}$ are shown.

| Trait 1 | Trait 2 | Category | $r_g$ | SE | Z-score | P-value | Consortia | Sample Size | Ethnicity | PMID |
|---|---|---|---|---|---|---|---|---|---|---|
| Varicose veins | Height_2010 | Anthropometric | 0.16 | 0.03 | 5.69 | $1.28\times10^{-8}$ | GIANT | 133,859 | European | 20881960 |
| Varicose veins | Height; Females at age 10 and males at age 12 | Anthropometric | 0.21 | 0.05 | 4.63 | $3.59\times10^{-6}$ | EGG | 13,960 | European | 23449627 |
| Varicose veins | Extreme height | Anthropometric | 0.17 | 0.04 | 4.51 | $6.36\times10^{-6}$ | GIANT | 16,196 | European | 23563607 |
| Varicose veins | Hip circumference | Anthropometric | 0.13 | 0.03 | 4.47 | $7.72\times10^{-6}$ | GIANT | 213,038 | European | 25673412 |
| Varicose veins | Waist circumference | Anthropometric | 0.10 | 0.03 | 3.80 | $1.00\times10^{-4}$ | GIANT | 232,101 | European | 25673412 |
| Varicose veins | Child birth weight | Anthropometric | 0.21 | 0.06 | 3.77 | $2.00\times10^{-4}$ | EGG | 26,836 | European | 23202124 |
| Varicose veins | Own birth weight | Anthropometric | 0.11 | 0.03 | 3.71 | $2.00\times10^{-4}$ | Warrington EGG | 321,223 | Mixed | 31043758 |
| Varicose veins | Offspring birth weight | Anthropometric | 0.11 | 0.03 | 3.71 | $2.00\times10^{-4}$ | Warrington EGG | 230,069 | Mixed | 31043758 |
| Varicose veins | Own birth weight | Anthropometric | 0.11 | 0.03 | 3.64 | $3.00\times10^{-4}$ | Warrington EGG | 286,870 | European | 31043758 |
| Varicose veins | Birth weight | Anthropometric | 0.12 | 0.04 | 3.25 | $1.20\times10^{-3}$ | NA | 143,677 | European | 27680694 |
| Varicose veins | Systemic lupus erythematosus | Autoimmune | 0.19 | 0.06 | 3.17 | $1.50\times10^{-3}$ | Alkes Group | 23,210 | European | 26502338 |
| Varicose veins | Body mass index | Anthropometric | 0.09 | 0.03 | 2.87 | $4.20\times10^{-3}$ | GIANT | 123,912 | European | 20935630 |

**Supplementary Table 10. Enriched drug pathways from the drug target enrichment analysis.** Mapped genes were interrogated with the Open Targets Platform to enrich drug pathways relating to the identified gene targets. The 200 gene targets identified by the Open Targets Platform mapped to 622 drug pathways, of which, 42 reached a nominal significance P < 0.05 and are shown below.

| Pathway | P-value | Number of targets | Targets in pathway |
|---|---|---|---|
| Butyrophilin (BTN) family interactions | 2.60E-07 | 6 | *BTN1A1, BTN2A1, BTN3A1, BTN2A2, BTN3A2, BTN3A3* |
| Calcineurin activates NFAT | 0.00094 | 2 | *NFATC2, PPP3R1* |
| Transcriptional regulation by RUNX1 | 0.0016 | 14 | *H4C2, RPN1, H2BC4, SGSM2, HIST1H3C, H3C6, HIST1H3B, ZFPM1, RSPO3, CHMP1A, NFATC2, EBF1, SMARCD1, GATA2* |
| CLEC7A (Dectin-1) induces NFAT activation | 0.0017 | 2 | *NFATC2, PPP3R1* |
| Vitamin B1 (thiamin) metabolism | 0.005 | 2 | *SMARCD1, CARMIL1* |
| Transcriptional regulation of granulopoiesis | 0.0083 | 7 | *H4C2, H2BC4, GATA2, KLF5, HIST1H3C, H3C6, HIST1H3B* |
| HDACs deacetylate histones | 0.0088 | 7 | *H4C2, H2BC4, H2AC11, REST, HIST1H3C, H3C6, HIST1H3B* |
| Detoxification of Reactive Oxygen Species | 0.0094 | 3 | *GPX5, GPX6, CYBA* |
| VEGF ligand-receptor interactions | 0.012 | 1 | *VEGFA* |
| VEGF binds to VEGFR leading to receptor dimerization | 0.012 | 1 | *VEGFA* |
| RUNX1 regulates genes involved in megakaryocyte differentiation and platelet function | 0.013 | 7 | *CHMP1A, H4C2, H2BC4, HIST1H3C, H3C6, HIST1H3B, ZFPM1* |
| Assembly of the pre-replicative complex | 0.014 | 4 | *RPN1, SGSM2, CDT1, MCM2* |
| G protein gated Potassium channels | 0.016 | 3 | *KCNJ16, GABBR1, KCNJ2* |
| Activation of G protein gated Potassium channels | 0.016 | 3 | *KCNJ16, GABBR1, KCNJ2* |
| Inhibition of voltage gated Ca2+ channels via Gbeta/gamma subunits | 0.016 | 3 | *KCNJ16, GABBR1, KCNJ2* |

| | | | |
|---|---|---|---|
| Interleukin-7 signaling | 0.016 | 4 | *L2HGDH, HIST1H3C, H3C6, HIST1H3B* |
| Orc1 removal from chromatin | 0.017 | 4 | *RPN1, SGSM2, CDT1, MCM2* |
| RUNX1 regulates transcription of genes involved in differentiation of HSCs | 0.018 | 8 | *H4C2, RPN1, H2BC4, SGSM2, GATA2, HIST1H3C, H3C6, HIST1H3B* |
| RMTs methylate histone arginines | 0.02 | 6 | *H4C2, H2AC11, SMARCD1, HIST1H3C, H3C6, HIST1H3B* |
| Defective SLC34A1 causes hypophosphatemic nephrolithiasis/osteoporosis 1 (NPHLOP1) | 0.021 | 1 | *SLC17A2* |
| Defective SLC4A1 causes hereditary spherocytosis type 4 (HSP4), distal renal tubular acidosis (dRTA) and dRTA with hemolytic anemia (dRTA-HA) | 0.021 | 1 | *SMARCD1* |
| MPS IV - Morquio syndrome A | 0.021 | 1 | *GALNS* |
| Unwinding of DNA | 0.026 | 1 | *MCM2* |
| Inwardly rectifying K+ channels | 0.028 | 3 | *KCNJ16, GABBR1, KCNJ2* |
| Downstream signaling events of B Cell Receptor (BCR) | 0.029 | 4 | *NFATC2, RPN1, SGSM2, PPP3R1* |
| RNA Polymerase I Promoter Opening | 0.03 | 5 | *H4C2, H2BC4, HIST1H3C, H3C6, HIST1H3B* |
| Activation of the pre-replicative complex | 0.032 | 2 | *CDT1, MCM2* |
| DNA Replication Pre-Initiation | 0.034 | 4 | *RPN1, SGSM2, CDT1, MCM2* |
| DNA methylation | 0.035 | 5 | *H4C2, H2BC4, HIST1H3C, H3C6, HIST1H3B* |
| B-WICH complex positively regulates rRNA expression | 0.038 | 6 | *H4C2, H2BC4, POLR1B, HIST1H3C, H3C6, HIST1H3B* |
| RNA Polymerase I Promoter Escape | 0.038 | 6 | *H4C2, H2BC4, POLR1B, HIST1H3C, H3C6, HIST1H3B* |
| Formation of the beta-catenin:TCF transactivating complex | 0.038 | 6 | *CHMP1A, H4C2, H2BC4, HIST1H3C, H3C6, HIST1H3B* |

| | | | |
|---|---|---|---|
| Olfactory Signaling Pathway | 0.038 | 15 | *OR2H1, OR2J3, OR2H2, OR2J1, OR2J2, OR2B3, OR2B6, OR2B2, OR12D2, OR2W1, OR12D3, OR11A1, OR10C1, OR14J1, OR5V1* |
| Activated PKN1 stimulates transcription of AR (androgen receptor) regulated genes KLK2 and KLK3 | 0.04 | 5 | *H4C2, H2BC4, HIST1H3C, H3C6, HIST1H3B* |
| Biogenic amines are oxidatively deaminated to aldehydes by MAOA and MAOB | 0.041 | 1 | *KLF2* |
| Switching of origins to a post-replicative state | 0.041 | 4 | *RPN1, SGSM2, CDT1, MCM2* |
| Activation of HOX genes during differentiation | 0.043 | 7 | *CHMP1A, H4C2, POLR2B, H2BC4, HIST1H3C, H3C6, HIST1H3B* |
| Activation of anterior HOX genes in hindbrain development during early embryogenesis | 0.043 | 7 | *CHMP1A, H4C2, POLR2B, H2BC4, HIST1H3C, H3C6, HIST1H3B* |
| SIRT1 negatively regulates rRNA expression | 0.043 | 5 | *H4C2, H2BC4, HIST1H3C, H3C6, HIST1H3B* |
| Oxidative Stress Induced Senescence | 0.048 | 7 | *H4C2, EBF1, H2BC4, MAPK10, HIST1H3C, H3C6, HIST1H3B* |
| GABA B receptor activation | 0.049 | 3 | *KCNJ16, GABBR1, KCNJ2* |
| Activation of GABAB receptors | 0.049 | 3 | *KCNJ16, GABBR1, KCNJ2* |

**Supplementary Table 11. Tractability information for targets in the drug-target enrichment analysis.** The 237 mapped genes were interrogated within the Open Targets Platform to determine their tractability to small molecule and antibody targeting. 200 genes targets were identified by the Open Targets Platform of which, tractability information was available for 105 gene targets (left column).

| | **Small molecule** | | | **Antibody** | | |
|---|---|---|---|---|---|---|
| **Gene Target** | **Clinical precedence** | **Discovery precedence** | **Predicted tractable** | **Clinical precedence** | **Predicted tractable high confidence** | **Predicted tractable mid-low confidence** |
| *ANKRD11* | | | | | Yes | |
| *AP1M1* | | | | | Yes | |
| *APRT* | | Yes | Yes | | Yes | |
| *ASIC1* | | Yes | Yes | | Yes | Yes |
| *ATF1* | | | Yes | | | |
| *BCAS2* | | Yes | | | | |
| *BTN1A1* | | | | | Yes | Yes |
| *BTN2A1* | | | | | Yes | Yes |
| *BTN2A2* | | | | | Yes | Yes |
| *BTN3A1* | | Yes | | | Yes | Yes |
| *BTN3A2* | | | | | Yes | Yes |
| *BTN3A3* | | Yes | | | Yes | Yes |
| *C3orf20* | | | | | | Yes |
| *CARMIL1* | | | | | Yes | |
| *CDH15* | | | | | Yes | Yes |
| *CDK10* | Yes | | Yes | | | |
| *CDKL1* | | Yes | Yes | | | |
| *CERS5* | | | | | | Yes |
| *CHMP1A* | | Yes | | | | |

| | | | | | | |
|---|---|---|---|---|---|---|
| *CNGB3* | | | Yes | | | Yes |
| *CNRIP1* | | | | | Yes | |
| *COL27A1* | | | | | Yes | Yes |
| *COX14* | | | | | | Yes |
| *CPNE2* | | | | | | Yes |
| *CPNE3* | | | | | Yes | |
| *CTXN3* | | | | | | Yes |
| *CYBA* | | | | | Yes | |
| *DOCK8* | | | | | Yes | |
| *EBF1* | | Yes | | | | |
| *EFEMP1* | | | | | Yes | Yes |
| *FAM32A* | | Yes | | | | |
| *FBLN7* | | | | | | Yes |
| *FBN2* | | | | | Yes | Yes |
| *FGD5* | | | | | Yes | Yes |
| *GABBR1* | Yes | Yes | Yes | | Yes | Yes |
| *GALNS* | | Yes | | | Yes | Yes |
| *GNAS* | | Yes | | | Yes | |
| *GPD1* | | Yes | | | | |
| *GPX5* | | | | | Yes | Yes |
| *GPX6* | | | | | | Yes |
| *H1-4* | | Yes | | | | |
| *H1-5* | | Yes | | | | |
| *H2AC8* | | Yes | | | | |
| *H2BC11* | | Yes | | | Yes | |

| | | | | | | |
|---|---|---|---|---|---|---|
| *H3C6* | | Yes | | | Yes | |
| *H4C2* | | Yes | | | Yes | |
| *HFE* | | | | | Yes | Yes |
| *IGFBP7* | | | | | Yes | Yes |
| *IL17C* | | | | | Yes | Yes |
| *INSIG2* | | | | | | Yes |
| *KCNJ16* | | | Yes | | Yes | Yes |
| *KCNJ2* | Yes | | Yes | | Yes | Yes |
| *KLF5* | | | Yes | | | |
| *LIMA1* | | | | | Yes | Yes |
| *LPP* | | | | | | Yes |
| *MAPK10* | Yes | Yes | Yes | | | Yes |
| *MAS1L* | | | Yes | | Yes | Yes |
| *MCM2* | | Yes | | | | |
| *METTL7A* | | | | | Yes | Yes |
| *MOG* | | | | | Yes | Yes |
| *MVD* | | | Yes | | | |
| *OPRL1* | Yes | Yes | Yes | | Yes | Yes |
| *OR10C1* | | | | | Yes | Yes |
| *OR11A1* | | | Yes | | Yes | Yes |
| *OR12D2* | | | | | Yes | Yes |
| *OR12D3* | | | | | Yes | Yes |
| *OR14J1* | | | Yes | | Yes | Yes |
| *OR2B2* | | | | | Yes | Yes |
| *OR2B3* | | | | | Yes | Yes |

| | | | | | |
|---|---|---|---|---|---|
| *OR2B6* | | | | Yes | Yes |
| *OR2H1* | | Yes | | Yes | Yes |
| *OR2H2* | | Yes | | Yes | Yes |
| *OR2J1* | | | | Yes | Yes |
| *OR2J2* | | | | Yes | Yes |
| *OR2J3* | | | | Yes | Yes |
| *OR2W1* | | | | Yes | Yes |
| *OR5V1* | | | | Yes | Yes |
| *PIEZO1* | | | | Yes | Yes |
| *PLXND1* | Yes | | | Yes | Yes |
| *POLR2B* | Yes | | | | |
| *PPP3R1* | Yes | Yes | | Yes | Yes |
| *PRPF6* | Yes | | | | |
| *PRSS16* | | | | | Yes |
| *RAB7A* | Yes | Yes | | Yes | |
| *RGS19* | | | | Yes | |
| *RPN1* | | | | | Yes |
| *RSPO3* | | | | Yes | Yes |
| *SCGN* | | | | | Yes |
| *SLC12A2* | | Yes | | Yes | Yes |
| *SLC17A1* | | | | Yes | Yes |
| *SLC17A2* | | | | | Yes |
| *SLC17A3* | | | | Yes | Yes |
| *SLC17A4* | | | | Yes | Yes |
| *SLC22A31* | | Yes | | | Yes |

| | | | | | |
|---|---|---|---|---|---|
| *SMG6* | | Yes | | | |
| *SORBS2* | | | | | Yes |
| *SRR* | | Yes | | | Yes |
| *SYCP1* | Yes | | | | |
| *TMEM87B* | | | | | Yes |
| *TNC* | | | Yes | Yes | Yes |
| *TSHB* | | | | Yes | Yes |
| *TSPAN2* | | | | | Yes |
| *VEGFA* | Yes | Yes | Yes | Yes | Yes |
| *WDR92* | | Yes | | | |
| *ZFPM1* | Yes | | | | |

**Supplementary Table 12. Pharmacologically-active targets identified in drug-target enrichment** Of the 200 varicose-veins associated genes targets identified by the Open Targets Platform, eight gene targets have known pharmaceutical interactions and are presently, or in the past have been, investigated in clinical trials in different phases for the treatment of several diseases. Diseases shown below are those relating to vascular disorders.

| Gene target | Drug | Type | Maximum Phase | Mechanism of action | Target | Activity | Disease |
|---|---|---|---|---|---|---|---|
| *CDK10* | Roniciclib | Small molecule | 2 | Cyclin-dependent kinase inhibitor | Cyclin-dependent kinase | Negative modulator | - |
| *CDK10* | AT-7519 | Small molecule | 2 | Cyclin-dependent kinase inhibitor | Cyclin-dependent kinase | Negative modulator | - |
| *CDK10* | PHA-793887 | Small molecule | 1 | Cyclin-dependent kinase inhibitor | Cyclin-dependent kinase | Negative modulator | - |
| *CDK10* | AZD-5438 | Small molecule | 1 | Cyclin-dependent kinase inhibitor | Cyclin-dependent kinase | Negative modulator | - |
| *COL27A1* | Collagenase Clostridium Histolyticum | Enzyme | 4 | Collagen hydrolytic enzyme | Collagen | Other | Abnormality of connective tissue, decubitus ulcer, diabetic foot, skin ulcer, skin wound |
| *COL27A1* | Ocriplasmin | Enzyme | 4 | Collagen hydrolytic enzyme | Collagen | Other | Deep vein thrombosis, diabetic macular oedema, retinal vein occlusion, stroke |
| *GABBR1* | Baclofen | Small molecule | 4 | GABA-B receptor agonist | GABA-B receptor | Positive modulator | - |
| *GABBR1* | Oxybate | Small molecule | 4 | GABA-B receptor agonist | GABA-B receptor | Positive modulator | - |
| *GABBR1* | Arbaclofen Placarbil | Small molecule | 3 | GABA-B receptor agonist | GABA-B receptor | Positive modulator | - |
| *GABBR1* | Arbaclofen | Small molecule | 3 | GABA-B receptor agonist | GABA-B receptor | Positive modulator | - |
| *GABBR1* | Lesogaberan | Small molecule | 2 | GABA-B receptor 1 agonist | GABA-B receptor 1 agonist | Positive modulator | - |

| | | | | | | | |
|---|---|---|---|---|---|---|---|
| *KCNJ2* | Dronedarone | Small molecule | 4 | Inward rectifier potassium channel 2 blocker | Inward rectifier potassium channel 2 | Negative modulator | - |
| *MAPK10* | Tanzisertib | Small molecule | 2 | c-Jun N-terminal kinase 3 inhibitor | c-Jun N-terminal kinase 3 | Negative modulator | Systemic lupus erythematosus, idiopathic pulmonary fibrosis |
| *MAPK10* | CC-401 | Small molecule | 2 | c-Jun N-terminal kinase 3 inhibitor | c-Jun N-terminal kinase 3 | Negative modulator | - |
| *OPRL1* | LY2940094 | Small molecule | 2 | Nociceptin receptor antagonist | Nociceptin receptor | Negative modulator | - |
| *OPRL1* | Cebranopadol | Small molecule | 3 | Nociceptin receptor agonist | Nociceptin receptor | Positive modulator | Pain, diabetic neuropathy |
| *TNC* | F16il2 | Antibody | 2 | Tenascin other | Tenascin | Other | - |
| *TNC* | 81C6 131I | Antibody | 2 | Tenascin binding agent | Tenascin | Other | - |
| *TNC* | Tenatumomab | Antibody | 2 | Tenascin other | Tenascin | Other | - |
| *TNC* | F16SIP 131I | Antibody | 2 | Tenascin binding agent | Tenascin | - | - |
| *VEGFA* | Aflibercept | Protein | 4 | Vascular endothelial growth factor A inhibitor | Vascular endothelial growth factor A | Negative modulator | Age-related macular degeneration,, neovascular glaucoma, ocular vascular disease, proliferative diabetic retinopathy, vitreous haemorrhage, wet macular degeneration |
| *VEGFA* | Ranibizumab | Antibody | 4 | Vascular endothelial growth factor A inhibitor | Vascular endothelial growth factor A | Negative modulator | Age-related macular degeneration, diabetes mellitus, diabetic macular oedema, diabetic retinopathy, neovascular glaucoma, ocular vascular disease, proliferative diabetic retinopathy, retinal detachment, retinal vein occlusion, wet macular degeneration |
| *VEGFA* | Bevacizumab | Antibody | 4 | Vascular endothelial growth factor A inhibitor | Vascular endothelial growth factor A | Negative modulator | Age-related macular degeneration, angiosarcoma, anterior ischemic optic neuropathy, corneal neovascularization, diabetic macular oedema, diabetic retinopathy, Hereditary haemorrhagic telangiectasia, proliferative diabetic |

| | | | | | | | |
|---|---|---|---|---|---|---|---|
| | | | | | | | retinopathy, Pulmonary venoocclusive disease, retinal detachment, retinal vein occlusion, vascular disease, vitreous haemorrhage, wet macular degeneration |
| *VEGFA* | Pegaptanib Sodium | Unknown | 4 | Vascular endothelial growth factor A antagonist | Vascular endothelial growth factor A | Negative modulator | Age-related macular degeneration, diabetic macular oedema, ocular vascular disease, proliferative diabetic retinopathy, retinal vein occlusion |
| *VEGFA* | Conbercept | Protein | 3 | Vascular endothelial growth factor A inhibitor | Vascular endothelial growth factor A | Negative modulator | Age-related macular degeneration, diabetic macular oedema, proliferative diabetic retinopathy, retinal vein occlusion, vitreous haemorrhage |
| *VEGFA* | Muparfostat | Oligosaccharide | 3 | Vascular endothelial growth factor A inhibitor | Vascular endothelial growth factor A | Negative modulator | - |
| *VEGFA* | Brolucizumab | Antibody | 3 | Vascular endothelial growth factor A inhibitor | Vascular endothelial growth factor A | Negative modulator | Age-related macular degeneration, diabetic macular oedema, retinal vein occlusion |
| *VEGFA* | Vanucizumab | Antibody | 2 | Vascular endothelial growth factor A inhibitor | Vascular endothelial growth factor A | Negative modulator | - |

**Supplementary Table 13. Functional categories of the gene clusters.** The varicose veins susceptibility loci map to genes implicated in five functional categories. Several genes map to more than one category. Genes not associated with varicose veins previously are highlighted in bold.

| Biological Process | Gene | Cytogenetic location | OMIM Entry | Function | References |
|---|---|---|---|---|---|
| **Angiogenesis and Lymphangiogenesis** | Castor Zinc Finger Protein 1 (CASZ1) | 1p36.22 | 609895 | Involved in vascular formation and sprouting. | Charpentier MS. et al. CASZ1 promotes vascular assembly and morphogenesis through the direct regulation of an EGFL7/RhoA-mediated pathway. Dev Cell. 2013;25: 132–143. (2013) |
| | Fibulin 7 (FBLN7) | 2q13-q14 | 611551 | Secreted glycoprotein that interacts with extracellular matrix proteins, it is expressed in blood vessels with the c-terminal fragment (FBLN7-C) known to demonstrate anti-angiogenic activity and disrupt tube formation and vessel sprouting in human vein. | De Vega, S. et al. A C-terminal fragment of fibulin-7 interacts with endothelial cells and inhibits their tube formation in culture. Arch. Biochem. Biophys. 545, 148–153 (2014). |
| | **FYVE, Rhogef, and PH Domain-containing Protein 5 (FGD5)** | **3p25.1** | **614788** | **Necessary for angiogenesis through its critical role in endothelial cell responses to VEGF stimulation- and additionally prevents against pro-apoptotic stresses in the endothelium.** | **Nakhaei-Nejad. et al. Q.-X. & Murray, A. G. Facio-genital dysplasia-5 regulates matrix adhesion and survival of human endothelial cells. Arterioscler. Thromb. Vasc. Biol. 32, 2694–701 (2012).** |
| | GATA-Binding Protein 2 (GATA2) | 3q21.3 | 137295 | Involved in VEGF-induced angiogenesis and lymphangiogenesis, it is necessary for vascular and lymphovascular integrity. | Spinner, M. A. et al. GATA2 deficiency: A protean disorder of hematopoiesis, lymphatics, and immunity. Blood 123, 809–821 (2014). |
| | **Insulin-Like Growth Factor-Binding Protein 7; IGFBP7** | **4q12** | **60286** | **Blocks VEGF-induced angiogenesis through interfering with VEGF expression in human endothelial cells. IGFBP7** | **Tamura, K. et al. Insulin-like growth factor binding protein-7 (IGFBP7) blocks vascular endothelial cell growth factor (VEGF)-induced angiogenesis in human vascular endothelial cells. Eur. J. Pharmacol. 610, 61–67 (2009).** |

| | | | | |
|---|---|---|---|---|
| | | | **mutations result in familial retinal arterial macro-aneurysm (FRAM).** | **Abu-Safieh, L. et al. Mutation of IGFBP7 causes upregulation of BRAF/MEK/ERK pathway and familial retinal arterial macroaneurysms. Am. J. Hum. Genet. 89, 313–319 (2011).** |
| **Kruppel-Like Factor 2 (Klf2)** | **19p13.11** | **602016** | **Encodes a shear-responsive transcription factor that is a key inducer of endothelial gene expression in response to fluid shear stress.** | **Dekker, R. J. et al. Prolonged fluid shear stress induces a distinct set of endothelial cell genes, most specifically lung Krüppel-like factor (KLF2). Blood 100, 1689–1698 (2002).** |
| Piezo-Type Mechanosensitive Ion Channel Component 1 (PIEZO1) | 16q24.3 | 611184 | Encodes a mechanosensitive cation channel induced by fluid shear stress and membrane stretch. Involved in regulating vascular architecture and arterial remodelling. It also plays a significant role in the normal function of the lymphatic system. | Li J, Hou B, Tumova S, Muraki K, Bruns A, Ludlow MJ, et al. Piezo1 integration of vascular architecture with physiological force. Nature. 2014;515: 279–282. doi:10.1038/nature13701<br>Fotiou, E. et al. Novel mutations in PIEZO1 cause an autosomal recessive generalized lymphatic dysplasia with non-immune hydrops fetalis. Nat. Commun. 6, 1–6 (2015). |
| **Prospero-Related Homeobox 1 (PROX1)** | **1q32.3** | **601546** | **Master inducer of lymphatic vasculature, necessary for lymphovenous valve development.** | **Bazigou, E. et al. Genes regulating lymphangiogenesis control venous valve formation and maintenance in mice. J. Clin. Invest. 121, 2984–2992 (2011).**<br>**Sathish Srinivasan, R. & Oliver, G. Prox1 dosage controls the number of lymphatic endothelial cell progenitors and the formation of the lymphovenous valves. Genes Dev. 25, 2187–2197 (2011).** |
| R-Spondin 3 (RSPO3) | 6q22.33 | 610574 | A WNT signalling enhancer that is highly expressed in vasculature. Endothelial RSPO3 controls vascular pruning and stability through involvement in a critical maintenance pathway in remodelling vasculature. | Scholz B. et al. Endothelial RSPO3 Controls Vascular Stability and Pruning through Non-canonical WNT/Ca(2+)/NFAT Signaling. Dev Cell. 2016;36(1):79–93 (2015) |
| SRY-Box 18 (SOX18) | 20q13.33 | 601618 | Encodes a protein involved the development of lymphatic vasculature, with SOX18 mutations resulting in hypotrichosis-lymphoedema-telangiectasia. | Irrthum, A. et al. Mutations in the transcription factor gene SOX18 underlie recessive and dominant forms of hypotrichosis-lymphedema-telangiectasia. Am. J. Hum. Genet. 72, 1470–1478 (2003). |

| | | | | | |
|---|---|---|---|---|---|
| | **Vascular Endothelial Growth Factor A (VEGFA)** | **6p21.1** | **192240** | **A central regulator of physiological and pathological angiogenesis, with a key role in vascular integrity and maintenance.** | **Ferrara, N. Molecular and biological properties of vascular endothelial growth factor. Journal of Molecular Medicine 77, 527–543 (1999).** |
| **Vascular smooth muscle cell proliferation and migration** | **Family with Sequence Similarity 13, Member A (FAM13A)** | **4q22.1** | **613299** | **FAM13A regulates β-Catenin signalling pathways which promotes endothelial-mesenchymal transition in aortic valve endothelia.** | **Zhong, A., Mirzaei, Z. & Simmons, C. A. The Roles of Matrix Stiffness and ß-Catenin Signaling in Endothelial-to-Mesenchymal Transition of Aortic Valve Endothelial Cells. Cardiovasc. Eng. Technol. 9, 158–167 (2018).** |
| | **Kruppel-Like Factor 5 (KLF5)** | **13q22.1** | **602903** | **Plays a fundamental role in vascular remodelling via vSMC migration and proliferation.** | **Shindo, T. et al. Krüppel-like zinc-finger transcription factor KLF5/BTEB2 is a target for angiotensin II signaling and an essential regulator of cardiovascular remodeling. Nat. Med. 8, 856–863 (2002).** |
| | **Lim Domain-Containing Preferred Translocation Partner In Lipoma (LPP)** | **3q27-q28** | **600700** | **Induced in cultured vSMCs following vascular injury, involved in regulating cytoskeletal artchitecture and cell motility.** | **Majesky, M. W. Organizing motility: LIM domains, LPP, and smooth muscle migration. Circulation Research 98, 306–308 (2006).** |
| | **Sorbin and Sh3 Domains-Containing Protein 2 (SORBS2)** | **4q35.1** | **616349** | **Localises at apical junction in epithelial cells where it forms cell-cell contact. It is involved in endothelial cell migration and cellular senescence.** | **Anekal, P. V., Yong, J. & Manser, E. Arg kinase-binding protein 2 (ArgBP2) interaction with α-actinin and actin stress fibers inhibits cell migration. J. Biol. Chem. 290, 2112–2125 (2015).** |
| | **Tetraspanin 2 (TSPAN2)** | **1p13.2** | **613133** | **Selectively expressed in vSKC where it is associated with vSMC differentiation - induced in normal contractile phenotypes and suppresses proliferation and migration of vSMCs.** | **Todd, S. C., Doctor, V. S. & Levy, S. Sequences and expression of six new members of the tetraspanin/TM4SF family. Biochim. Biophys. Acta - Gene Struct. Expr. 1399, 101–104 (1998).** |

| | | | | | |
|---|---|---|---|---|---|
| | **Ubiquitin-Specific Peptidase 53 (USP53)** | **4q26** | **617431** | **Highly expressed in cardiac muscle, down-regulated during phenotyping modulation of vSMCs in atherosclerosis.** | **Matic, L. P. et al. Phenotypic modulation of smooth muscle cells in atherosclerosis is associated with downregulation of LMOD1, SYNPO2, PDLIM7, PLN, and SYNM. Arterioscler. Thromb. Vasc. Biol. 36, 1947–1961 (2016).** |
| **Extracellular matrix breakdown** | **Collagen, Type XXVII, Alpha-1 (COL27A1)** | **9q32** | **608461** | **Encodes a fibrillar colagen which supports extracellular matrices.** | **Pace, J. M., Corrado, M., Missero, C. & Byers, P. H. Identification, characterization and expression analysis of a new fibrillar collagen gene, COL27A1. Matrix Biol. 22, 3–14 (2003).** |
| | EGF-Containing Fibulin-Like Extracellular Matrix Protein 1 (EFEMP1) | 2p16.1 | 601548 | Highly-expressed in vein endothelia, where it encodes fibulin-3, which antagonises vascular development by inhibiting matrix metalloproteases and enhancing expression of their inhibitors. | Albig, A. R., Neil, J. R. & Schiemann, W. P. Fibulins 3 and 5 antagonize tumor angiogenesis in vivo. Cancer Res. 66, 2621–2629 (2006). |
| | Fibrillin 2 (FBN2) | 5q23.3 | 612570 | Preferentially expressed in elastic matrices where it egulates elastic fibre assembly. | Zhang, H. et al. Structure and expression of fibrillin-2, a novel microfibrillar component preferentially located in elastic matrices. J. Cell Biol. 124, 855–863 (1994). |
| | **Gnas Complex Locus (GNAS)** | **20q13.32** | **139320** | **Encodes a protein involved in the cAMP/Protein Kinase A pathway which has been suggested to inhibit fibroblast proliferation in skin and inhibits collagen synthesis in endothelial cells.** | **D'Angelo, G., Lee, H. & Weiner, R. I. cAMP-dependent protein kinase inhibits the mitogenic action of vascular endothelial growth factor and fibroblast growth factor in capillary endothelial cells by blocking Raf activation. J. Cell. Biochem. 67, 353–66 (1997).** |
| **Immune response** | **Af4/Fmr2 Family, Member 1 (AFF1)** | **4q21.3-q22.1** | **159557** | **Expressed in CD4+ and CD19+ peripheral blood lymphocytes and found to be associated with the autoimmune disease, systemic lupus erythematosus.** | **Okada, Y. et al. A Genome-Wide Association Study Identified AFF1 as a Susceptibility Locus for Systemic Lupus Eyrthematosus in Japanese. PLoS Genet. 8, e1002455 (2012).** |

| | | | | | |
|---|---|---|---|---|---|
| | **Dedicator Of Cytokinesis 8 (DOCK8)** | **9p24.3** | **611432** | **Implicated in the innate and adaptive immune system, with deletion resulting in the primary immunodeficiency, hyper-IgE symdrome.** | **Engelhardt, K. R. et al. Large deletions and point mutations involving the dedicator of cytokinesis 8 (DOCK8) in the autosomal-recessive form of hyper-IgE syndrome. J. Allergy Clin. Immunol. 124, (2009).** |
| | Early B-Cell Factor 1 (EBF1) | 5q33.3 | 164343 | Involved in regulating B-cell differentiation. | Gisler R, Jacobsen EW, Sigvardsson M. Cloning of human early B-cell factor and identification of target genes suggest a conserved role in B-cell development in man and mouse. Blood. 2000;96: 1457-1464. |
| | Nuclear Factor Of Activated T Cells, Cytoplasmic, Calcineurin-Dependent 2 (NFATC2) | 20q13.2 | 600490 | Involved in the regulation of activation of naïve T-Cells, where it is a negative regulator of cell cycle progression in normal cells. | Caetano MS, Vieira-de-Abreu A, Teixeira LK, Werneck MB, Barcinski MA, Viola JP. NFATC2 transcription factor regulates cell cycle progression during lymphocyte activation: evidence of its involvement in the control of cyclin gene expression. FASEB J. 2002;16(14):1940–1942. doi:10.1096/fj.02-0282fje |
| | Protein Phosphatase 3, Regulatory Subunit B, Alpha (PPP3R1) | 2p14 | 601302 | Encodes a subunit of calcineurin, which is involved in the activation of naive T-cells. | Wu W, Chen Q, Geng F, Tong L, Yang R, Yang J, et al. Calcineurin B stimulates cytokine production through a CD14-independent Toll-like receptor 4 pathway. Immunol Cell Biol. 2016;94: 285–292. doi:10.1038/icb.2015.91 |
| **Apoptosis** | **Ceramide Synthase 5 (CERS5)** | **12q13.12** | **615335** | **CERS5 is involved in apoptosis and autophagy pathways, particularly in response to cellular stress and in several cancers.** | **Hannun, Y. A. Functions of ceramide in coordinating cellular responses to stress. Science (80-. ). 274, 1855–1859 (1996).** |
| | **Cyclin-Dependent Kinase-Like 1 (CDKL1)** | **14q21.3** | **603441** | **Interacts with cyclin to regulate the cell cycle and cell death.** | **Wang, Y. & Huang, Q. Downregulation of CDKL1 promotes gastric cell apoptosis through inhibiting cell growth and colony formation. Int J Clin Exp Med 10, (2017).** |
| | Cyclin-Dependent Kinase 10 (CDK10) | 16q24.3 | 603464 | Essential for cell proliferation (positive control), acting during the G2-M phase of the cell cycle. | Li S, MacLachlan TK, De Luca A, Claudio PP, Condorelli G, Giordano A. The cdc-2-related kinase, PISSLRE, is essential for cell growth and acts in G2 phase of the cell cycle. Cancer Res. 1995; 55:3992–95. |

#### 2. Supplementary Figures

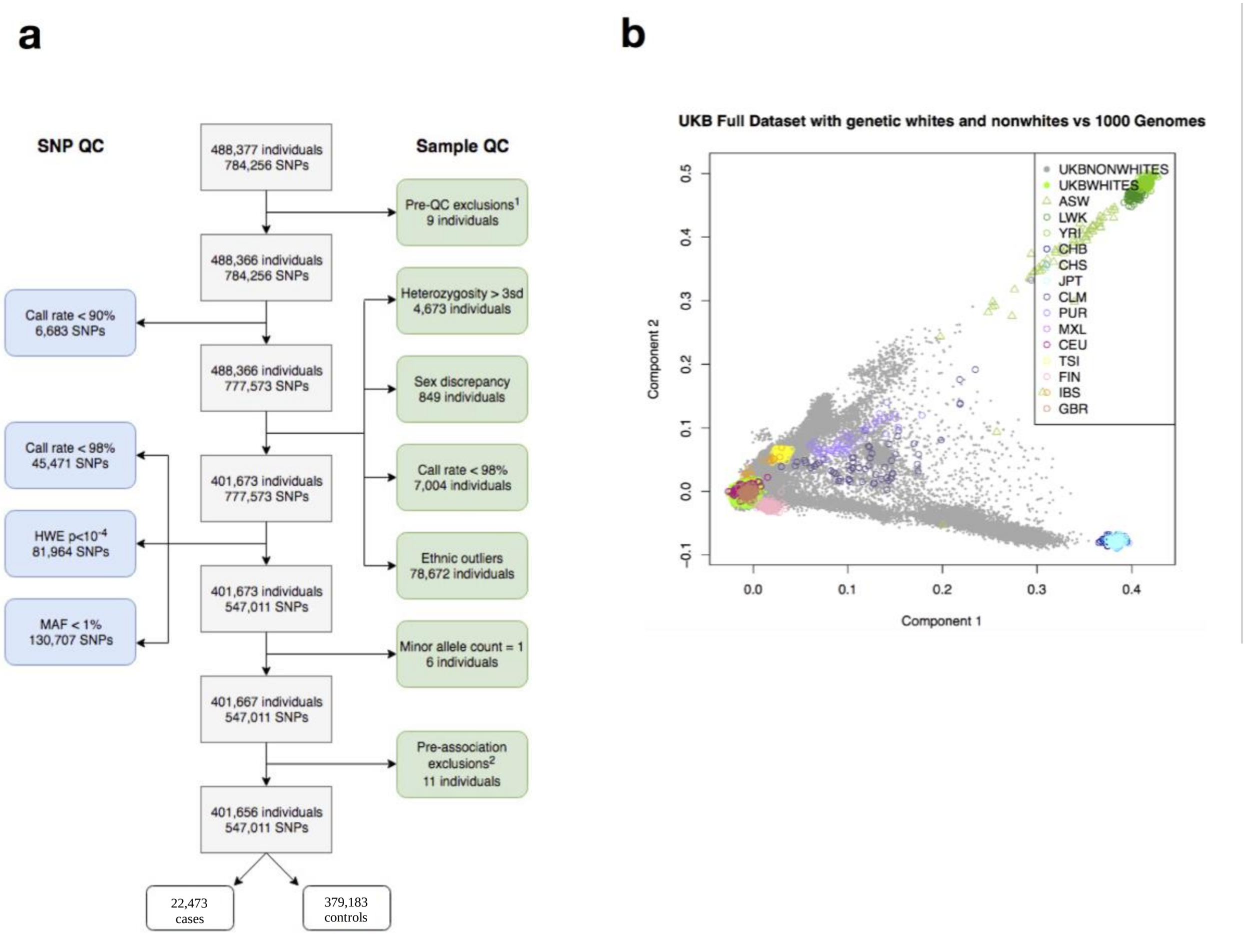

**Supplementary Figure 1. Overview of Quality Control (QC). a**, Flowchart summarising QC protocol. Excluded SNPs are in blue panels on the left and excluded individuals are in green panels on the right. [1]Pre-QC exclusions: 3 individuals with invalid IDs and sex, and 8 individuals who have withdrawn from UK Biobank were excluded prior to QC. [2]Pre-association exclusions: 11 individuals who were not present in UK Biobank's sample file accompanying the BGEN files were excluded prior to association. **b**, Principal Component Analysis (PCA) for demonstration of ethnicity of UK Biobank individuals. The UK Biobank cohort was merged with publicly available data from the 1000 Genomes Project and PCA was performed using flashpca. Individuals identified by UK Biobank as having white British ancestry are coloured in lime green, and the remaining UK Biobank individuals are in grey. In this graph of principal component 1 vs principal component 2, a near-perfect overlap can be seen between the UK Biobank "white British" individuals and both GBR (British in England and Scotland - light brown) and CEU (Utah residents with Northern and Western European ancestry - magenta) individuals from the 1000 Genomes Project.

**Supplementary Figure 2. Regional Locus Zoom plots of all varicose veins associated loci.** LocusZoom plots of the 49 independent genome-wide significant SNPs at the 46 replicated varicose veins associated susceptibility loci. Plots are ordered by chromosome number and genomic position. SNP position is shown on the x-axis, and strength of association on the y-axis (-log10 P-value). The linkage disequilibrium (LD) relationship between the lead SNP and the surrounding SNPs is indicated by the r2 legend. In the lower panel of each sub-figure, genes within 500kb of the index SNP are shown. The position on each chromosome is shown in relation to Human Genome build hg19.

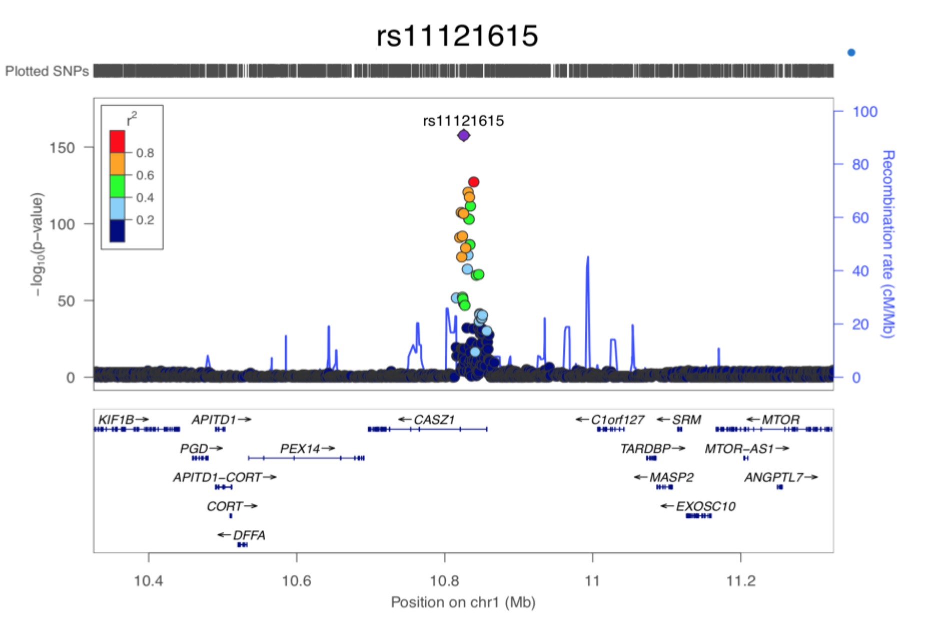


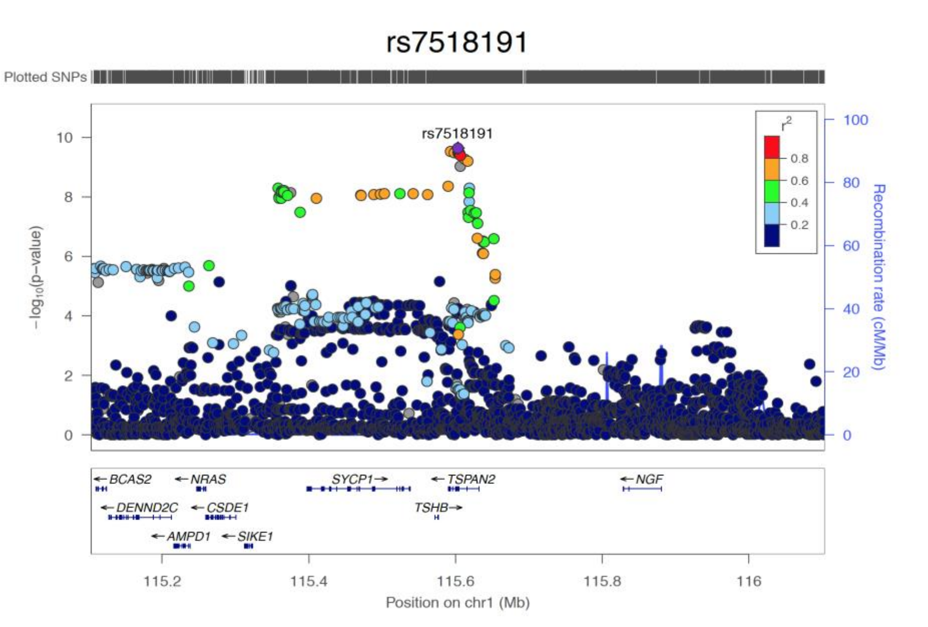

rs17712208

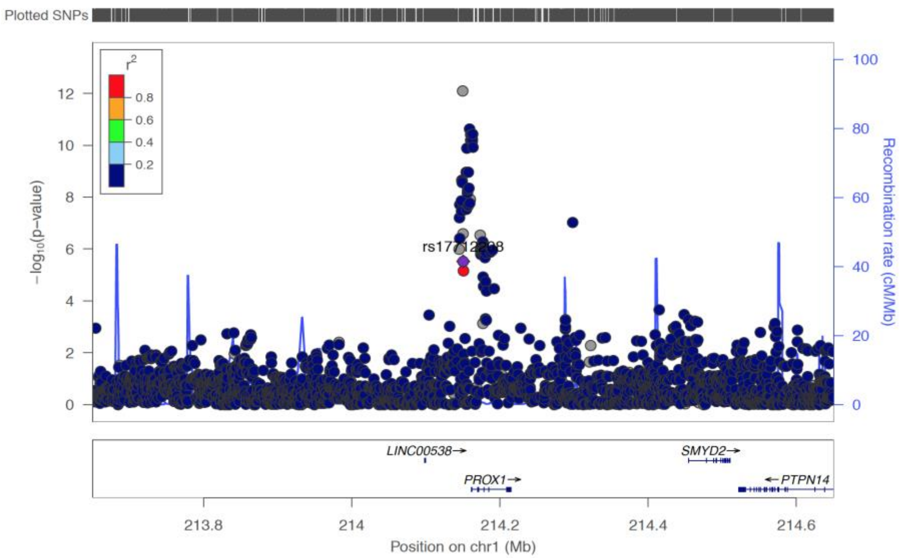


rs340875

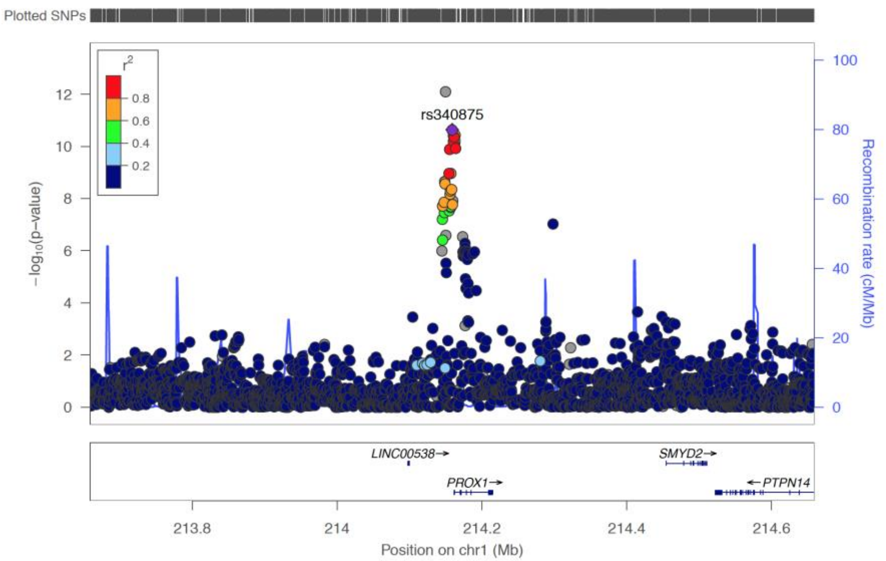


rs2820464

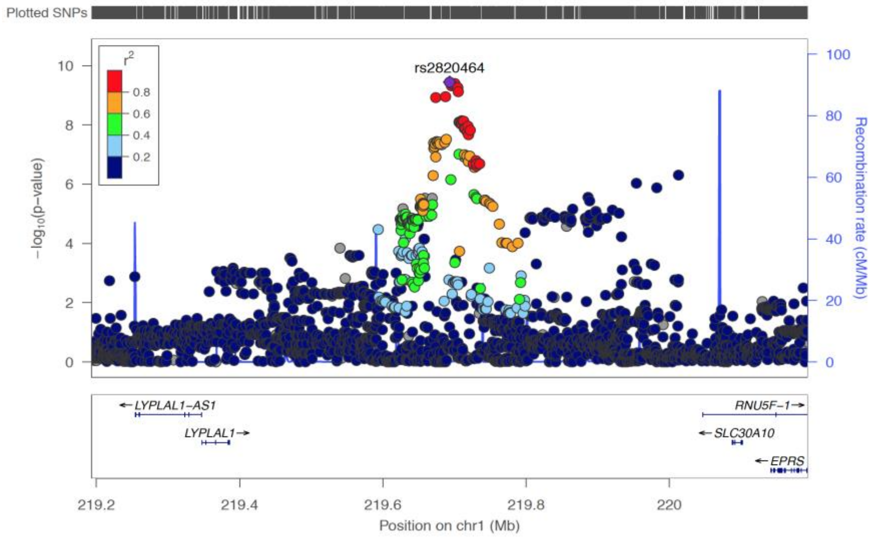


rs9967884

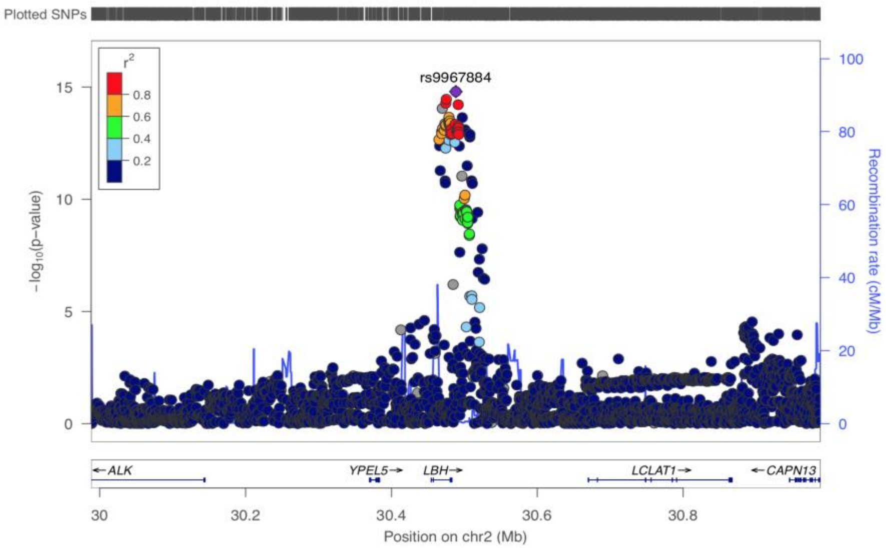

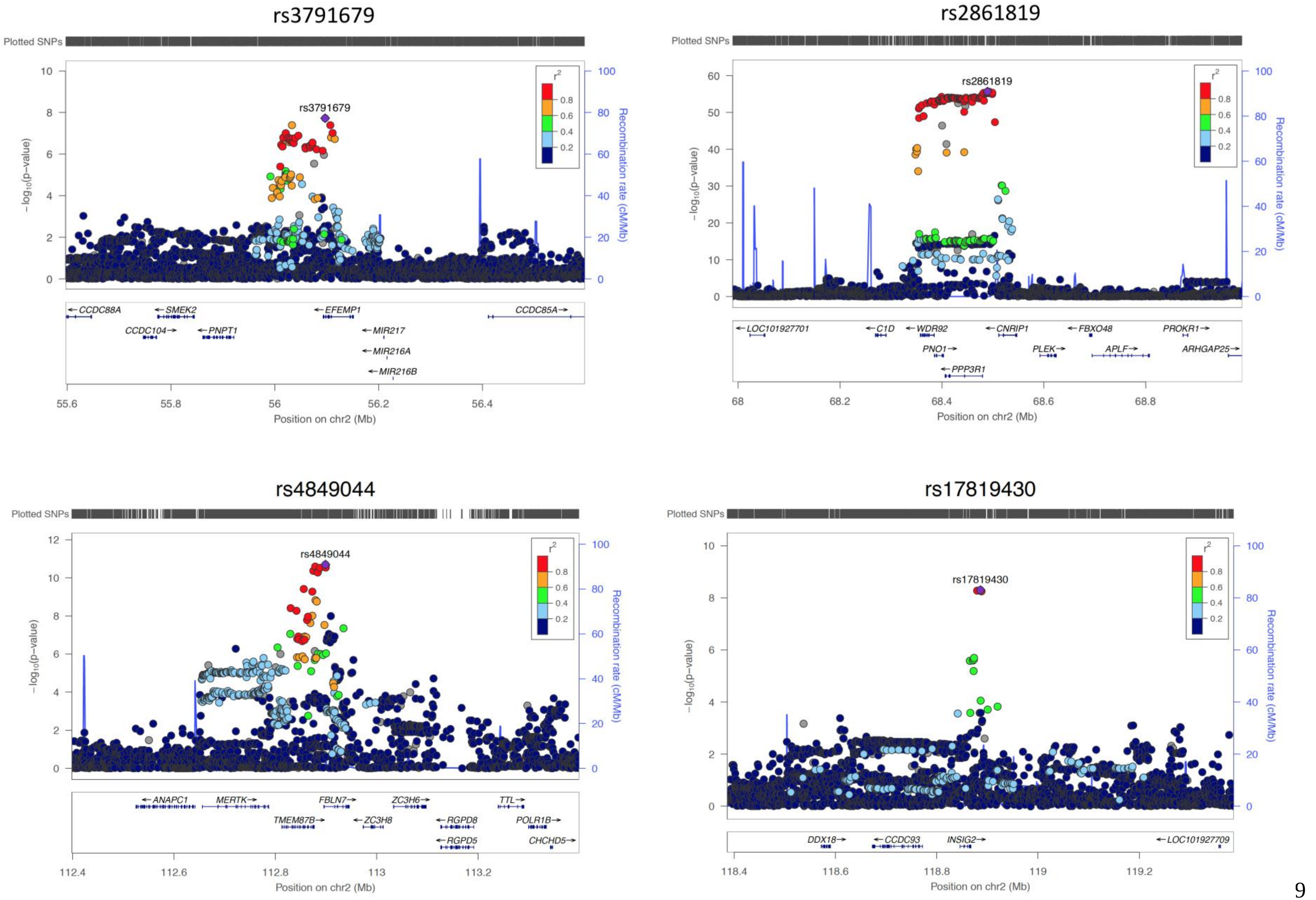

rs3791679
Plotted SNPs
rs3791679
r²
0.8
0.6
0.4
0.2
−log10(p−value)
Recombination rate (cM/Mb)
CCDC88A
SMEK2
EFEMP1
CCDC85A
CCDC104
PNPT1
MIR217
MIR216A
MIR216B
55.6
55.8
56
56.2
56.4
Position on chr2 (Mb)
rs2861819
Plotted SNPs
rs2861819
LOC101927701
C1D
WDR92
CNRIP1
FBXO48
PROKR1
PNO1
PLEK
APLF
ARHGAP25
PPP3R1
68
68.2
68.4
68.6
68.8
Position on chr2 (Mb)
rs4849044
Plotted SNPs
rs4849044
ANAPC1
MERTK
FBLN7
ZC3H6
TTL
TMEM87B
ZC3H8
RGPD8
POLR1B
RGPD5
CHCHD5
112.4
112.6
112.8
113
113.2
Position on chr2 (Mb)
rs17819430
Plotted SNPs
rs17819430
DDX18
CCDC93
INSIG2
LOC101927709
118.4
118.6
118.8
119
119.2
Position on chr2 (Mb)

## rs55889669

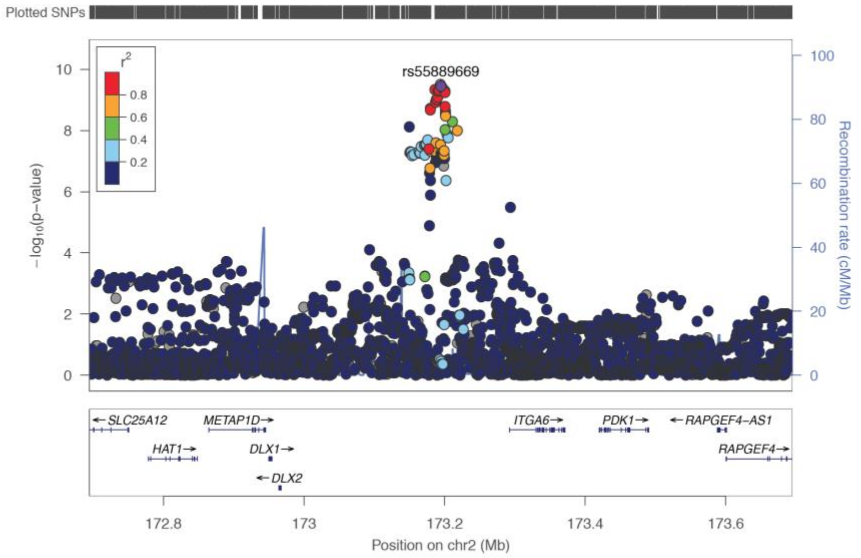


## rs844176

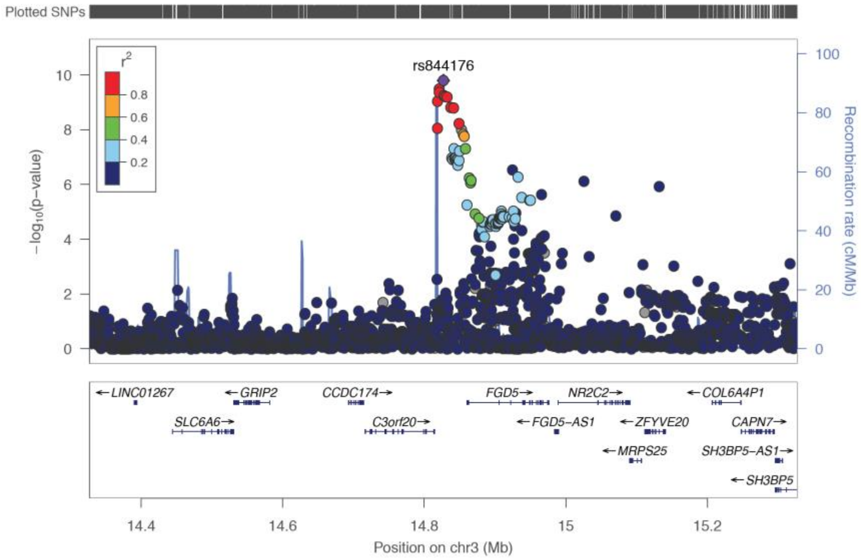


## rs2713575

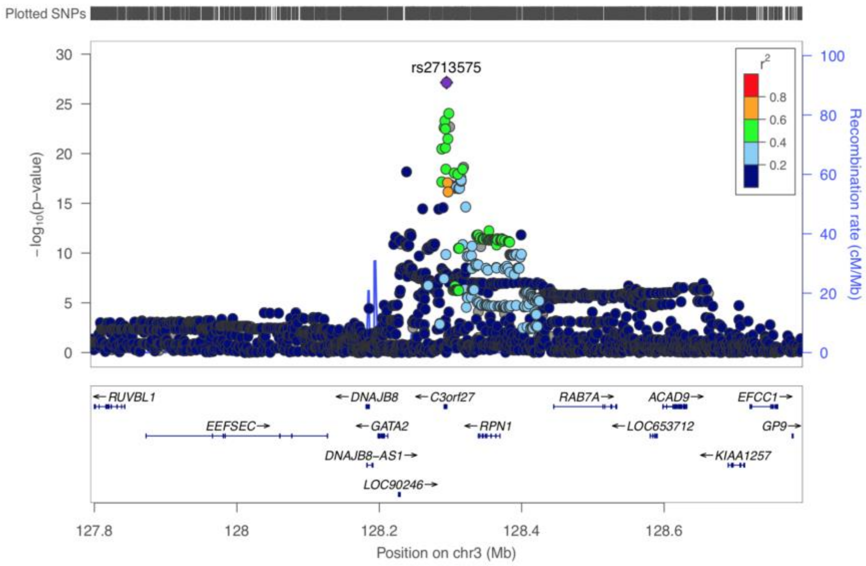


## rs9877579

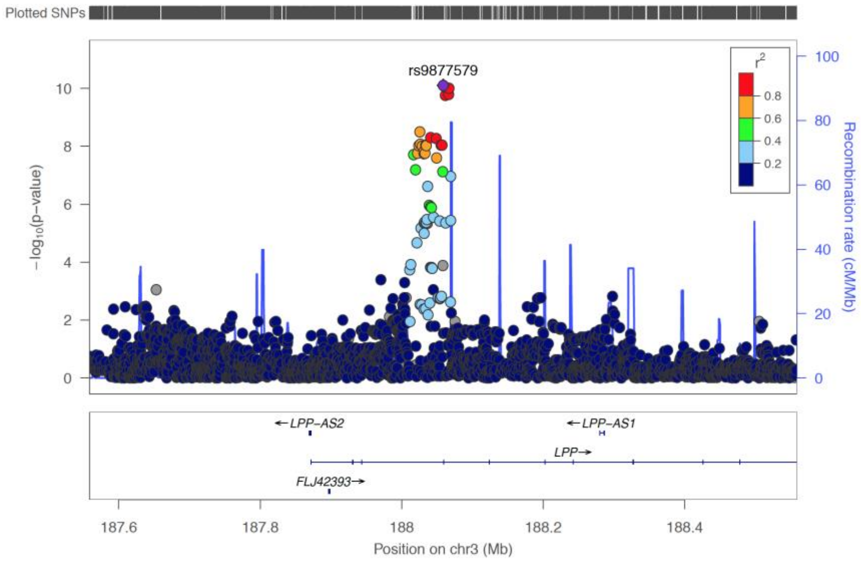

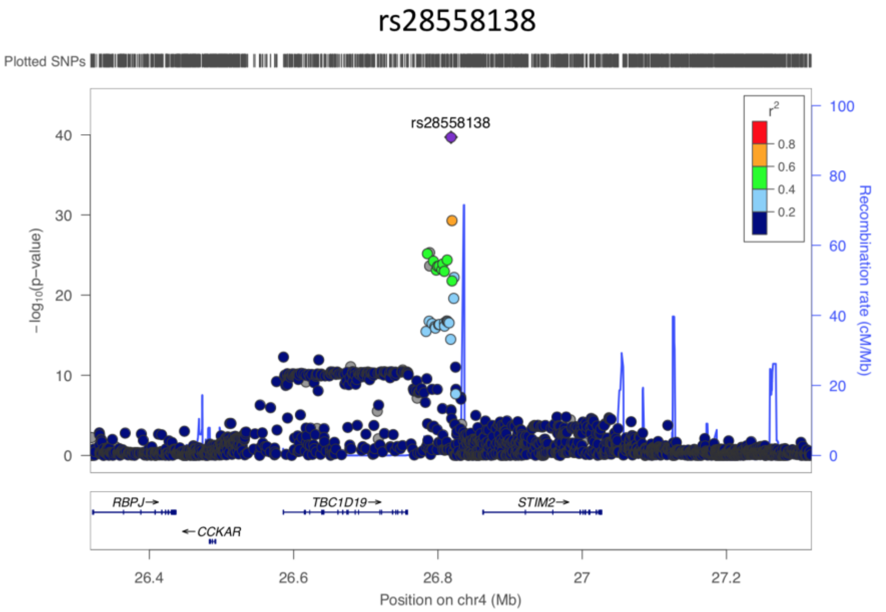
rs28558138
Plotted SNPs
rs28558138
r²
0.8
0.6
0.4
0.2
−log10(p−value)
Recombination rate (cM/Mb)
RBPJ→
←CCKAR
TBC1D19→
STIM2→
26.4
26.6
26.8
27
27.2
Position on chr4 (Mb)

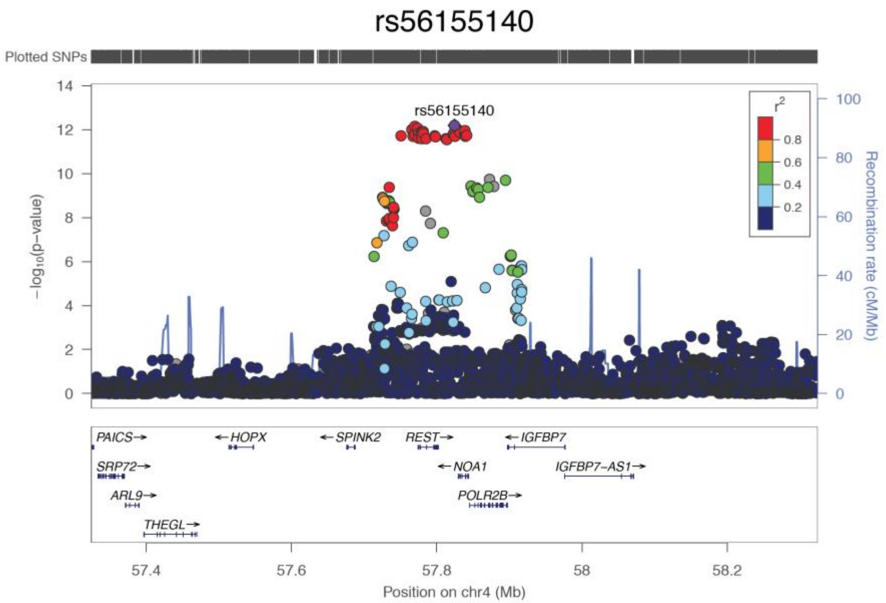
rs56155140
Plotted SNPs
rs56155140
r²
0.8
0.6
0.4
0.2
−log10(p−value)
Recombination rate (cM/Mb)
PAICS→
←HOPX
←SPINK2
REST→
←IGFBP7
SRP72→
←NOA1
IGFBP7−AS1→
ARL9→
POLR2B→
THEGL→
57.4
57.6
57.8
58
58.2
Position on chr4 (Mb)

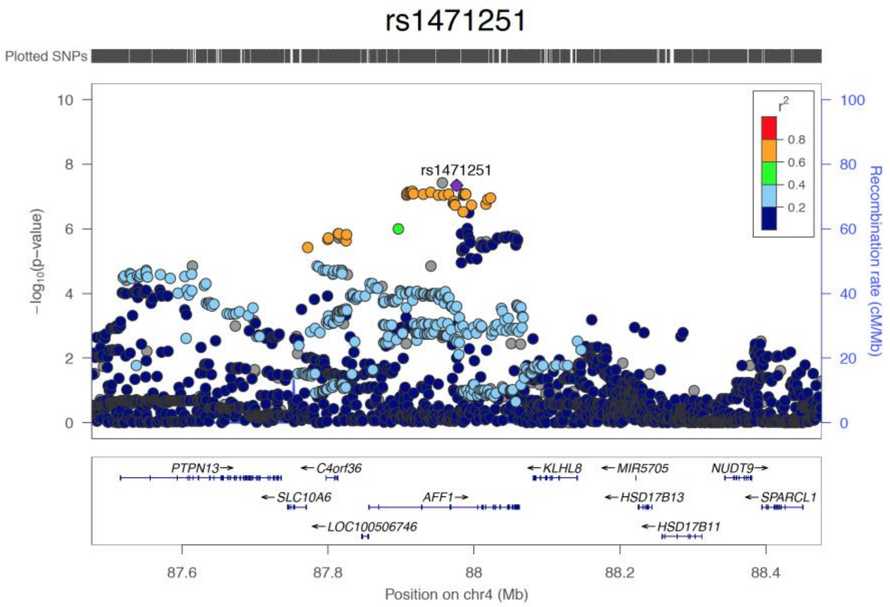
rs1471251
Plotted SNPs
rs1471251
r²
0.8
0.6
0.4
0.2
−log10(p−value)
Recombination rate (cM/Mb)
PTPN13→
←C4orf36
←KLHL8
←MIR5705
NUDT9→
←SLC10A6
AFF1→
←HSD17B13
←SPARCL1
←LOC100506746
←HSD17B11
87.6
87.8
88
88.2
88.4
Position on chr4 (Mb)

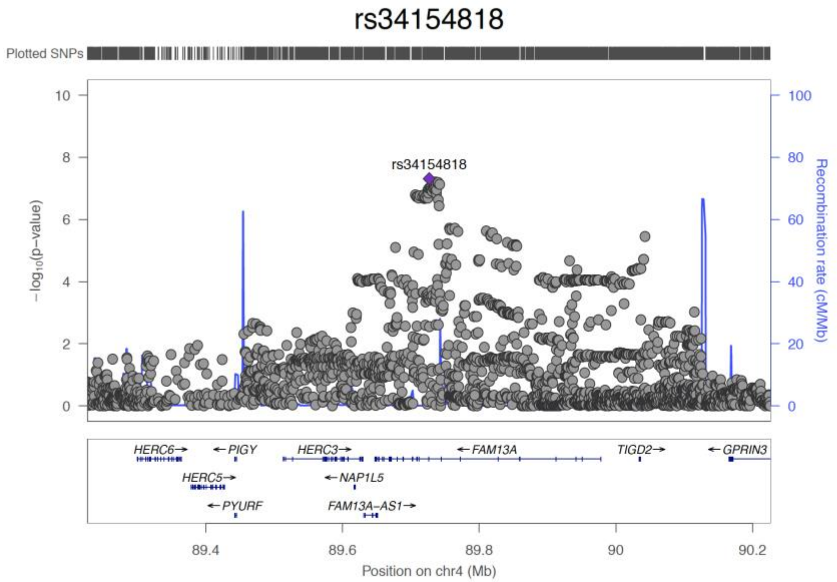
rs34154818
Plotted SNPs
rs34154818
−log10(p−value)
Recombination rate (cM/Mb)
HERC6→
←PIGY
HERC3→
←FAM13A
TIGD2→
←GPRIN3
HERC5→
←NAP1L5
←PYURF
FAM13A−AS1→
89.4
89.6
89.8
90
90.2
Position on chr4 (Mb)

rs10007409

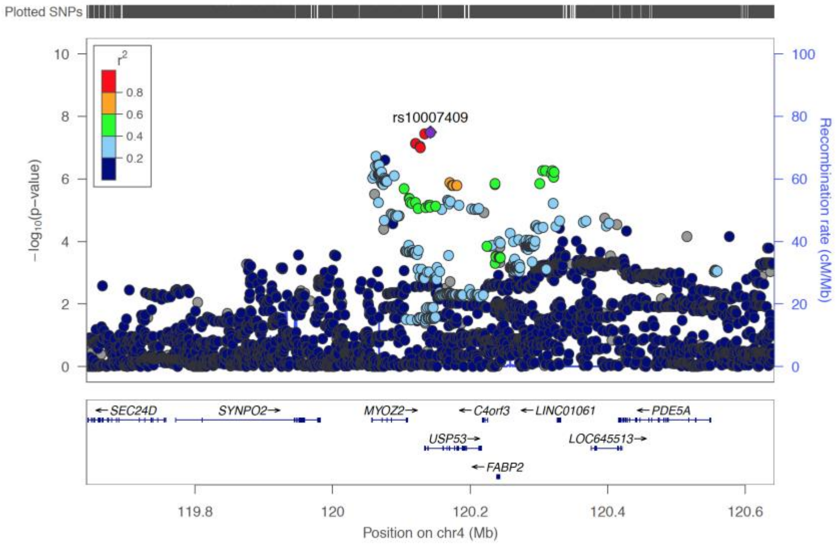


rs11728719

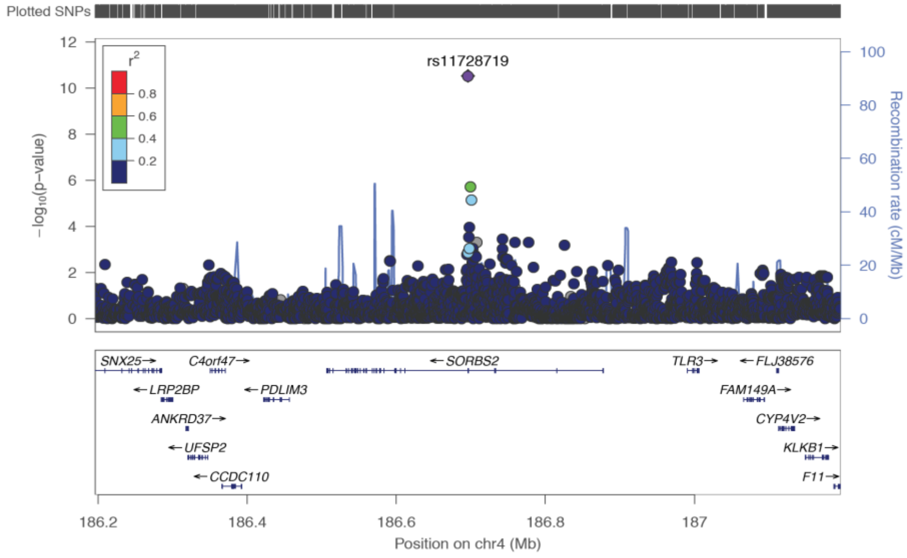


rs57253948

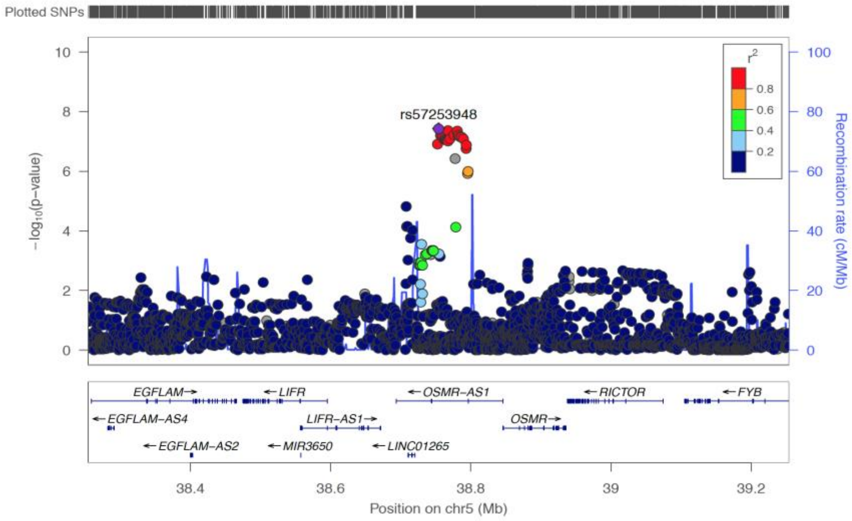


rs3749748

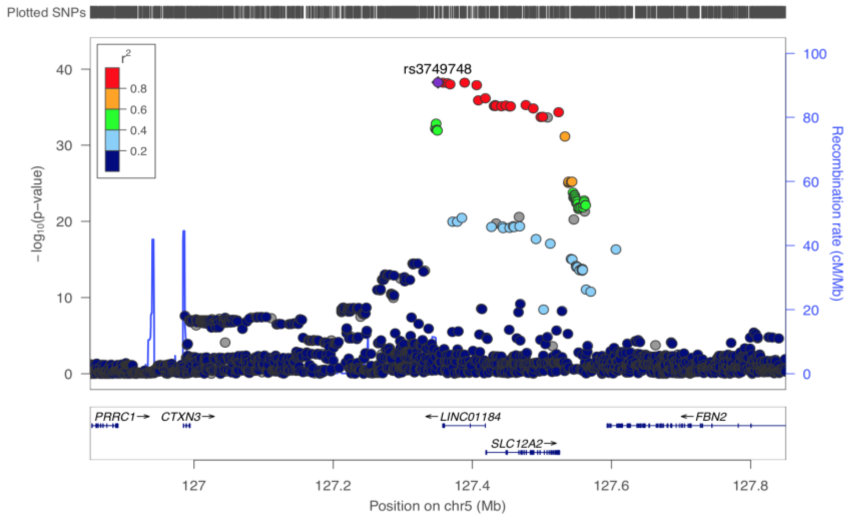

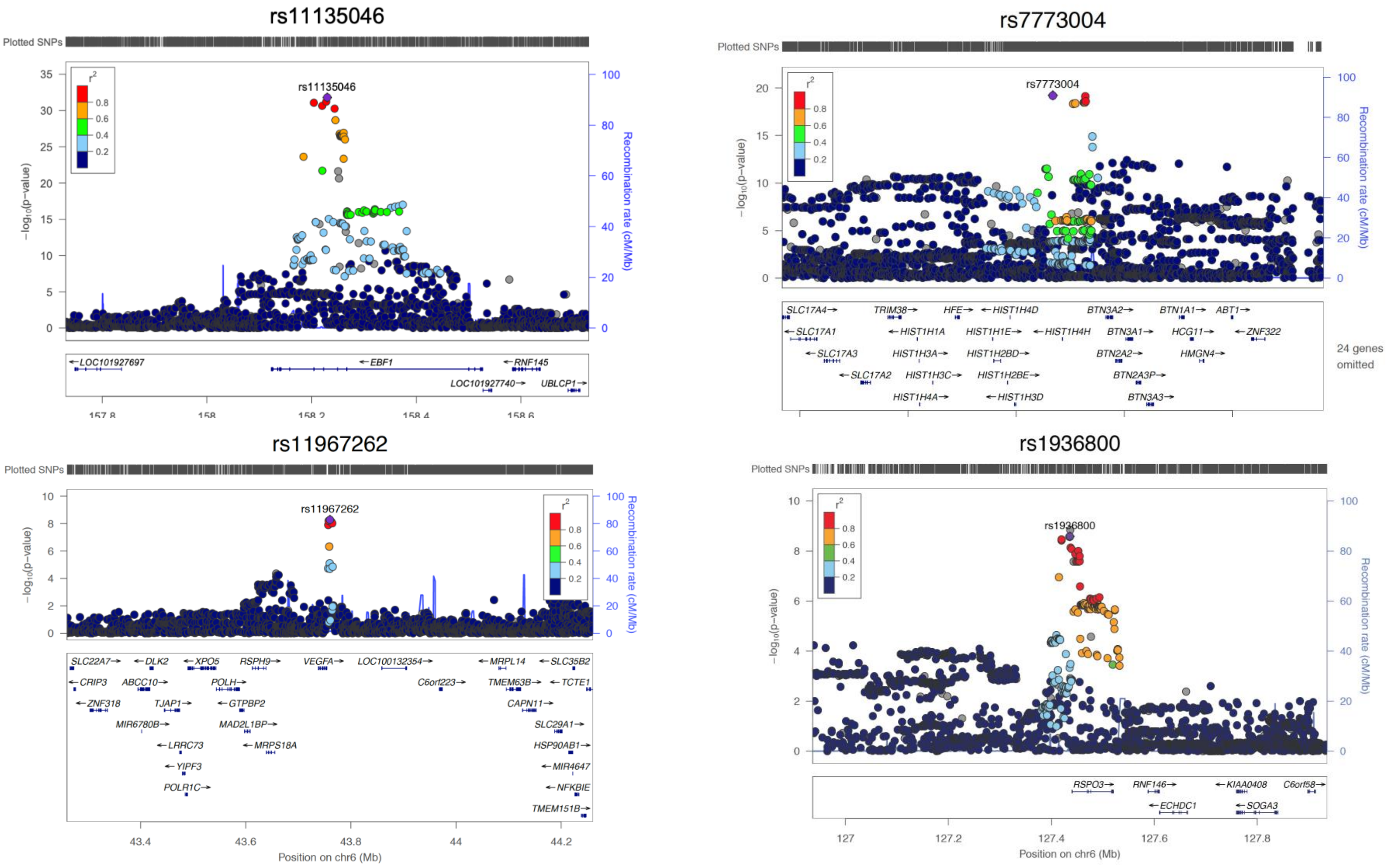
rs11135046
Plotted SNPs
−log10(p−value)
Recombination rate (cM/Mb)
r2
0.8
0.6
0.4
0.2
rs11135046
LOC101927697
EBF1
RNF145
LOC101927740
UBLCP1
157.8
158
158.2
158.4
158.6
rs7773004
Plotted SNPs
−log10(p−value)
Recombination rate (cM/Mb)
rs7773004
SLC17A4
TRIM38
HFE
HIST1H4D
BTN3A2
BTN1A1
ABT1
SLC17A1
HIST1H1A
HIST1H1E
HIST1H4H
BTN3A1
HCG11
ZNF322
SLC17A3
HIST1H3A
HIST1H2BD
BTN2A2
HMGN4
SLC17A2
HIST1H3C
HIST1H2BE
BTN2A3P
HIST1H4A
HIST1H3D
BTN3A3
24 genes omitted
rs11967262
Plotted SNPs
−log10(p−value)
Recombination rate (cM/Mb)
rs11967262
SLC22A7
DLK2
XPO5
RSPH9
VEGFA
LOC100132354
MRPL14
SLC35B2
CRIP3
ABCC10
POLH
C6orf223
TMEM63B
TCTE1
ZNF318
TJAP1
GTPBP2
CAPN11
MIR6780B
MAD2L1BP
SLC29A1
LRRC73
MRPS18A
HSP90AB1
YIPF3
MIR4647
POLR1C
NFKBIE
TMEM151B
43.4
43.6
43.8
44
44.2
Position on chr6 (Mb)
rs1936800
Plotted SNPs
−log10(p−value)
Recombination rate (cM/Mb)
rs1936800
RSPO3
RNF146
KIAA0408
C6orf58
ECHDC1
SOGA3
127
127.2
127.4
127.6
127.8
Position on chr6 (Mb)

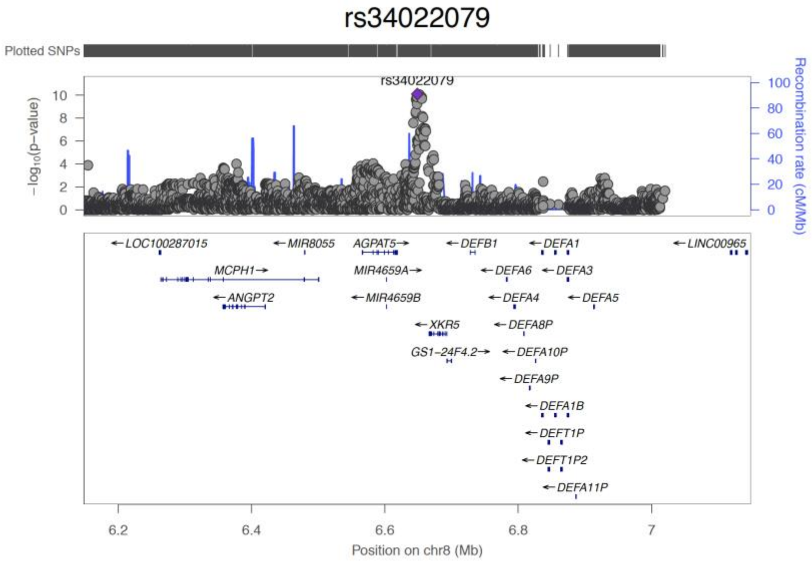
rs34022079
Plotted SNPs
rs34022079
−log10(p−value)
Recombination rate (cM/Mb)
LOC100287015
MIR8055
AGPAT5
DEFB1
DEFA1
LINC00965
MCPH1
MIR4659A
DEFA6
DEFA3
ANGPT2
MIR4659B
DEFA4
DEFA5
XKR5
DEFA8P
GS1−24F4.2
DEFA10P
DEFA9P
DEFA1B
DEFT1P
DEFT1P2
DEFA11P
Position on chr8 (Mb)

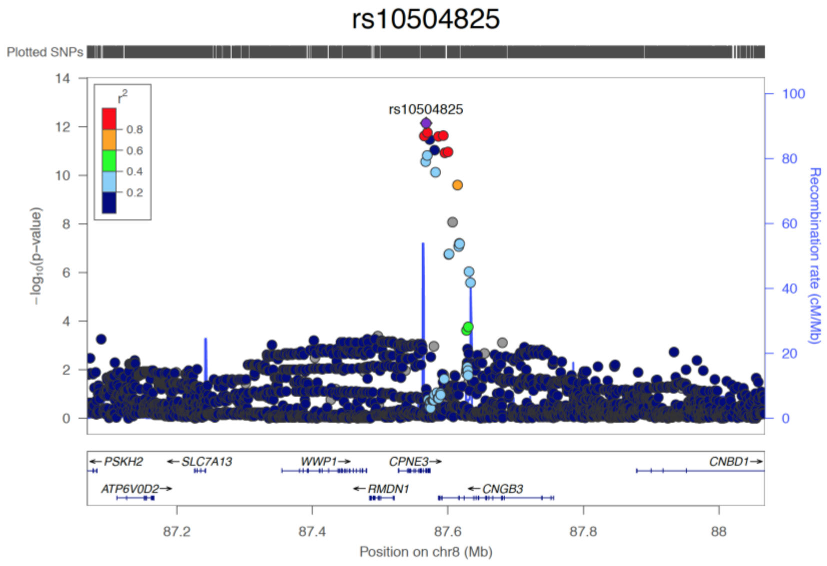
rs10504825
Plotted SNPs
rs10504825
−log10(p−value)
Recombination rate (cM/Mb)
PSKH2
SLC7A13
WWP1
CPNE3
CNBD1
ATP6V0D2
RMDN1
CNGB3
Position on chr8 (Mb)

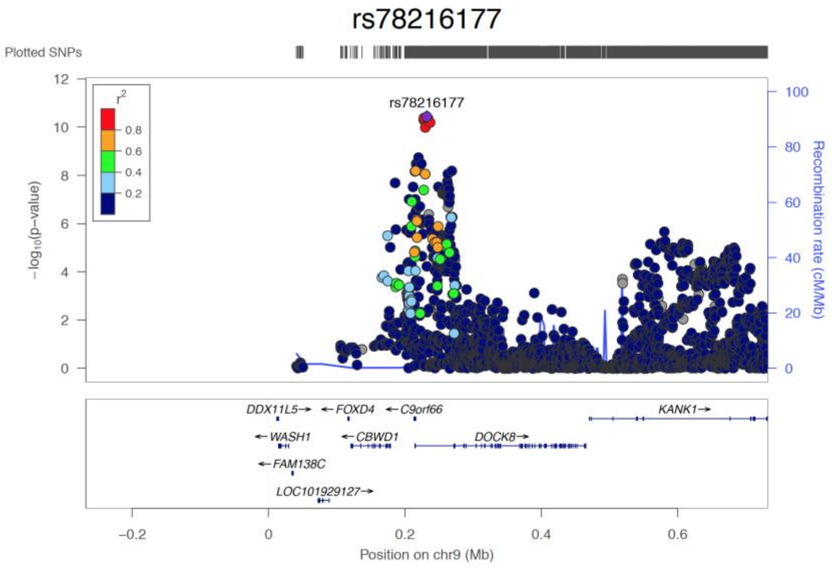
rs78216177
Plotted SNPs
rs78216177
−log10(p−value)
Recombination rate (cM/Mb)
DDX11L5
FOXD4
C9orf66
KANK1
WASH1
CBWD1
DOCK8
FAM138C
LOC101929127
Position on chr9 (Mb)

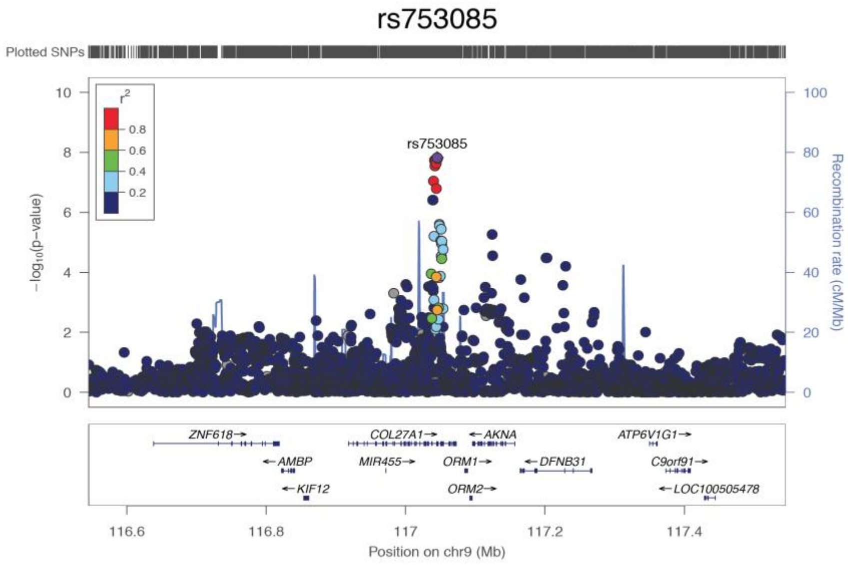
rs753085
Plotted SNPs
rs753085
−log10(p−value)
Recombination rate (cM/Mb)
ZNF618
COL27A1
AKNA
ATP6V1G1
AMBP
MIR455
ORM1
DFNB31
C9orf91
KIF12
ORM2
LOC100505478
Position on chr9 (Mb)

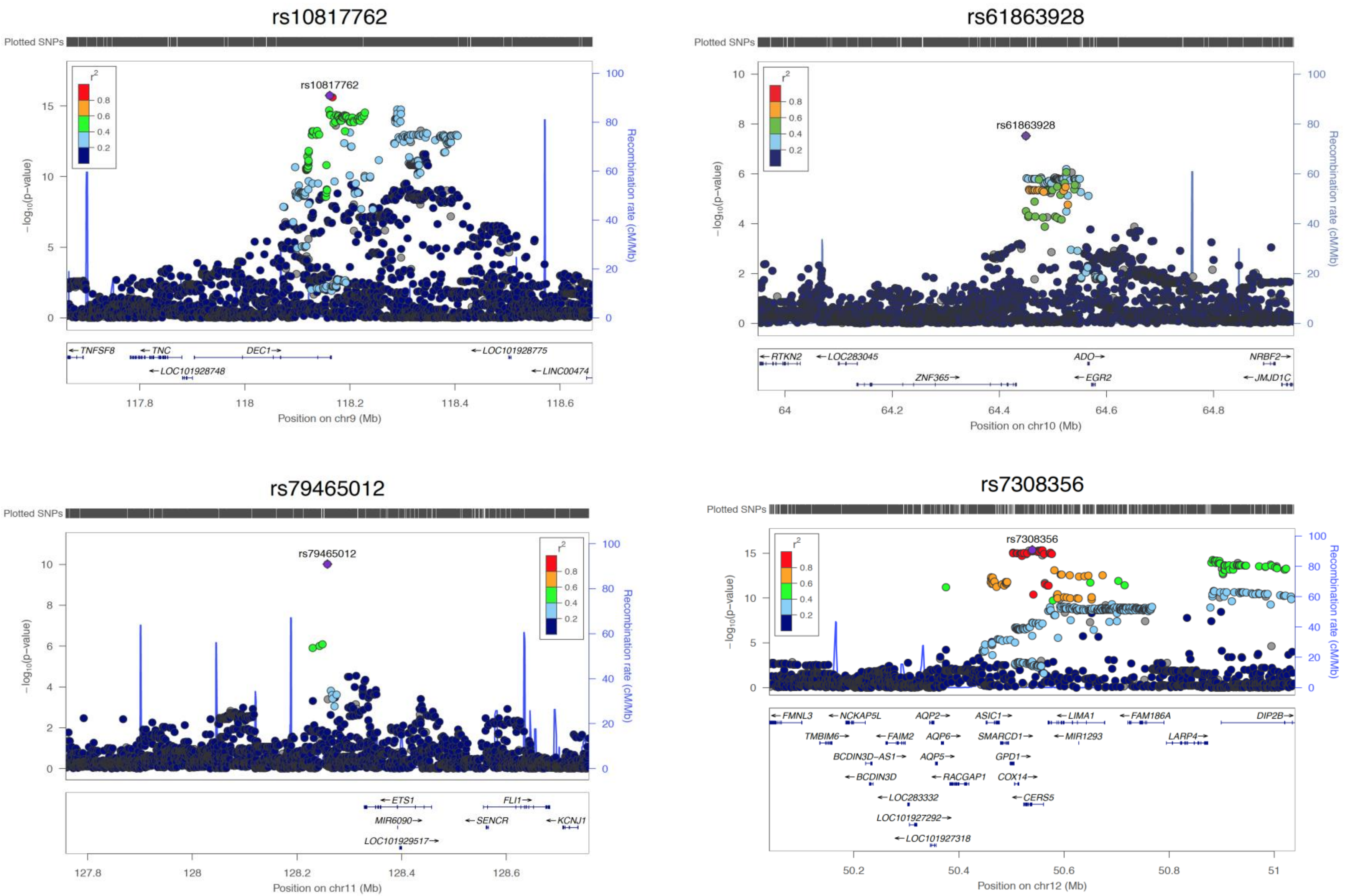
rs10817762
Plotted SNPs
r2
0.8
0.6
0.4
0.2
-log10(p-value)
Recombination rate (cM/Mb)
TNFSF8
TNC
DEC1
LOC101928775
LOC101928748
LINC00474
Position on chr9 (Mb)
rs61863928
RTKN2
LOC283045
ADO
NRBF2
ZNF365
EGR2
JMJD1C
Position on chr10 (Mb)
rs79465012
ETS1
FLI1
MIR6090
SENCR
KCNJ1
LOC101929517
Position on chr11 (Mb)
rs7308356
FMNL3
NCKAP5L
AQP2
ASIC1
LIMA1
FAM186A
DIP2B
TMBIM6
FAIM2
AQP6
SMARCD1
MIR1293
LARP4
BCDIN3D-AS1
AQP5
GPD1
BCDIN3D
RACGAP1
COX14
LOC283332
CERS5
LOC101927292
LOC101927318
Position on chr12 (Mb)

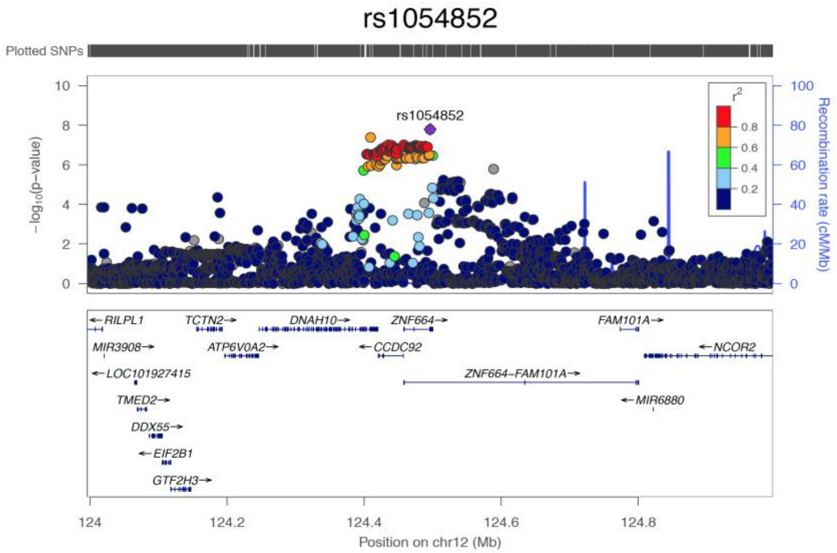
rs1054852
Plotted SNPs
rs1054852
−log10(p−value)
Recombination rate (cM/Mb)
r2
RILPL1
TCTN2
DNAH10
ZNF664
FAM101A
MIR3908
ATP6V0A2
CCDC92
NCOR2
LOC101927415
ZNF664−FAM101A
TMED2
MIR6880
DDX55
EIF2B1
GTF2H3
Position on chr12 (Mb)

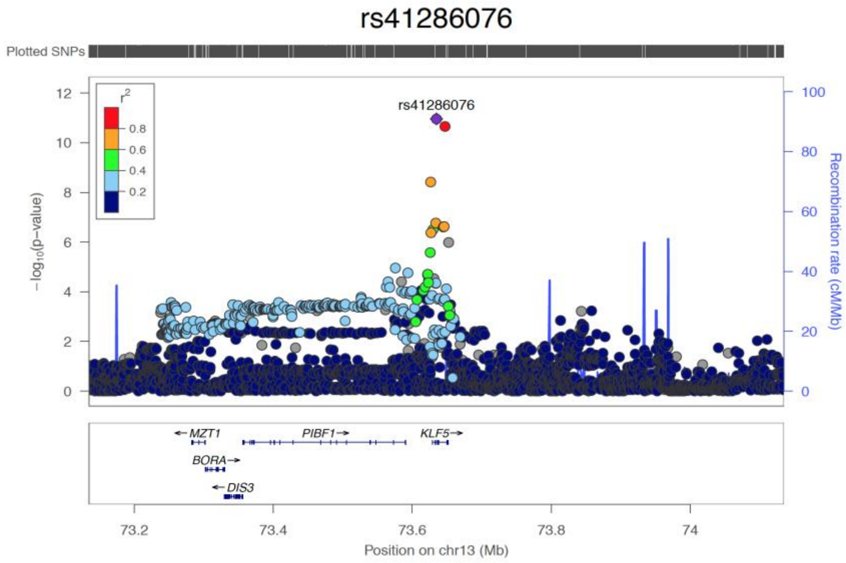
rs41286076
Plotted SNPs
rs41286076
−log10(p−value)
Recombination rate (cM/Mb)
r2
MZT1
PIBF1
KLF5
BORA
DIS3
Position on chr13 (Mb)

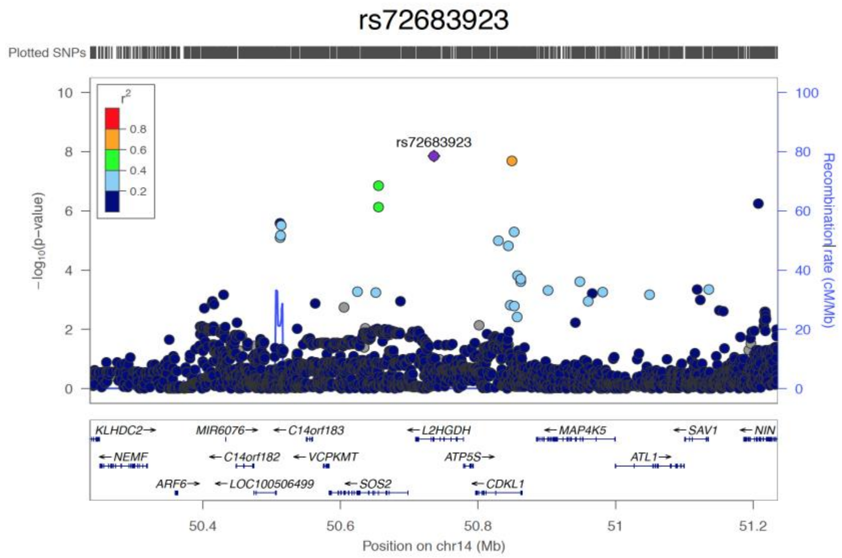
rs72683923
Plotted SNPs
rs72683923
−log10(p−value)
Recombination rate (cM/Mb)
r2
KLHDC2
MIR6076
C14orf183
L2HGDH
MAP4K5
SAV1
NIN
NEMF
C14orf182
VCPKMT
ATP5S
ATL1
ARF6
LOC100506499
SOS2
CDKL1
Position on chr14 (Mb)

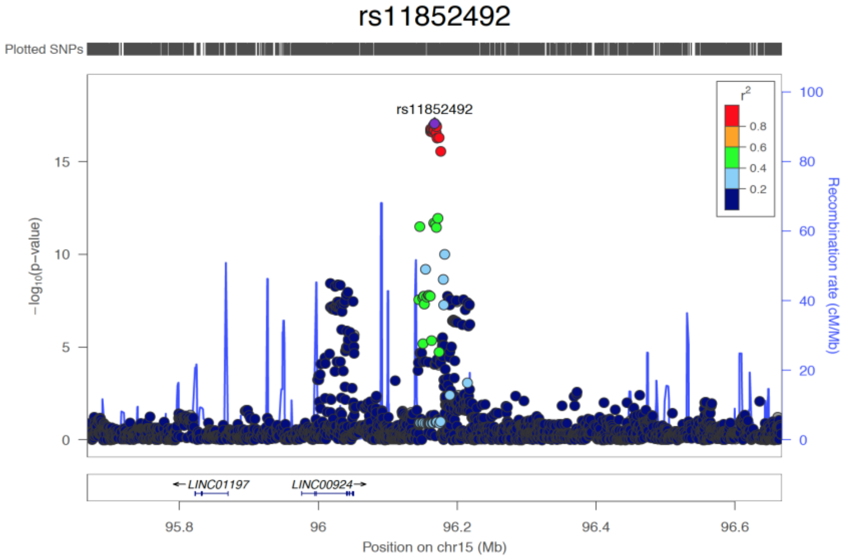
rs11852492
Plotted SNPs
rs11852492
−log10(p−value)
Recombination rate (cM/Mb)
r2
LINC01197
LINC00924
Position on chr15 (Mb)

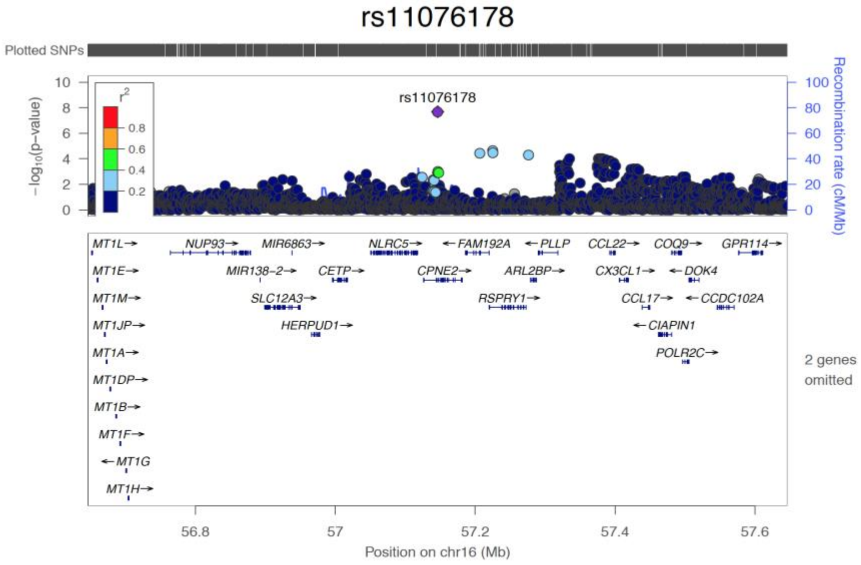
rs11076178
Plotted SNPs
−log10(p−value)
Recombination rate (cM/Mb)
r2
rs11076178
MT1L→ NUP93→ MIR6863→ NLRC5→ ←FAM192A ←PLLP CCL22→ COQ9→ GPR114→
MT1E→ MIR138−2→ CETP→ CPNE2→ ARL2BP→ CX3CL1→ ←DOK4
MT1M→ SLC12A3→ RSPRY1→ CCL17→ ←CCDC102A
MT1JP→ HERPUD1→ ←CIAPIN1
MT1A→ POLR2C→
MT1DP→
MT1B→
MT1F→
←MT1G
MT1H→
2 genes omitted
Position on chr16 (Mb)

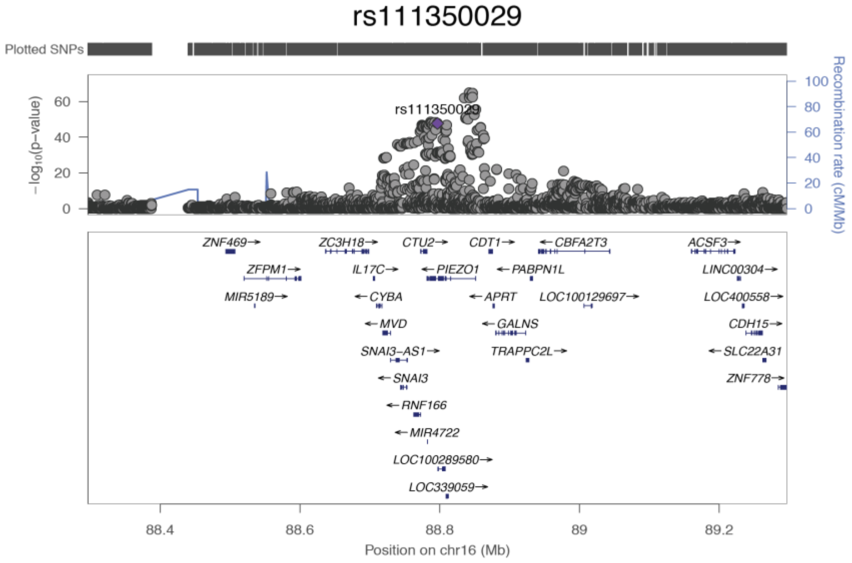
rs111350029
Plotted SNPs
−log10(p−value)
Recombination rate (cM/Mb)
rs111350029
ZNF469→ ZC3H18→ CTU2→ CDT1→ ←CBFA2T3 ACSF3→
ZFPM1→ IL17C→ ←PIEZO1 ←PABPN1L LINC00304→
MIR5189→ ←CYBA ←APRT LOC100129697→ LOC400558→
←MVD ←GALNS CDH15→
SNAI3−AS1→ TRAPPC2L→ ←SLC22A31
←SNAI3 ZNF778→
←RNF166
←MIR4722
LOC100289580→
LOC339059→
Position on chr16 (Mb)

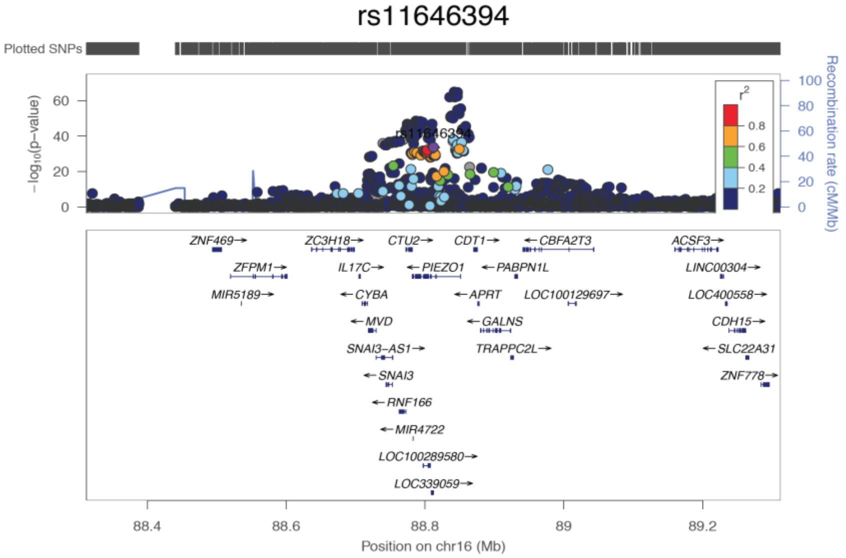
rs11646394
Plotted SNPs
−log10(p−value)
Recombination rate (cM/Mb)
r2
rs11646394
ZNF469→ ZC3H18→ CTU2→ CDT1→ ←CBFA2T3 ACSF3→
ZFPM1→ IL17C→ ←PIEZO1 ←PABPN1L LINC00304→
MIR5189→ ←CYBA ←APRT LOC100129697→ LOC400558→
←MVD ←GALNS CDH15→
SNAI3−AS1→ TRAPPC2L→ ←SLC22A31
←SNAI3 ZNF778→
←RNF166
←MIR4722
LOC100289580→
LOC339059→
Position on chr16 (Mb)

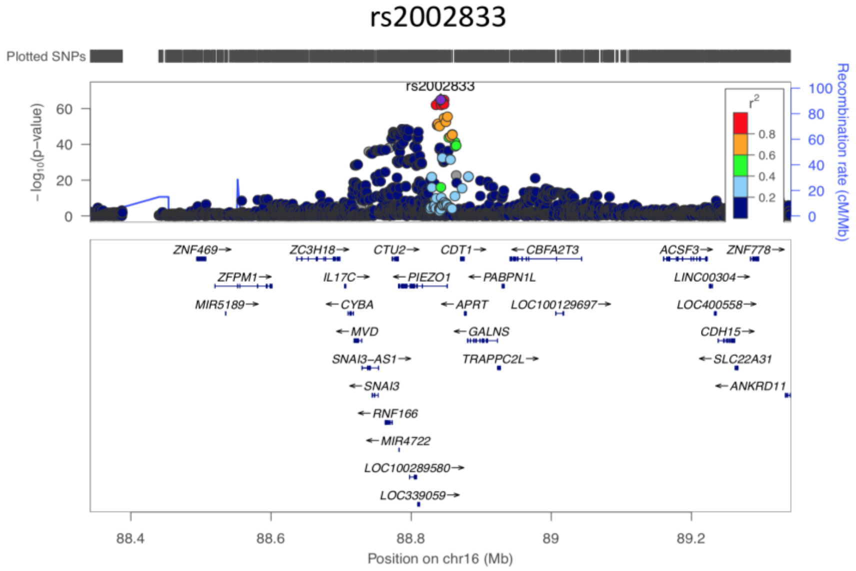
rs2002833
Plotted SNPs
−log10(p−value)
Recombination rate (cM/Mb)
r2
rs2002833
ZNF469→ ZC3H18→ CTU2→ CDT1→ ←CBFA2T3 ACSF3→ ZNF778→
ZFPM1→ IL17C→ ←PIEZO1 ←PABPN1L LINC00304→
MIR5189→ ←CYBA ←APRT LOC100129697→ LOC400558→
←MVD ←GALNS CDH15→
SNAI3−AS1→ TRAPPC2L→ ←SLC22A31
←SNAI3 ←ANKRD11
←RNF166
←MIR4722
LOC100289580→
LOC339059→
Position on chr16 (Mb)

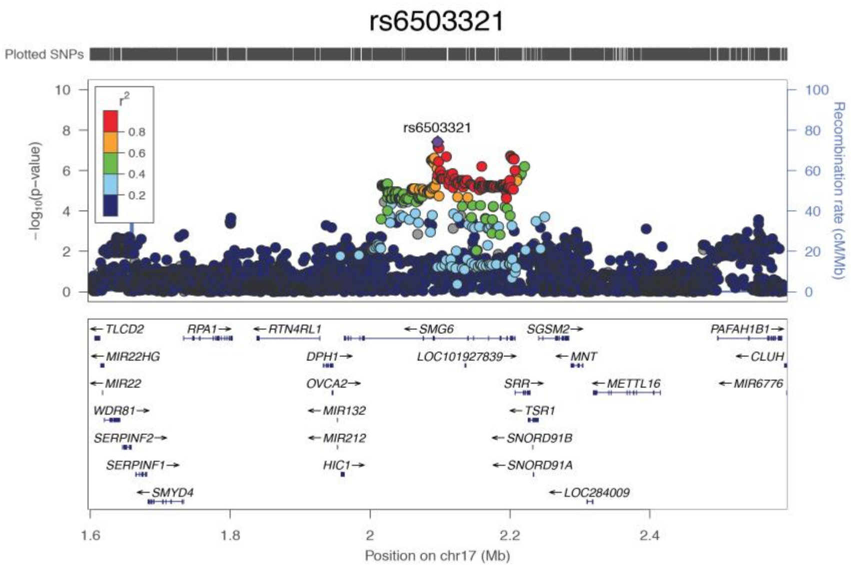
rs6503321
Plotted SNPs
rs6503321
−log10(p−value)
Recombination rate (cM/Mb)
r2
TLCD2
RPA1
RTN4RL1
SMG6
SGSM2
PAFAH1B1
MIR22HG
DPH1
LOC101927839
MNT
CLUH
MIR22
OVCA2
SRR
METTL16
MIR6776
WDR81
MIR132
TSR1
SERPINF2
MIR212
SNORD91B
SERPINF1
HIC1
SNORD91A
SMYD4
LOC284009
Position on chr17 (Mb)

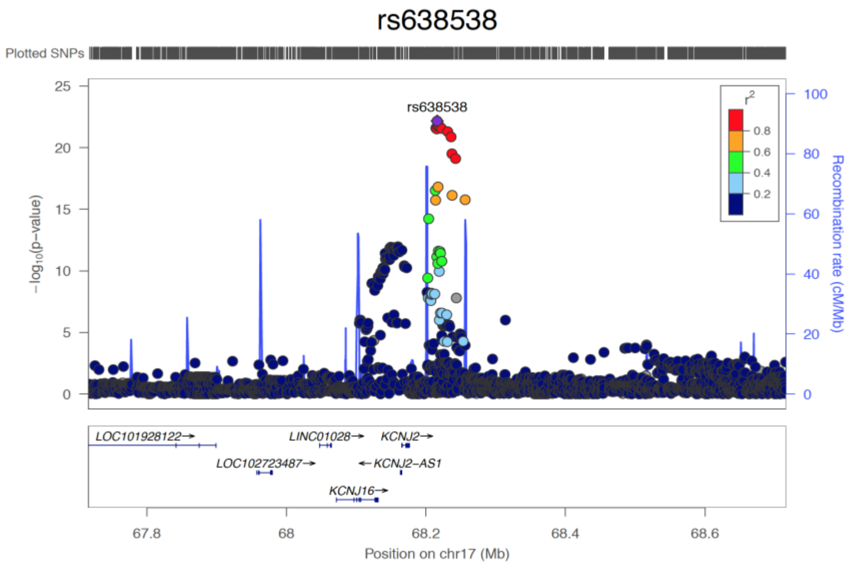
rs638538
Plotted SNPs
rs638538
−log10(p−value)
Recombination rate (cM/Mb)
r2
LOC101928122
LINC01028
KCNJ2
LOC102723487
KCNJ2-AS1
KCNJ16
Position on chr17 (Mb)

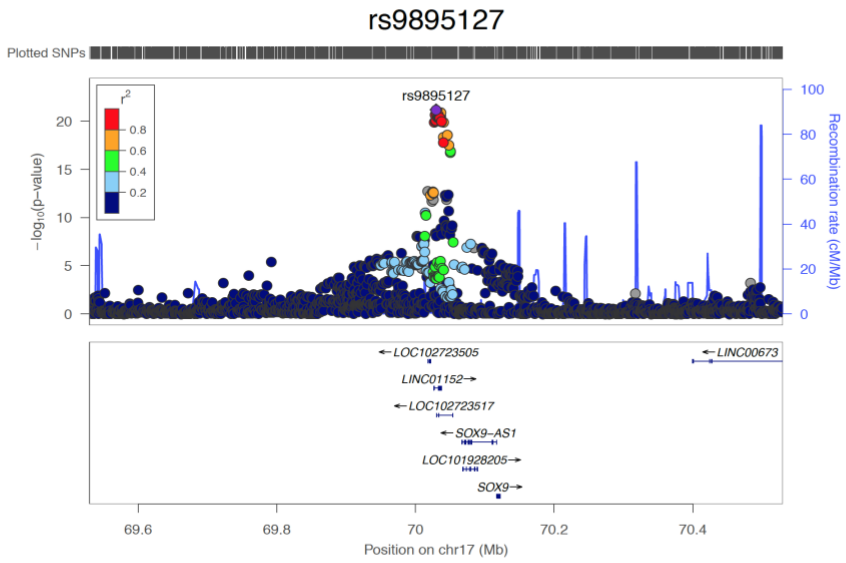
rs9895127
Plotted SNPs
rs9895127
−log10(p−value)
Recombination rate (cM/Mb)
r2
LOC102723505
LINC00673
LINC01152
LOC102723517
SOX9-AS1
LOC101928205
SOX9
Position on chr17 (Mb)

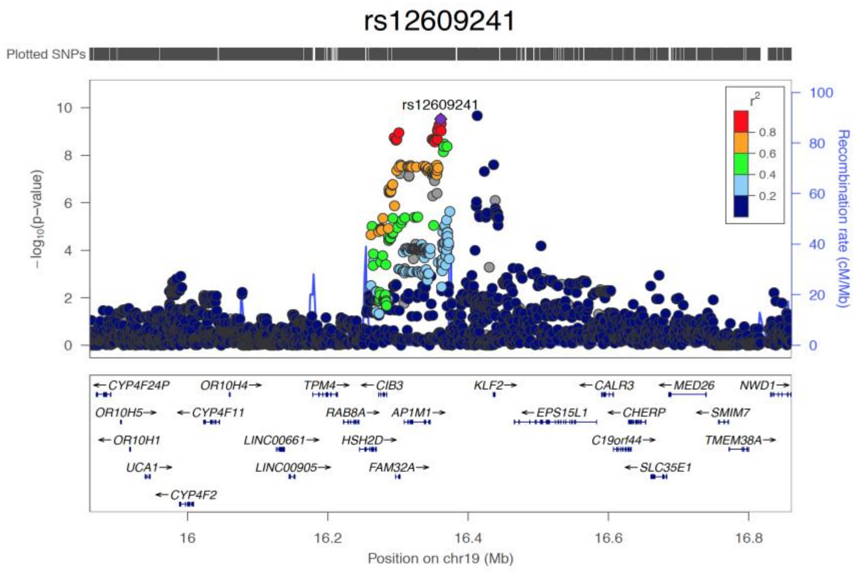
rs12609241
Plotted SNPs
rs12609241
−log10(p−value)
Recombination rate (cM/Mb)
r2
CYP4F24P
OR10H4
TPM4
CIB3
KLF2
CALR3
MED26
NWD1
OR10H5
CYP4F11
RAB8A
AP1M1
EPS15L1
CHERP
SMIM7
OR10H1
LINC00661
HSH2D
C19orf44
TMEM38A
UCA1
LINC00905
FAM32A
SLC35E1
CYP4F2
Position on chr19 (Mb)

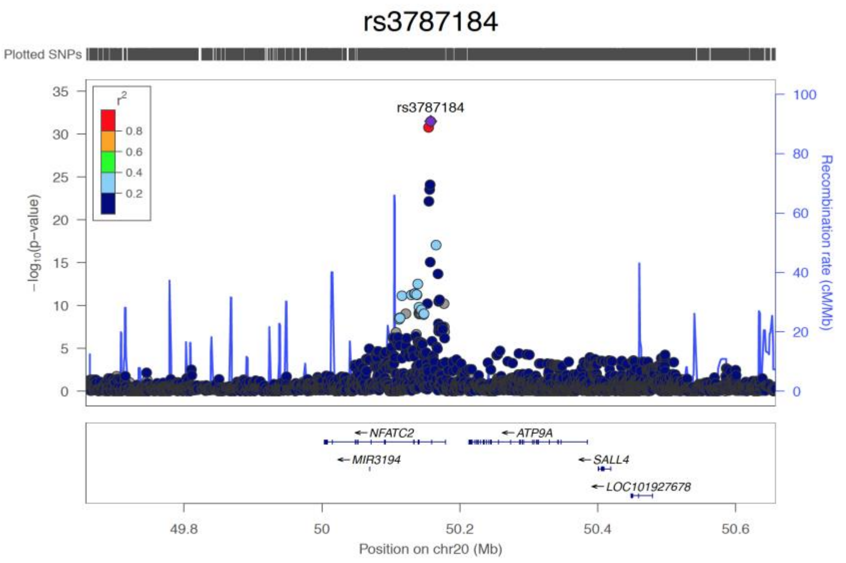
rs3787184
Plotted SNPs
rs3787184
r2
0.8
0.6
0.4
0.2
−log10(p−value)
Recombination rate (cM/Mb)
NFATC2
ATP9A
MIR3194
SALL4
LOC101927678
49.8
50
50.2
50.4
50.6
Position on chr20 (Mb)

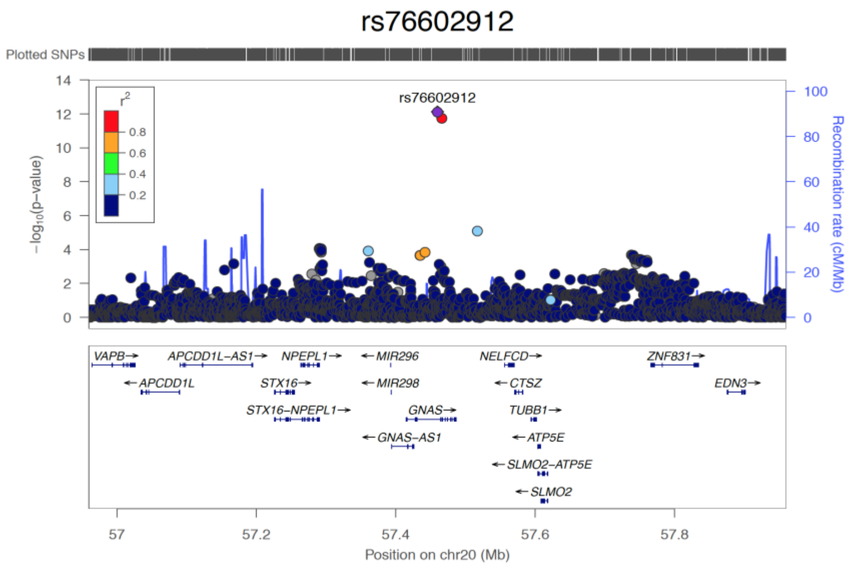
rs76602912
Plotted SNPs
rs76602912
r2
0.8
0.6
0.4
0.2
−log10(p−value)
Recombination rate (cM/Mb)
VAPB
APCDD1L−AS1
NPEPL1
MIR296
NELFCD
ZNF831
APCDD1L
STX16
MIR298
CTSZ
EDN3
STX16−NPEPL1
GNAS
TUBB1
GNAS−AS1
ATP5E
SLMO2−ATP5E
SLMO2
57
57.2
57.4
57.6
57.8
Position on chr20 (Mb)

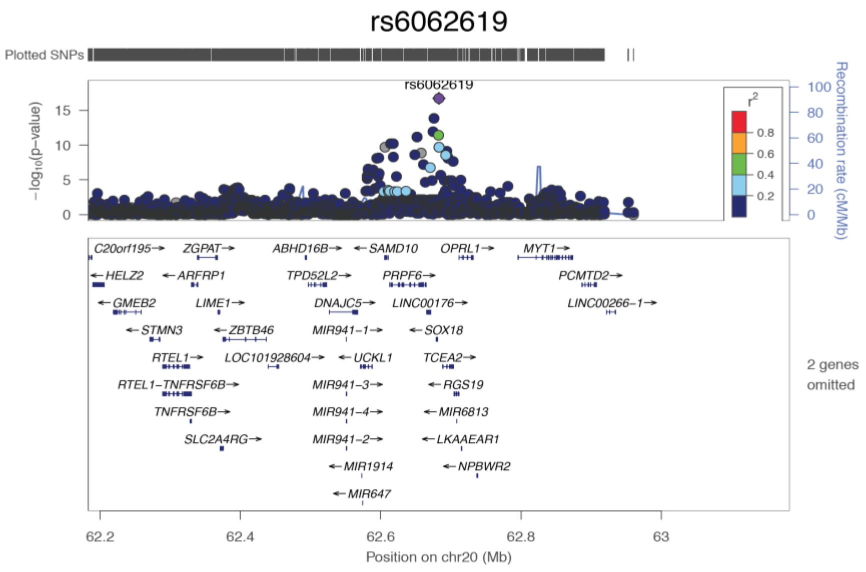
rs6062619
Plotted SNPs
rs6062619
r2
0.8
0.6
0.4
0.2
−log10(p−value)
Recombination rate (cM/Mb)
C20orf195
ZGPAT
ABHD16B
SAMD10
OPRL1
MYT1
HELZ2
ARFRP1
TPD52L2
PRPF6
PCMTD2
GMEB2
LIME1
DNAJC5
LINC00176
LINC00266−1
STMN3
ZBTB46
MIR941−1
SOX18
RTEL1
LOC101928604
UCKL1
TCEA2
RTEL1−TNFRSF6B
MIR941−3
RGS19
TNFRSF6B
MIR941−4
MIR6813
SLC2A4RG
MIR941−2
LKAAEAR1
MIR1914
NPBWR2
MIR647
2 genes omitted
62.2
62.4
62.6
62.8
63
Position on chr20 (Mb)

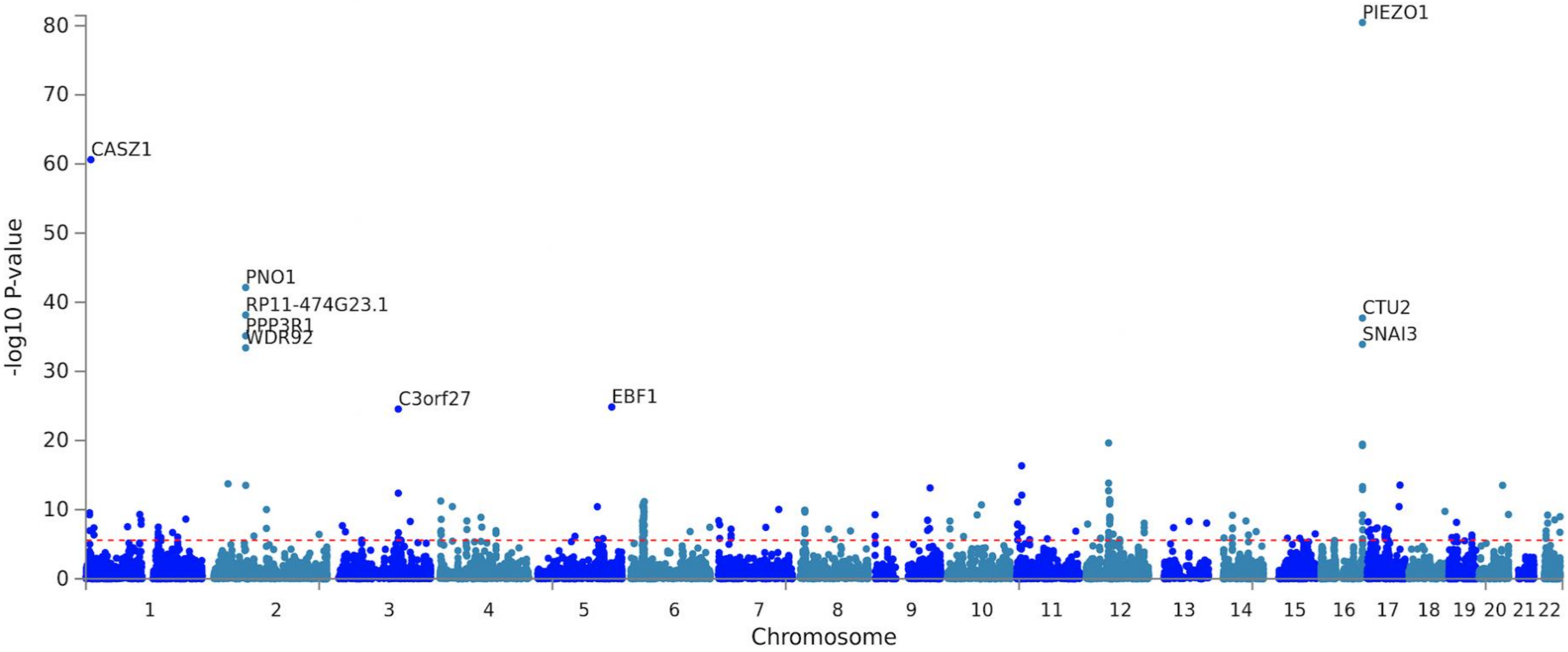


**Supplementary Figure 3. MAGMA Gene-based association analysis Manhattan plot.** Association results for varicose veins in MAGMA gene-based association analysis. The dotted red line indicates the threshold for genome-wide significance ($P < 2.68x10^{-6}$). 248 genes reached genome-wide significance in this analysis, with the top-ten genes highlighted in this figure.

#### 3. Supplementary Data

**Supplementary Data 1. Genome-wide significant variants at all significant discovery loci.** 12,391 genome-wide significant variants ($P < 5x10^{-8}$) at all 109 genomic risk loci identified in the discovery GWAS. Where the column headings pertain to: the SNP ID (RsID), chromosome, base position within NCBI Genome Build 37 (hg19), the effect allele, the alternate (non-effect) allele, the effect allele frequency in the study population, the imputation quality score; where 1 = genotyped SNP, the SNP effect size (BETA), the standard error of the BETA, and the GWAS association P-value. All variants are presented in ascending order according to chromosome and base position

**Supplementary Data 2. Genome-wide significant variants at the replicated susceptibility loci.** 5,315 genome-wide significant variants ($P < 5x10^{-8}$) identified by FUMA SNP2GENE at 45 of 46 replicated genomic risk associated with varicose veins. Where the column headings pertain to: the unique ID of the variant, the SNP ID (RsID), chromosome, base position within NCBI Genome Build 37 (hg19), the effect allele, the alternate (non-effect) allele, the effect allele frequency in the study population, the SNP effect size (BETA), the standard error of the BETA, the nearest gene according to positional mapping in FUMA, the functionality of the SNP as derived from ANNOVAR, the Combined Annotation Dependent Depletion (CADD) score of the variant, the RegulomeDB (RDB) score of the variant, the minimum chromatin state of the variant.

**Supplementary Data 3. MAGMA gene set analysis.** Gene sets were obtained from MsigDB v7.0 and total of 15,496 gene sets (Curated gene sets: 5,500, GO terms: 9,996) were tested.  Curated gene sets consists of 9 data resources including KEGG, Reactome and BioCarta (http://software.broadinstitute.org/gsea/msigdb/collection_details.jsp#C2 for details).  GO terms consists of three categories, biological processes (bp), cellular components (cc) and molecular functions (mf).  All parameters were set as default (competitive test). Gene sets are arranged in order of ascending P-value, with a P-value < 3.23 x10-6 indicative of a significant gene set, accounting for multiple testing (0.05/15,496).

**Supplementary Data 4. UK Biobank Summary Statistics**. The full summary statistics for the discovery GWAS of varicose veins in UK Biobank can be found on the Oxford Research Archive (ORA). URL: TBA
